# Registered Report: Replication and Extension of Nozaradan, Peretz, Missal and Mouraux (2011)

**DOI:** 10.1101/2025.03.13.643168

**Authors:** Karli M. Nave, Erin E. Hannon, Joel S. Snyder, Replication of Auditory Frequency Tagging Consortium

**Author notes:** **Address Correspondence to:** Karli Nave. **Contributing authors:** Emma Alexandrov, Sarah C. Allen, Fernando Barbosa, Mara Breen, Alexandre Celma-Miralles, Thibault Chain, Joshua R. de Leeuw, Fernando Ferreira-Santos, Ahren B. Fitzroy, Ningyao Geng, Sean A. Gilmore, Reyna L. Gordon, Jessica A. Grahn, Paz Har-Shai Yahav, Kiara Holm, Falk Huettig, Anne Keitel, Christian Keitel, Eniko Ladanyi, Yiguang Liu, Psyche Loui, Grace A. Lourie, Cyrille L. Magne, Inês Macedo, Cyrille L. Magne, Cecilie Møller, David Moreau, Srishti Nayak, Markus Ostarek, Rita Pasion, Mariana R. Pereira, Danna S. Pinto, Eva D. Poort, Meg Renzelman, Frank A. Russo, Jan Stupacher, Parker Tichko, Lucy Wight, Chu Yi Yu, Elana M. Zion Golumbic. **Protocol vetted by:** Daniel J. Simons. **Protocol edited by:** Karli M. Nave. **Multi-lab replication and extension of:** Nozaradan, S., Peretz, I., Missal, M., & Mouraux, A. (2011). Tagging the neuronal entrainment to beat and meter. The Journal of Neuroscience, 31(28), 10234-10240. **Registered protocols:** https://osf.io/rpvde/.

## Abstract

Cognitive neuroscience has long sought to disentangle stimulus-driven processing from conscious perceptual processing. Some prior evidence for neural processing of perceived musical beat (periodic pulse) may be confounded by stimulus-driven activity. Notably, Nozaradan et al. (2011) controlled for stimulus factors and used frequency tagging to show increased brain activity at imagery-related frequencies when listeners imagined a beat pattern during an isochronous stimulus. However, it remains unclear whether this effect is replicable and whether it reliably reflects conscious beat perception. This registered report presents 13 independent replications using the same vetted protocol. Listeners performed the same experimental paradigm as in Nozaradan et al. (2011), with an added behavioral task on each trial to assess conscious perception of the imagined beat. Pre-registered meta-analyses revealed smaller raw effect sizes of imagery condition (Binary: 0.03 µV, Ternary: 0.03 µV) than the original study (Binary: 0.12 µV, Ternary: 0.20 µV), with confidence intervals all overlapping with 0. Differences in full-sample estimated effect sizes (this study: n = 152, η_p_^2^ = .03–.04; 2011 study: n = 8, η_p_^2^ = .62– .76) suggest larger sample sizes are necessary to detect these effects reliably, if they exist. Additionally, only neural activity at the stimulus frequency predicted imagery task accuracy, contradicting our hypothesis that beat-related frequencies would predict performance. Our findings suggest an overall failure to replicate all main effects from the original study. We discuss potential reasons for discrepancies with the original study as well as implications for the utility of frequency tagging for studying beat perception.

## 1 Introduction

In cognitive neuroscience research, a pervasive problem is disentangling neural activity that reflects the processing of stimulus features from activity that reflects conscious experience and other high-level processes. Much research has been devoted to solving this problem in several areas of cognitive science, including visual mental representation (Kosslyn et al., 1978; Pylyshyn, 1981), speech perception (e.g., Dehaene-Lambertz et al., 2005) auditory spatial orienting (e.g., Rosen et al., 1999), visual and auditory bistable perception (e.g., Pressnitzer & Hupé, 2006), and musical rhythm perception (e.g., Iversen, Repp, & Patel, 2009). Musical *rhythm* refers to a pattern of temporal intervals arranged in a sequence, and musical *beat*^1^ refers to the quasi-isochronous pattern of prominent timepoints that often results from listening to a rhythm (Large & Palmer, 2002). Prior work provided evidence for a neural correlate of conscious beat perception, likely the result of activity in auditory cortical areas. Specifically, the study reported that human listeners had larger neural responses at beat related frequencies than at non-beat-related frequencies in complex rhythms (Nozaradan et al., 2016), which was interpreted as evidence for neural correlates of endogenous, top-down musical beat perception on the part of listeners. However, a later study presented the same rhythmic sequences to anesthetized animals and showed that stronger on-beat than off-beat responses could be recorded from the midbrain (Rajendran et al., 2017), raising the possibility that stimulus-driven exogenous processes, rather than top-down endogenous processes, might drive differential human brain responses to on-versus off-beat events.

Attempts to differentiate exogenous and endogenous processes in the brain are not unique to research on musical beat perception. A related controversy concerns whether mental *imagery* (process of accessing perceptual information from memory) relies on the same or distinct processes as stimulus processing, a debate addressed by studies of visual (Kosslyn et al., 1978, 1993; Pylyshyn, 1981, 2003) and auditory mental imagery (Halpern & Zatorre, 1999; Zatorre et al., 1996). The distinction between bottom-up, stimulus-driven and top-down, percept-driven processes is also evident in research on speech perception. For example, under certain conditions listeners can comprehend a linguistic message when presented with sine-wave speech, even though sine-wave speech has none of the acoustic attributes traditionally assumed to underlie speech perception (e.g., formants, formant transitions, fundamental frequency, etc.) (Remez et al., 1981). The amplitude, latency, and localization of brain responses are different when listeners hear the sine-wave speech as speech than when they hear it as non-speech, even when the same physical stimulus is presented across conditions (Dehaene-Lambertz et al., 2005). Such approaches are particularly promising for distinguishing between stimulus processing and conscious processing because they examine neural responses to the same physical stimulus across conditions in which the perception of that stimulus is altered, either by context or task goals.

Musical beat is an excellent candidate for examining the distinction between stimulus-driven and perception-related electrophysiological responses in the auditory central nervous system because its regularity is so prominent to the music listener, and it does not always depend on continuous physical input. For some musical rhythms, two or more different yet valid interpretations of the musical beat pattern can be heard, as with other auditory and visual bistable stimuli (Iversen et al., 2009; Kim & Blake, 2005; Pressnitzer & Hupé, 2006). This allows for more confident conclusions to be drawn about the higher-level processes involved in perception. Research on rhythm perception has indeed revealed brain activity that is to some degree isomorphic with the beat, as measured using both electroencephalography (EEG) and magnetoencephalography (MEG) (Fujioka et al., 2009; Nozaradan et al., 2011; Snyder & Large, 2005; Zanto et al., 2006). According to the Dynamic Attending Theory, listeners’ attention fluctuates at frequencies isomorphic with rhythmic external stimuli, causing the brain to form temporal expectancies about incoming auditory information (Jones & Boltz, 1989; Large & Jones, 1999). Similarly, Predictive Coding Theory posits that the human brain forms predictions based on the probability of a recurring pattern of events, such as musical beats in an auditory stimulus, specifically with the goal of minimizing prediction error (Vuust & Witek, 2014). Auditory rhythms are presumed to not only lead to entrainment at stimulus-related frequencies in the brain activity, but also to entrainment at perception-related frequencies in the brain. However, it is not entirely clear whether these previously discovered neural markers of auditory rhythm processing truly index perception of beat and meter, or whether they simply reflect stimulus processing that is propagated from low-level to high-level areas of the brain. Recent evidence suggests that that there may be significant contributions from low-level auditory brain areas that give rise to the perception of musical beat, including scalp-recorded activity that primarily arises from the brainstem (Tierney & Kraus, 2013) and more direct recording of action potential firing rates from midbrain neurons (Rajendran et al., 2017).

Although studies have attempted to disentangle lower-level processes from higher-level processes in musical beat perception, they often confound stimulus features with perception. For example, one study compared EEG responses while participants listened to a rhythm with a beat that was physically present or to a rhythm that required listeners to infer a beat that was not physically present in the stimulus (Nozaradan et al., 2018). Results showed that the latter beat frequency was evident in cortical brain activity but not in a measure reflecting brainstem activity (the *frequency-following response)*. While this finding was interpreted as supporting the claim that beat-related activity in the brain reflects conscious perception of the beat, physically distinct sound stimuli were used to generate these different brain responses. Several other studies also attempted to highlight differences between perception-related and stimulus-evoked neural processing of musical beat, yet they used different stimuli across conditions (e.g., Fujioka, Trainor, Large, & Ross, 2012; Nozaradan et al., 2016; Winkler, Háden, Ladinig, Sziller, & Honing, 2009).

At the time of pre-registration (2019), only one study had measured cortical responses at musical beat-related frequencies by manipulating the listener perception and holding the stimulus constant. Nozaradan et al. (2011) produced some of the first evidence that brain activity does not solely reflect the physical characteristics of the stimulus but rather listener perception. Listeners imagined a beat pattern (binary or ternary) while listening to a beat-ambiguous auditory stimulus (i.e., a metronome-like sequence of equal amplitude events that could be perceived as having either beat pattern). This study used frequency tagging by transforming the averaged event-related EEG responses from the time domain to the frequency domain and examined the SNR-corrected amplitude of brain activity at the beat frequencies. Higher amplitude neural activity was observed at frequencies corresponding to the imagined beat, as compared to non-beat frequencies. These brain responses that were isomorphic to the *imagined* beat frequencies showed an effect of the listener’s imposition of a specific structure onto the stimulus they heard.

This inspired other studies to use the frequency-tagging approach to examine music perception (Celma-Miralles et al., 2016; Chemin et al., 2014; Cirelli et al., 2016; Nozaradan et al., 2012; Stupacher et al., 2017; Tal et al., 2017a; Tierney & Kraus, 2015). Frequency-tagging has also been used to study perception in other areas of auditory research, such as word, phrase, and sentence level comprehension of speech (e.g., Ding, Melloni, Zhang, Tian, & Poeppel, 2016) and to study the effect of attention to targets during auditory scene analysis (e.g., Elhilali, Xiang, Shamma, & Simon, 2009). Earlier work used this technique in the visual domain to study visual bistable perception (e.g., Silberstein, 1995; Tononi, Srinivasan, Russell, & Edelman, 1998). Despite the breadth of research using this technique, there is little evidence, aside from the 2011 paper, indicating that frequency-related cortical brain responses reflect the listener’s perception of the beat in music, rather than low-level stimulus properties.

Furthermore, it is unclear to what extent this change in brain response at the beat frequency is influenced by other factors, such as the music training and dance training of the listener. Prior work has suggested that not only are high-level cortical auditory evoked potentials more reflective of beat and meter in rhythmically-trained individuals compared to non-rhythmically-trained individuals (Jongsma et al., 2004), but lower-level responses at the level of the brainstem are enhanced for musicians compared to non-musicians (Parbery-Clark et al., 2009). Musicians are also more sensitive to musical meter (hierarchical organization of strong and weak beats) than non-musicians (Palmer & Krumhansl, 1990). In addition, dancers exhibit enhanced processing of certain aspects of natural music, such as early neural responses to changes in the music when they are relevant to movement (Poikonen et al., 2016). To our knowledge, only one study using beat-related frequency-tagging has attempted to address the contribution of musical expertise. In this study, significant beat-related cortical activity was observed for two different types of stimuli: a rhythm with one clear beat pattern (quadruple meter) and a rhythm with a complex pattern comprised of two simultaneous beat patterns (quadruple and triple meters; i.e., a 4:3 polyrhythm) (Stupacher et al., 2017). Both musicians and non-musicians showed significant beat-related activity during the quadruple rhythm and the 4:3 polyrhythm. Interestingly, during a silent period following the rhythm 4:3 polyrhythm, only musicians showed significant beat-related activity, and this activity was correlated with their performance on a beat-matching task that followed the silent period. This suggests that music experience may indeed moderate the processes involved in auditory entrainment to musical rhythms. However, this study did not disentangle whether musical experience influences entrainment processes during auditory rhythm listening; both rhythms contained physical energy at the beat-frequencies themselves, making it impossible to dissociate stimulus-driven activity from perception-driven activity. To date, no one has investigated the extent to which music and dance training influence the perception-related activity demonstrated by Nozaradan et al. (2011). Moreover, given that nearly half of participants in the original 2011 study had extensive musical expertise (15-25 years of music training), it is particularly important to assess perception-driven brain responses in listeners with and without musical expertise.

In the current RR, we conducted a multi-lab study where we conducted a conceptual replication of the above paper, titled “Tagging the neural entrainment to beat and meter” (Nozaradan et al., 2011) and extended the findings to measure 1) the direct relation between the magnitude of the endogenous beat-related brain response and conscious perception of musical beat and 2) the relation between the magnitude of the beat-related brain response and music training. In this project, 14 labs completed independent, pre-registered studies, each using the same RR protocol. A minimum of six labs were required for the project.

The RR had three aims. The first aim was to estimate the true sizes of the effects reported in the original 2011 study by performing meta-analyses across research labs. The study was conducted using methods designed to be as closely matched to the original 2011 study as possible, allowing for an estimate of the meta-analytic replication of the original effects reported. All additional procedures were conducted such that they would not influence the replicated effects. Meta-analytic estimates examined the effect of condition (binary imagery, ternary imagery, or control task) on the SNR-corrected amplitude of electrical brain activity in the frequency spectrum at the binary frequency and the ternary frequency, as measured by EEG (outcomes that showed an effect in the original study).^2^ We hypothesized that the true effect size estimates would be similar to those reported by Nozaradan et al. (2011). The second aim of the RR was to improve upon the original methods by collecting a behavioral measure of beat perception on each trial, which allows a closer comparison between perception and brain activity and may additionally enhance listener attention and effort during the imagination task. We hypothesized that performance on the behavioral measure of conscious perception would be predicted by the magnitude of the beat-related brain response. The third and final aim of this RR was to extend our understanding of factors related to the findings of the original 2011 paper by measuring two hypothesized covariates: music training and dance training. We hypothesized that music and dance experience would be related to the magnitude of the beat-related brain response. By measuring years of training for both music and dance, we aimed to account for different types of interaction with music, which varies largely across cultures. The meta-analytic results of these analyses are presented in the results section.

The procedure followed in this RR was specific, unbiased, and transparent. We created a detailed study protocol, including explicit training instructions for the participant tasks, experimental materials, and a detailed experiment set-up guide. We designed a detailed analysis protocol, MATLAB script to process the EEG data, and R scripts to conduct the meta-analyses before viewing the data. Finally, the introduction, methods section, and results section (with placeholders for the final statistical results) were written prior to analyzing the data. All of these materials are publicly available on the Open Science Framework (OSF).

## 2 Disclosures

### 2.1 Preregistration

All preregistered materials are available at the project OSF page: https://osf.io/rpvde/.

### 2.2 Data, materials, and online resources

All materials necessary to conduct the study as well as summary data are available on the OSF project page: https://osf.io/rpvde. All data analyzed in this study are available on Harvard Dataverse (https://doi.org/10.7910/DVN/ELZFWK) (Nave, 2026).

### 2.3 Reporting

We report how we determined our sample size, all data exclusions, all manipulations, and all measures in the study.

### 2.4 Ethical Approval

All participating labs were required to collect their sample of data with the approval of an institutional review board in accordance with the Declaration of Helsinki and provide the protocol number.

## 3 Protocol Development and Requirements

Nave, Hannon, and Snyder proposed this RR project and developed the protocol. The first author of the original study provided the stimulus that they used. The protocol is available on the OSF project page for the RR project (https://osf.io/rpvde/).

*Advanced Methods and Practices in Psychological Science* publicly announced a call for laboratories interested in participating in this RR project on April 23, 2019. The data collection start date was May 1^st^, 2019, although labs were allowed to register to participate and begin data collection any time before the data collection end date. Data collection concluded on April 30^th^, 2022. Initially, we planned for the data collection period to last 12 months (ending April 30^th^, 2020), but due to the COVID-19 pandemic and local requirements to social distance, many labs were unable to resume EEG studies until as late as January 2022.

Prior to conducting the study, each lab submitted a pre-registered plan for implementing the approved protocol, and the first author reviewed each plan to ensure that it met the requirements of the protocol. Summaries of the labs’ pre-registered plans of protocol implementation can be found in Appendix A. Labs were required to note any deviations from the standard protocol, as well as any departures from their pre-registration that occurred during data collection (see Table S1).

## 4 Methods

### 4.1 Participants

Each lab committed to testing a minimum of sixteen^3^ healthy volunteers (after replacing participants who met the exclusion criteria below) between the ages of 18^4^ and 45, with normal hearing and no history of neurological or psychiatric disorders. While we hypothesized that the main analyses conducted in the original study (one-way ANOVAs) may have produced large effect sizes due to music and dance experience of the original sample, we expected that the true effect could be smaller in non-musicians and non-dancers. An a priori power analysis using G*Power 3.1 revealed that each lab would need 15 participants to detect a medium effect size (η^2^= .06) at .80 power (other parameters: alpha = .05, number of conditions = 3, number of repeated measures = 10, correlation among repeated measures = 0.5, non-sphericity correction = 1). Thus, we required 16 participants for a full lab sample to allow us to detect a medium to large effect size in order to capture a true effect that is smaller in size from the original study (η^2^ = .62, η^2^ = .76).

Of the 18 labs that applied to participate, 15 labs collected data for this study, and 14 labs completed the entire study protocol. Lab 14 was removed from any subsequent analyses after participant exclusions revealed a final sample that was too small (*n* = 4; minimum of eight required to be included in main analyses). After all exclusions, 13 labs were included in the pre-registered analyses, which included a total of 152 participants (see Table 1 for Descriptives).

**Table 1.** Descriptive results and general information for each of the 14 participating labs. For each participating lab, means and standard deviations (in parentheses) are provided for the sample demographics, the mean SNR-corrected amplitude of brain activity at the frequencies of interest (e.g., SSEPs), as well as mean performance on the behavioral measures. Values reported for Nozaradan et al. (2011) are medians, as reported in the original paper. ⁺ Lab 14 excluded due to n<8 included participants.

| Lab Num | Country | Test Lang | Total Tested | Total Included | Age | Music Years | Dance Years | Control Task |  |  |  |  | Binary Imagery |  |  |  |  |  | Ternary Imagery |  |  |  |  |  |
| --- | --- | --- | --- | --- | --- | --- | --- | --- | --- | --- | --- | --- | --- | --- | --- | --- | --- | --- | --- | --- | --- | --- | --- | --- |
|  |  |  |  |  |  |  |  | 0.8Hz | 1.2Hz | 1.6Hz | 2.4Hz | Task Acc | 0.8Hz | 1.2Hz | 1.6Hz | 2.4Hz | Task Acc | Conf Rating | 0.8Hz | 1.2Hz | 1.6Hz | 2.4Hz | Task Acc | Conf Rating |
| 0 | - | - | 8<br>(3f,5m) | 8<br>(3f,5m) | 30<br>(22-32) | - | - | 0.02 | 0.004 | 0.02 | 0.23 | - | -0.002 | 0.12 | -0.003 | 0.2 | - | - | 0.18 | 0.004 | 0.08 | 0.22 | - | - |
| 1 | Israel | Hebrew | 17<br>(11f,6m) | 11<br>(7f,4m) | 26.91<br>(22 - 43) | 0.64 (0-3)<br>0 yrs: n=8 | 2.18 (0-6)<br>0 yrs: n=5 | -0.025<br>(0.034) | -0.01<br>(0.029) | -86.76<br>(0.021) | 0.068<br>(0.044) | 0.977<br>(0.054) | -0.019<br>(0.032) | 0.025<br>(0.041) | -0.003<br>(0.013) | 0.074<br>(0.07) | 0.702<br>(0.175) | 4.391<br>(1.103) | 0.002<br>(0.031) | 0.006<br>(0.017) | 0.01<br>(0.016) | 0.071<br>(0.036) | 0.526<br>(0.118) | 4.482<br>(0.966) |
| 2 | USA | English | 14<br>(14f,0m) | 12<br>(12f,0m) | 18.83<br>(18-21) | 3.00 (0-10)<br>0 yrs: n=6 | 0.63 (0-4)<br>0 yrs: n=10 | -0.004<br>(0.05) | 0.005<br>(0.031) | -0.017<br>(0.024) | 0.066<br>(0.033) | 1.00<br>(0) | 0.049<br>(0.078) | 0.03<br>(0.05) | -0.015<br>(0.025) | 0.079<br>(0.046) | 0.658<br>(0.219) | 5.219<br>(1.065) | 0.016<br>(0.041) | 0.016<br>(0.086) | -39.91<br>(0.033) | 0.067<br>(0.053) | 0.625<br>(0.205) | 5.106<br>(0.966) |
| 3 | USA | English | 15<br>(7f,8m) | 11<br>(5f,6m) | 19.73<br>(18-22) | 6.32 (0-15)<br>0 yrs: n=3 | 0.36 (0-4)<br>0 yrs: n=10 | -0.095<br>(0.322) | -0.002<br>(0.04) | 0.052<br>(0.171) | 0.062<br>(0.037) | 0.97 (0.056) | -0.034<br>(0.093) | 0.036<br>(0.064) | -0.009<br>(0.051) | 0.059<br>(0.056) | 0.716<br>(0.238) | 5.564<br>(1.507) | -0.005<br>(0.054) | 0.008<br>(0.03) | -51.89<br>(0.024) | 0.071<br>(0.04) | 0.703<br>(0.242) | 5.482<br>(1.373) |
| 4 | United Kingdom | English | 17<br>(17f,0m) | 16<br>(16f,0m) | 21.86<br>(17-46) | 5.72 (0-15)<br>0 yrs: n=7 | 2.69 (0-16)<br>0 yrs: n=12 | 0.018<br>(0.024) | -0.002<br>(0.023) | -68.38<br>(0.024) | 0.035<br>(0.034) | 0.995<br>(0.021) | 0.014<br>(0.027) | 0.049<br>(0.066) | -0.019<br>(0.022) | 0.091<br>(0.08) | 0.762<br>(0.171) | 5.25 (0.903) | 0.059<br>(0.069) | 0.009<br>(0.033) | 0.023<br>(0.068) | 0.089<br>(0.115) | 0.713<br>(0.236) | 5.063<br>(0.701) |
| 5 | Canada | English | 11<br>(7f,4m) | 8<br>(5f,3m) | 23.75<br>(18-42) | 9.00 (0-37)<br>0 yrs: n=3 | 1.63 (0-6)<br>0 yrs: n=5 | -62.08<br>(0.037) | -0.007<br>(0.015) | -0.008<br>(0.014) | 0.059<br>(0.025) | 0.99 (0.029) | -0.008<br>(0.019) | 0.031<br>(0.029) | -0.004<br>(0.025) | 0.09<br>(0.038) | 0.626<br>(0.149) | 5.112<br>(0.476) | 0.018<br>(0.022) | -0.012<br>(0.015) | 0.004<br>(0.019) | 0.079<br>(0.042) | 0.679<br>(0.186) | 4.838<br>(0.605) |
| 6 | Netherlands | English | 14<br>(8f,6m) | 10<br>(5f,5m) | 25.00<br>(18-30) | 5.10 (0-20)<br>0 yrs: n=4 | 2.30 (0-12)<br>0 yrs: n=7 | 0.007<br>(0.036) | -0.012<br>(0.018) | -0.004<br>(0.013) | 0.067<br>(0.044) | 1.00<br>(0) | -0.007<br>(0.019) | 0.018<br>(0.02) | 0.003<br>(0.027) | 0.076<br>(0.06) | 0.56 (0.201) | 4.9 (0.97) | 0.012<br>(0.028) | 0.002<br>(0.033) | 0.009<br>(0.015) | 0.065<br>(0.036) | 0.63 (0.206) | 4.91 (0.923) |
| 7 | Portugal | Portuguese | 17<br>(13f,4m) | 9<br>(6f,3m) | 19.11<br>(18-22) | 3.83 (0-13)<br>0 yrs: n=5 | 2.78 (0-14)<br>0 yrs: n=6 | 0.002<br>(0.033) | 0.021<br>(0.035) | -0.008<br>(0.015) | 0.034<br>(0.037) | 0.963<br>(0.111) | -0.022<br>(0.041) | 0.009<br>(0.041) | 0.007<br>(0.025) | 0.055<br>(0.054) | 0.544<br>(0.224) | 4.244<br>(1.646) | 0.039<br>(0.041) | -0.004<br>(0.025) | 0.011<br>(0.024) | 0.075<br>(0.056) | 0.578<br>(0.233) | 4.233 (1.71) |
| 8 | Denmark | English | 16<br>(7f,9m) | 8<br>(3f,5m) | 24.00<br>(19-32) | 3.13 (0-9)<br>0 yrs: n=3 | 0.56 (0-3)<br>0 yrs: n=6 | -0.017<br>(0.031) | -0.009<br>(0.026) | -0.004<br>(0.019) | 0.041<br>(0.036) | 1.00<br>(0) | -0.023<br>(0.032) | -0.007<br>(0.027) | 0.001<br>(0.026) | 0.047<br>(0.045) | 0.55 (0.177) | 4.471<br>(0.923) | 0.024<br>(0.028) | 0.005<br>(0.028) | 0.001<br>(0.017) | 0.046<br>(0.036) | 0.618<br>(0.224) | 4.487<br>(0.964) |
| 9 | USA | English | 16<br>(10f,1m) | 13<br>(7f,5m) | 18.58<br>(18-22) | 2.54 (0-11)<br>0 yrs: n=8 | 0.85 (0-6)<br>0 yrs: n=11 | 0.045<br>(0.187) | -0.012<br>(0.035) | -0.006<br>(0.032) | 0.047<br>(0.073) | 0.936<br>(0.141) | -0.02<br>(0.094) | 0.053<br>(0.079) | -0.01<br>(0.065) | 0.098<br>(0.075) | 0.738 (0.18) | 5.577 (0.65) | 0.069<br>(0.075) | 0.015<br>(0.059) | 0.017<br>(0.06) | 0.089<br>(0.087) | 0.792<br>(0.233) | 5.708<br>(0.554) |
| 10 | New Zealand | English | 16<br>(8f,8m) | 11<br>(5f,6m) | 28.09<br>(18-41) | 4.00 (0-10)<br>0 yrs: n=4 | 1.09 (0-5)<br>0 yrs: n=7 | -0.004<br>(0.061) | 0.001<br>(0.041) | 0.006<br>(0.02) | 0.055<br>(0.038) | 0.992<br>(0.025) | -0.025<br>(0.042) | 0.09<br>(0.171) | 0.005<br>(0.021) | 0.107<br>(0.06) | 0.705<br>(0.205) | 4.964<br>(1.338) | 0.055<br>(0.081) | -0.03<br>(0.044) | 0.026<br>(0.071) | 0.115<br>(0.065) | 0.614<br>(0.247) | 4.691 (1.5) |
| 11 | Canada | English | 11<br>(5f,6m) | 10<br>(5f,5m) | 27.90<br>(20-45) | 5.60 (0-15)<br>0 yrs: n=5 | 0.30 (0-2)<br>0 yrs: n=8 | -0.028<br>(0.054) | -0.039<br>(0.046) | -0.035<br>(0.081) | 0.067<br>(0.078) | 0.992<br>(0.026) | 0.013<br>(0.086) | 0.054<br>(0.047) | -0.017<br>(0.018) | 0.118<br>(0.077) | 0.55 (0.276) | 5.16 (1.289) | 0.071<br>(0.116) | -0.003<br>(0.03) | 0.03<br>(0.092) | 0.102<br>(0.038) | 0.68 (0.22) | 5.17 (1.223) |
| 12 | USA | English | 23<br>(18f,5m) | 19<br>(15f,4m) | 22.00<br>(18-41) | 6.37 (0-35)<br>0 yrs: n=7 | 0.00 (0-0)<br>0 yrs: n=19 | -51.56<br>(0.055) | -0.003<br>(0.033) | -0.014<br>(0.04) | 0.085<br>(0.043) | 0.969<br>(0.041) | 0.008<br>(0.062) | 0.011<br>(0.029) | 0.006<br>(0.033) | 0.078<br>(0.041) | 0.632<br>(0.216) | 5.177<br>(0.939) | 0.011<br>(0.061) | 0.01<br>(0.026) | 0.005<br>(0.021) | 0.066<br>(0.038) | 0.581<br>(0.217) | 5.066<br>(1.005) |
| 13 | United Kingdom | English | 17<br>(13f,4m) | 14<br>(11f,3m) | 21.43<br>(19-29) | 7.00 (0-17)<br>0 yrs: n=5 | 3.25 (0-17)<br>0 yrs: n=8 | -0.042<br>(0.029) | -0.009<br>(0.044) | -0.008<br>(0.021) | 0.043<br>(0.045) | 0.97 (0.062) | -0.021<br>(0.059) | 0.043<br>(0.056) | -0.011<br>(0.026) | 0.093<br>(0.078) | 0.814<br>(0.146) | 5.114<br>(0.956) | -0.007<br>(0.073) | -0.014<br>(0.019) | 0.026<br>(0.041) | 0.071<br>(0.069) | 0.714<br>(0.207) | 5.121<br>(0.937) |
| 14 | USA | English | 8<br>(6f,2m) | 4*<br>(3f,1m) | 28.00<br>(19-33) | 6.38 (0-15)<br>0 yrs: n=1 | 5.00 (0-20)<br>0 yrs: n=3 | -0.07<br>(0.085) | 0.003<br>(0.024) | -0.009<br>(0.014) | 0.042<br>(0.024) | 1.00<br>(0) | -103.14<br>(0.032) | 0.045<br>(0.036) | -0.004<br>(0.021) | 0.044<br>(0.034) | 0.675<br>(0.359) | 6.075<br>(0.574) | 0.08<br>(0.079) | -0.009<br>(0.051) | 0.019<br>(0.054) | 0.051<br>(0.028) | 0.8 (0.082) | 5.425<br>(0.806) |

### 4.2 Stimuli

The stimulus, which was 33 s in duration, was obtained directly from the first author of Nozaradan et al. (2011). It consisted of a 333.3 Hz pure tone, which was amplitude-modulated with a 2.4 Hz periodicity, using an asymmetrical Hanning envelope (22 ms rise time and 394 ms fall time, amplitude modulation between 0 and 1) and then amplitude-modulated using an 11 Hz sinusoidal function, generating small irregularities resulting in a pseudo-periodic structure. To create a behavioral measure of conscious perception, we asked listeners to evaluate the fit of a probe that occurred after the initial 33 s stimulus. To do this, 3 s of the waveform were copied from the original and appended to the end of the stimulus and a probe was presented that occurred at a target or non-target position (relative to the imagery condition). The probe tone was added to the original signal and was presented at 880 Hz for 40 ms. The probe tone always occurred on either a binary beat (34.184 s after the beginning of the trial) or a ternary beat (33.732 s after the beginning of the trial) (See Figure 1). The auditory stimuli were presented binaurally through earphones or speakers at a comfortable hearing level (approximately 65 dB SPL). If speakers were used, the experimenter was required to listen to masking sounds during the test trials.

**Figure 1.**
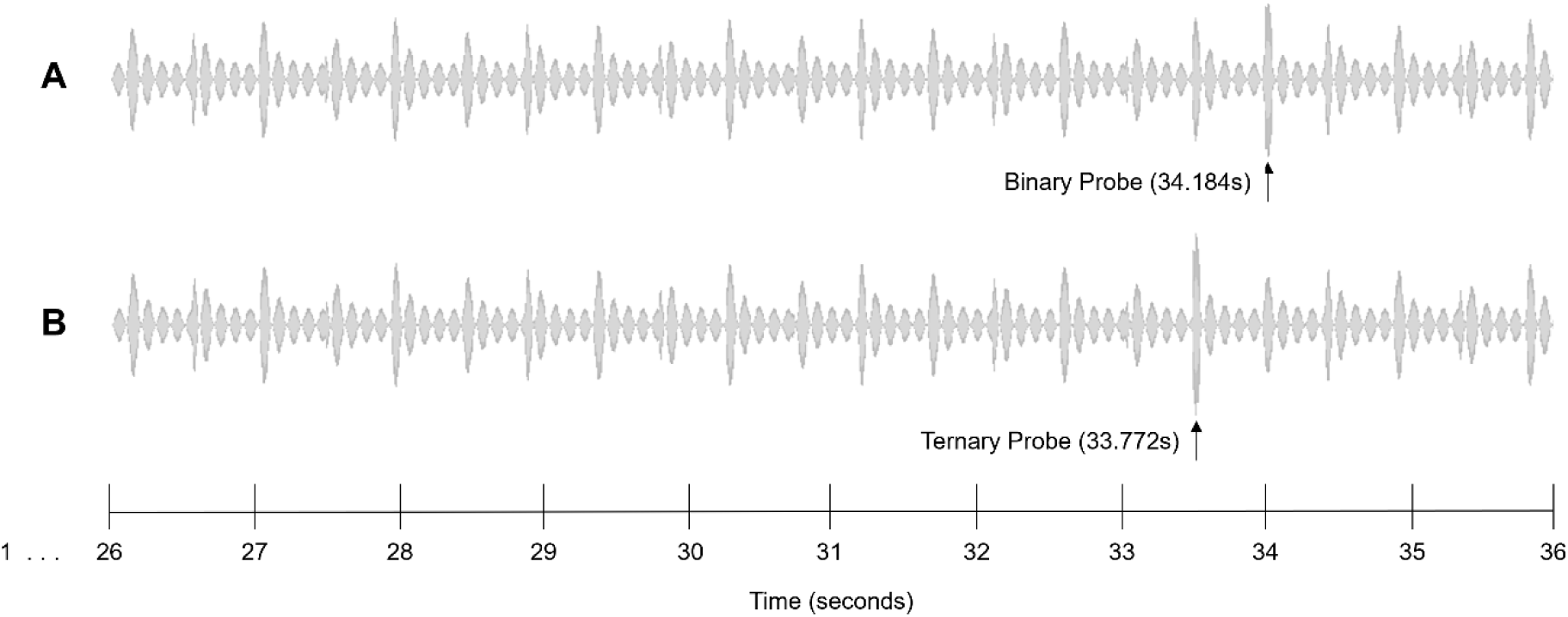
10 s excerpts of the 36 s auditory stimulus (x axis: time, y axis: sound amplitude). Note that this stimulus was extended by 3 s from the original stimulus used by Nozaradan et al. (2011), in order to allow imagery to be maintained for the same amount of time as the original study before introducing the probe tones. The probe tone (880 Hz) was superimposed onto the stimulus for 40ms. A) Binary probe tone, superimposed starting at 34.184 s. B) Ternary probe tone, superimposed starting at 33.772 s.

### 4.3 Task

Participants performed the same three conditions that were performed in the original study: a control condition; a binary imagery condition; and a ternary imagery condition. Each condition was presented as a separate block of trials: 12 trials (control condition) or 10 trials (imagery conditions) during which the auditory stimulus was presented after a 3 s silence. Stimulus presentation was self-paced. First, participants completed the block of control trials, during which they detected a very short (4 ms) sound interruption, which was inserted in two of the trials. The two trials containing a short interruption were excluded from analyses.

During the second and third blocks, participants completed the binary imagery condition and ternary imagery condition, counterbalanced for order. They imagined a binary beat structure or a ternary beat structure in the stimulus. Before participants completed each of the binary and ternary imagery conditions, they completed a standardized training procedure with an experimenter. All labs were required to send a video of a training session with one pilot participant for all three tasks, which was reviewed and approved by the first author prior to starting data collection (see OSF project page for example video made by first author). During training, participants were instructed to begin their imagery as soon as the stimulus began, making the first sound they heard a strong beat, and to maintain this imagery as consistently as possible throughout the entire trial. To help them understand exactly how to perform the imagery conditions, participants were asked to perform overt movements (hand tapping) and to count aloud, first with the help of the experimenter and then alone. Participants then completed a practice trial with just imagery, in which they practiced not moving or counting aloud while maintaining the imagery. Once they were comfortable with the imagery task, participants practiced maintaining the imagery and judging whether a probe tone occurring near the end of the stimulus was on the beat. All aforementioned training trials could be repeated until the participant felt comfortable to move on. Finally, participants completed two practice trials that included a response to the probe (*probe task*) and a rating to indicate their success at maintaining the imagery throughout the entire trial (*imagery success rating*). For the probe task, participants reported whether the probe tone occurred on a strong beat or a weak beat. For the imagery success task, they indicated how well they maintained the imagery throughout the entire trial using a 7-point rating scale, in which 1 was “Completely Got Lost” and 7 was “Maintained the Entire Time” ^5^. Participants had to respond correctly on a group of two probe trials (one “correct”, one “incorrect”) during training prior to advancing to the test trials. If the participant did not get both training trials correct, the trials were re-administered to the participant again in a randomized order. If the participant did not respond correctly to both training trials after four repetitions (a total of eight practice trials), the experiment moved on to the test trials. However, this participant was then excluded from the main analyses. Each participant completed 10 test trials during each imagery condition. Participating labs were provided with a detailed script for training. Finally, participants filled out a demographic questionnaire, including questions regarding music and dance experience. The experiment, not including EEG set-up time, took approximately 30 to 45 minutes, depending on time required to complete training with each participant.

### 4.4 Experiment Script

Participating laboratories were provided with a Presentation (Neurobehavioral Systems, Inc.) script that ran the study and collected all behavioral responses. In the case that a laboratory did not have a Presentation software license, they were instructed to use the RR project’s specific instruction manual on how to set the experiment up with their own software, and they were required to upload their script to the OSF website. If necessary, labs were permitted to adapt the experiment instructions to a language other than English. These labs were required to upload the translated materials as well as a translator statement. See Table S1 for specific details for each lab.

### 4.5 EEG Recording

Each lab used their own EEG systems to collect data while participants completed the three conditions described above. While participants were requested to refrain from movement during the experiment, it was still possible that participants would engage in small micro-movements while listening to the rhythmic auditory stimuli. Thus, participating labs had the option to contribute additional data by measuring overt rhythmic movements by placing two external electrodes, one over the Sternocleidomastoid Muscle (SCM) of the neck and one on the First Dorsal Interosseous (FDI) muscle of the dominant hand. Recording from the neck and hand was not required for participation, in case labs were unable to record movement activity but still wished to participate. Details about each lab’s EEG equipment, number of electrodes, and electrode placement are detailed in the Individual Lab Details (see Table S1). Following suggestions from the authors of the original study, we provided careful instruction and motivation to complete the task prior to each test block. Experimenters were asked to monitor compliance with the instructions and help motivate the participant to perform the imagery conditions to the best of their ability, either by staying in the room with the participant during the test trials or by monitoring a live video feed (see Appendix A for more details), and they recorded any observable movement in the session notes.

## 5 Data Analysis

The data analyses were conducted by the first author at University of Western Ontario in accordance with the preregistered analysis plan, available at the project OSF Website (https://osf.io/4xqwu/). All participating labs sent their raw EEG data with trial event codes indicating the beginning of trial times in each of the experimental blocks in one of the approved formats^6^. Participating labs were required to send a detailed description of how they generated their event code list, including how they accounted for latency issues (if any). Labs were also required to send their log of participants’ movement, behavioral data, and survey data in an excel workbook (see individual lab data summaries on OSF page). The core meta-analyses were based on SNR-corrected amplitudes at the imagery-related frequencies (binary: 1.2 Hz, ternary: 0.8 Hz) for the three conditions (control task, binary imagery, ternary imagery). This RR reports the average SNR-corrected amplitudes (SSEPs), behavioral responses, and effect sizes for each lab. The official EEG data processing steps were written without viewing the actual data and have been made publicly available.

### 5.1 Stopping Criteria and Exclusions

As part of registering to participate in this RR and prior to beginning data collection, each lab indicated whether it aimed to stop data collection after collecting a full sample of 16 participants or a half sample of eight participants. The stopping criteria were designed to ensure that each lab would meet the minimum data collection requirements for the protocol and that the decision to end data collection would not be influenced by the results of the study.

Data were pre-registered to be excluded if a participant did not fit the specified recruitment criteria, did not follow instructions, did not complete the experiment, or if the experiment was administered incorrectly for any reason (e.g., incorrect training procedures). Labs were asked to note this explicitly at the time of the experiment in a lab testing log before examining the data for that participant. Participants were also excluded if they were observed moving during EEG data collection during the auditory stimulus in any way that may have affected imagery (i.e., moving rhythmically, tapping their fingers, hands, or feet, bobbing their head). Experimenters were asked to make note of whether each participant was observed moving in a periodic manner. Fifty-nine of the 212 total participants were excluded: 42 for failing imagery training, eight due to excessive EEG artifacts (present in > 50% of EEG data), four due to experimenter or equipment error, one for observed rhythmic movement, and four due to the lab having too few participants after exclusions (Lab 14).

### 5.2 EEG Analysis

All EEG analyses were performed identically to the original paper, using MATLAB (The MathWorks), EEGLAB toolbox (Delorme & Makeig, 2004), and FieldTrip (Oostenveld et al., 2010). Prior to collecting the full dataset, all labs were required to conduct the full study with one pilot participant and send the data to the parent lab. These data files were used to create the final data processing scripts, which followed the analysis plan posted to the project OSF page during pre-registration.

All participating labs were asked to record the EEG signals with a low-pass filter of 500 Hz and a sampling rate of at least 1000 Hz^7^. Actual sampling rates for each lab are included in Table S1. In order to minimize the amount of resampling conducted on the data, labs reported their EEG system requirements and limitations prior to data collection, and the default sampling frequency [1000 Hz or 1024 Hz] was set to the sampling frequency most common among the EEG systems being utilized as of the data collection start date. Where necessary, data were downsampled to 1000 Hz or 1024 Hz (see Table S1 for individual lab details). Prior to completing any further processing steps, the first author inspected the raw data for all participants to ensure no excessive noise or artifacts were present. At this point, eight participants were excluded due to an excessive percentage of data including artifacts (> 50% of the data). Data were referenced to the average of all EEG electrodes and filtered using a 0.1 Hz high-pass Butterworth zero-phase filter to remove very slow drift in the recorded signals. EEG epochs began 1 s after the onset of the stimulus and lasted 32 s. Artifacts produced by eye blinks/saccades and muscles were removed using independent component analysis (ICA) (Jung et al., 2000), using the runica algorithm (Makeig et al., 1996). Selections of ICA components for removal were conducted by two independent researchers (K. Nave and T. Chabin) and compared for reliability (see the pre-registered analysis plan on the OSF page for more details). During ICA manual selection, researchers were blind to participant and condition. After ICA rejection, for each participant and condition, averages were conducted across trials and transformed using a discrete Fourier transform (Frigo & Johnson, 1998) with a frequency resolution of 0.03 Hz, producing a frequency spectrum ranging from 0 to 500 Hz. The contribution of residual noise was removed by subtracting the average amplitude measured at neighboring frequency bins (two frequency bins ranging -0.15 to -0.09 Hz and from +0.09 to +0.15 Hz relative to each bin) at each bin of the frequency spectra. Stimulus- and imagery-related frequency responses were estimated by averaging the signal SNR-corrected amplitude measured at the three frequency bins centered on the target frequency. The magnitude of these responses was averaged for each participant, condition, and target frequency across all scalp electrodes. Recordings from overt movements made by the neck and hand are reported and analyzed in the Exploratory Analysis section.

### 5.3 One-Way ANOVAs and Post-Hoc Tests on Effect of Condition

As in the original study, we conducted repeated-measures one-way ANOVAs to test the effect of condition (control, binary imagery, ternary imagery) on the SNR-corrected amplitude of brain activity at the stimulus frequency (2.4 Hz), each of the imagery-related frequencies (binary: 1.2 Hz, ternary: 0.8 Hz), and the first upper harmonic of the ternary frequency (1.6 Hz), separately for each lab and across the entire collected sample. To compare the effect of condition for the four frequencies of interest (ternary (0.8Hz), binary (1.2Hz), ternary harmonic (1.6Hz), and stimulus (2.4Hz)) with the original study, the results with estimated effect sizes are provided for each participating lab, across all labs combined, and for Nozaradan et al. (2011) in Table 2.

**Table 2.**
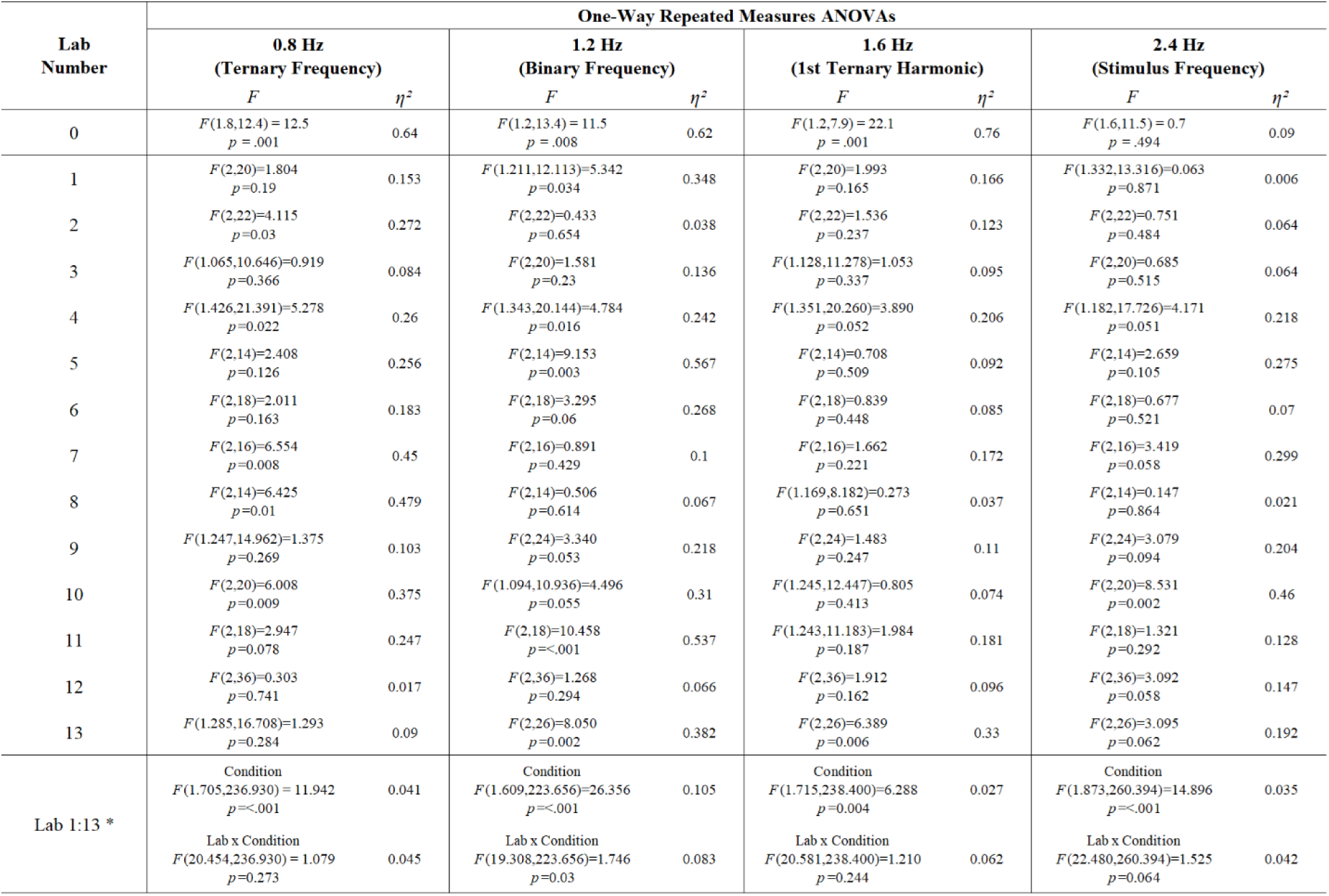
Results of Repeated Measures ANOVA for the original study and participating labs. For the Repeated Measures ANOVA applied to each lab, the main effect of Condition is reported. For the Repeated Measures ANOVA involving all 13 labs (last row, *N* = 152), the main effect of Condition and the interaction Lab x Condition are reported. Lab 0 = Nozaradan et al. (2011). *This analysis was not pre-registered and was added post-data collection.

The original paper also reported post-hoc tests, which revealed that both imagery-related frequencies only demonstrated higher SNR-corrected amplitudes of the brain response when the imagery frequency matched the imagery performed (e.g., the binary frequency was enhanced when participants performed the binary imagery, but not when they performed the ternary imagery or during the control condition). To compare the effect of condition for the three frequencies of interest (binary, ternary, and stimulus) with the original study, these are provided for each participating lab in Table 3.

**Table 3.**
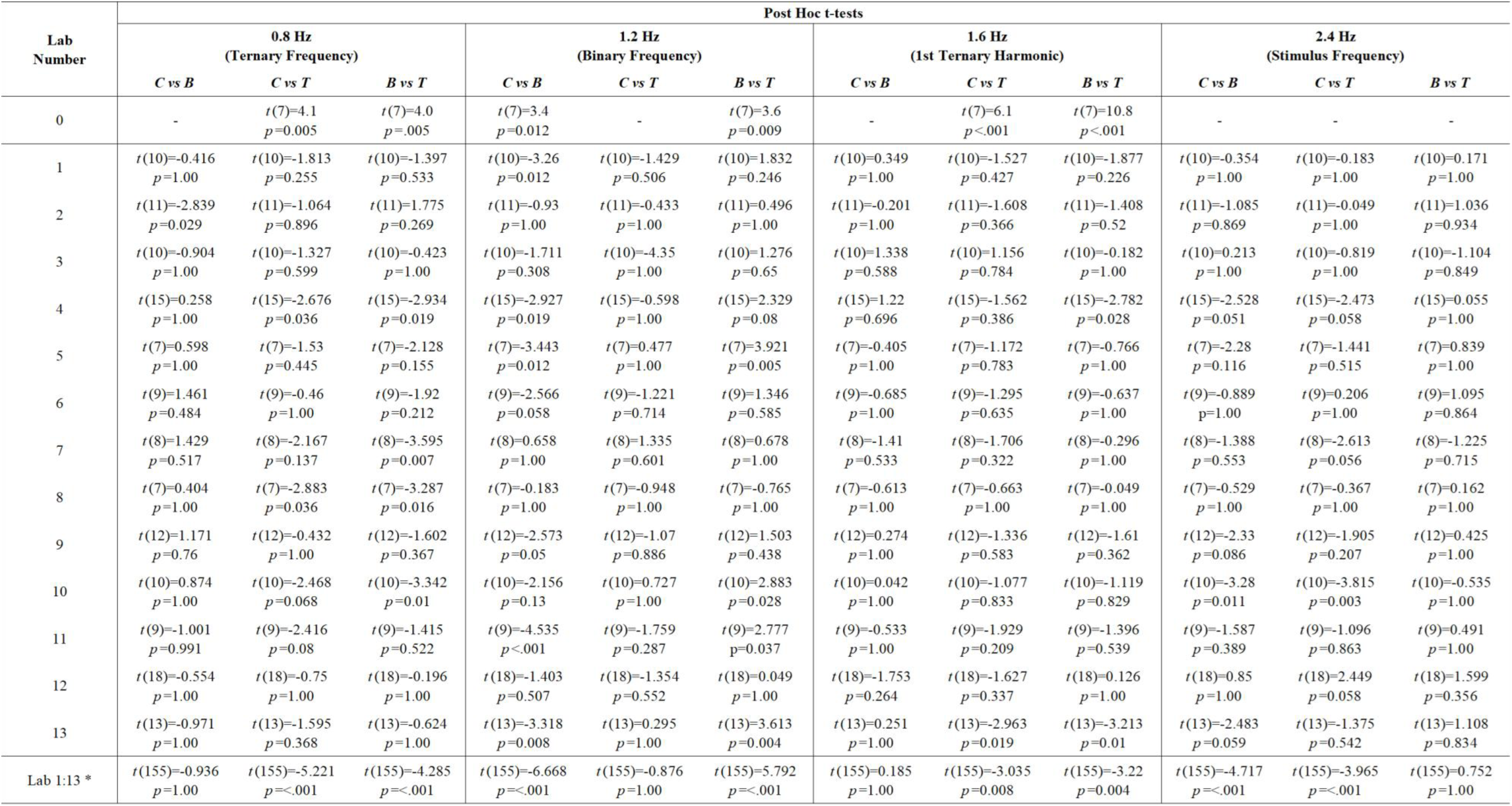
Results of post-hoc t-tests for the original study and participating labs. Final row indicates results when collapsing across all 13 participating labs (*N* = 152). Lab 0 = Nozaradan et al. (2011). *This analysis was not pre-registered and was added post-data collection.

### 5.4 Meta-Analytic Estimates on the Effect of Imagery

The intended analyses were pre-registered and tested on simulated data before inspecting the data. The RR measured the meta-analytic across-lab effects of the raw mean differences between conditions in frequency-domain SNR-corrected amplitudes of the brain responses, where the conditions compared were: the Control (C) task (e.g., in which no imagery took place), the Binary Imagery (B) task (e.g., in which beats were imagined on every other stimulus event), and the Ternary Imagery (T) task (e.g., in which beats were imagined on every third stimulus event). The original 2011 paper reported a statistically significant effect of condition (3-way Univariate ANOVA) on the SNR-corrected amplitude at imagery frequencies based on the beat pattern that the participant was asked to imagine, as well as post hoc significant comparisons between the conditions as expected (0.8 Hz: T > C, T > B; 1.2Hz: B > C, B > T). In order to examine these condition comparisons, we used random effects meta-analyses to estimate four raw effect sizes, calculated as the difference in mean SNR-corrected amplitude of brain activity between two conditions at each of the two beat imagery frequencies (i.e., ternary: 0.8 Hz, binary: 1.2 Hz): 1) Binary Control Effect: binary imagery minus control task at the binary frequency (1.2Hz), 2) Binary Active Effect: binary imagery minus ternary imagery at the binary frequency (1.2Hz), 3) Ternary Control Effect: ternary imagery minus control task at the ternary frequency (0.8Hz), and 4) Ternary Active Effect: ternary imagery minus binary imagery at the ternary frequency (0.8Hz). In all four cases, a significant positive estimate would provide evidence for the conclusions of the original study, suggesting a positive modulation of the SSEP amplitude for the imagery-related frequency. In addition, the RR implemented mixed-effects meta-analyses to test music experience and dance experience as moderating variables. The meta-analytic estimates are provided both with and without the moderators included. These random- and mixed-effects meta-analyses were conducted using the R package metaphor (Viechtbauer, 2010).

### 5.5 Logistic Regression to Predict Beat-Related Task Performance

If the effects observed in the original study indeed reflect the imagined beat pattern, then it would be expected that accuracy on the probe task would be related to the SNR-corrected amplitude of brain activity at imagined beat frequencies. In addition, prior work suggests that musicians and dancers may have enhanced sensitivity to rhythmic organization (Palmer & Krumhansl, 1990), as well as enhanced beat-related brain responses (Jongsma et al., 2004). To test the contribution of these factors to the size of the beat-related brain response, a generalized estimating equation (GEE) binary logistic regression was conducted to examine whether task condition predicted response accuracy while accounting for repeated observations within participants. The dependent variable was accuracy (0 = incorrect, 1 = correct). Categorical predictors included imagery type (binary or ternary) and trial type (ON beat vs OFF beat), and continuous covariates included years of music training, years of dance training, SNR-corrected amplitude at the imagined beat frequency (Beat Imagery SSEP), and SNR-corrected amplitude at the stimulus frequency (Stimulus SSEP). A binomial distribution with a logit link function was specified. Participant-level clustering was modeled by nesting trials within participant ID, who were further were nested within lab ID (LabID*PartID). An autoregressive (AR(1)) working correlation structure was used, and significance of predictors was assessed using Wald chi-square tests. Model convergence criteria followed Fisher scoring with a maximum of 1000 iterations and absolute convergence tolerance of .0001.

### 5.6 One-Tailed T-Tests comparing SSEPs to Noise Floor

Regardless of effects of imagined frequency, we expected every lab to observe robust neural activity at the stimulus frequency (2.4Hz, the frequency of the periodic physical stimulus). As such, activity at the stimulus frequency can be considered as a type of ‘positive control’, demonstrating that the experimental setup across labs was working as intended and regardless of imagery condition. To this end, we compared SNR-corrected amplitude to zero for each frequency (binary, ternary, and stimulus). All labs demonstrated average SSEPs at 2.4Hz that were statistically greater than 0 for all conditions, shown in Table S2.

## 6 Results

The goals of this RR were to 1) provide a precise measure of the size of the effects of beat imagery, and 2) extend the findings by relating the magnitude of the effects to a) a behavioral measure of beat perception and b) music/dance training of the participants – by combining the results from multiple, independently conducted studies. The results of all contributed studies are included in the analyses reported here regardless of their outcome, providing an unbiased meta-analysis of the effects. The analysis does not focus on null-hypothesis significance testing, but rather on the meta-analytic effect size for each outcome, with the confidence intervals estimated.

### 6.1 Descriptive Statistics

Descriptive statistics for each contributed study are provided in Table 1. Probe task accuracy was computed as the average correct responses across test trials, where a binary probe tone was indicated as “ON beat” during binary imagery only, and a ternary probe tone was indicated as “ON beat” during ternary imagery. Imagery success rating was computed as the average imagery success rating given (1 = “Completely got lost”, 7 = “maintained imagery the entire time”) across test trials. In addition, the average SNR-corrected amplitude of brain activity at the stimulus frequency, binary frequency, and ternary frequency is provided for each condition, along with average performance on all behavioral tasks. Results from the probe task measures are presented in Figure 2. Average frequency spectra for each lab and across all labs are displayed in Figure 3. All labs demonstrated average SSEPs at the auditory stimulus frequency (2.4Hz) that were statistically greater than 0 for all conditions (see Table S2), confirming the experimental setup was adequate to capture significant auditory SSEPs.

**Figure 2.**
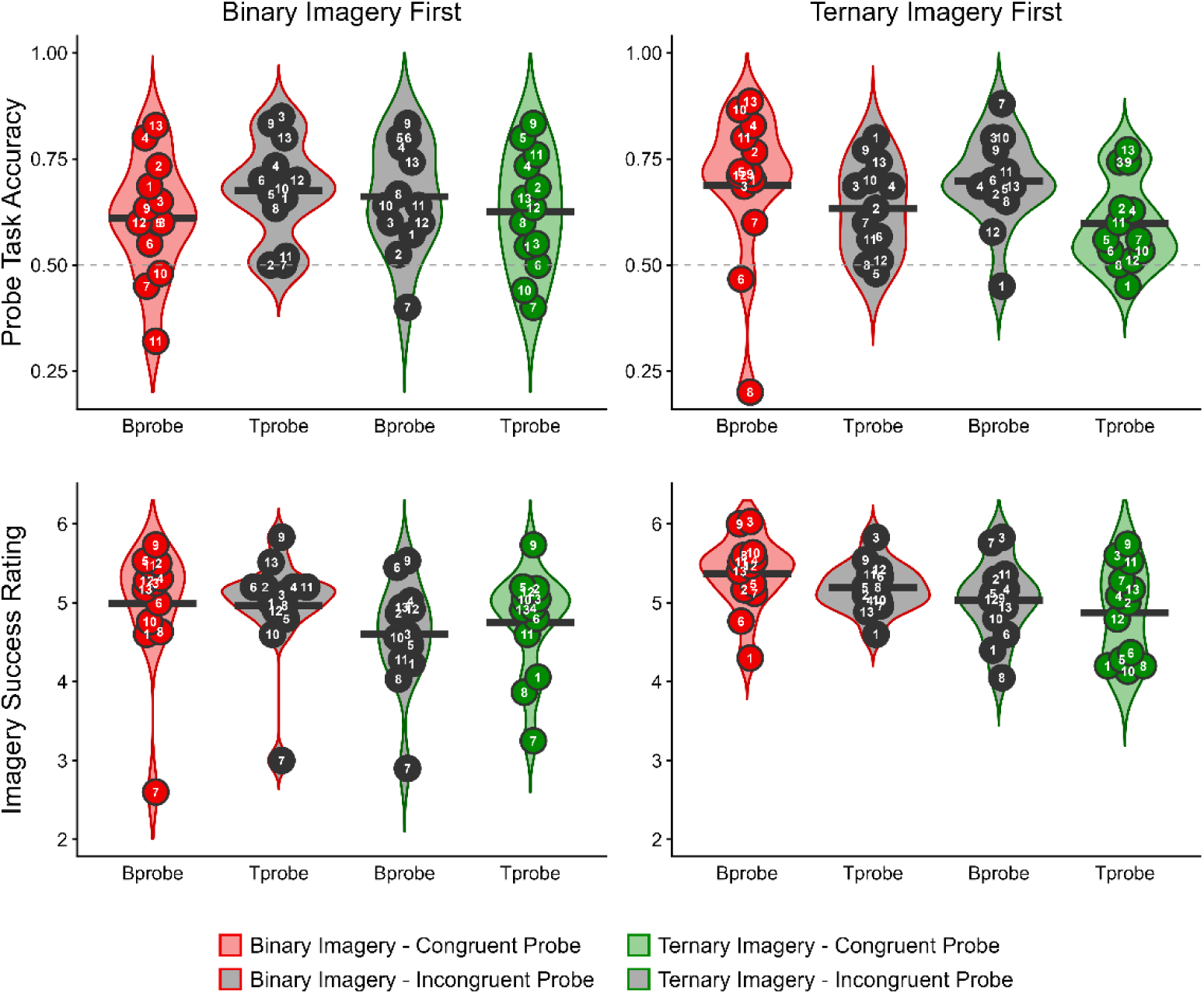
Average probe task accuracy (**top**) and average imagery success rating (**bottom**) across participants from all participating labs on the behavioral task during imagery conditions. Binary imagery trials (red) and ternary imagery trials (green) are shown separately based on probe type (Bprobe: congruent with binary beat / ON beat for binary imagery, Tprobe: congruent with ternary beat / ON beat for ternary imagery). Congruent trial conditions (e.g. ON beat trials) are color-shaded and incongruent trials (e.g. OFF beat trials) are shaded in grey. Data are visualized separately based on whether the participants completed the binary imagery block first (left column) or the ternary imagery block first (right column). Each numbered dot represents the mean of one of the participating labs, labeled with the lab number. Solid lines represent the mean across labs (N = 13 labs, *N* = 152 participants).

**Figure 3.**
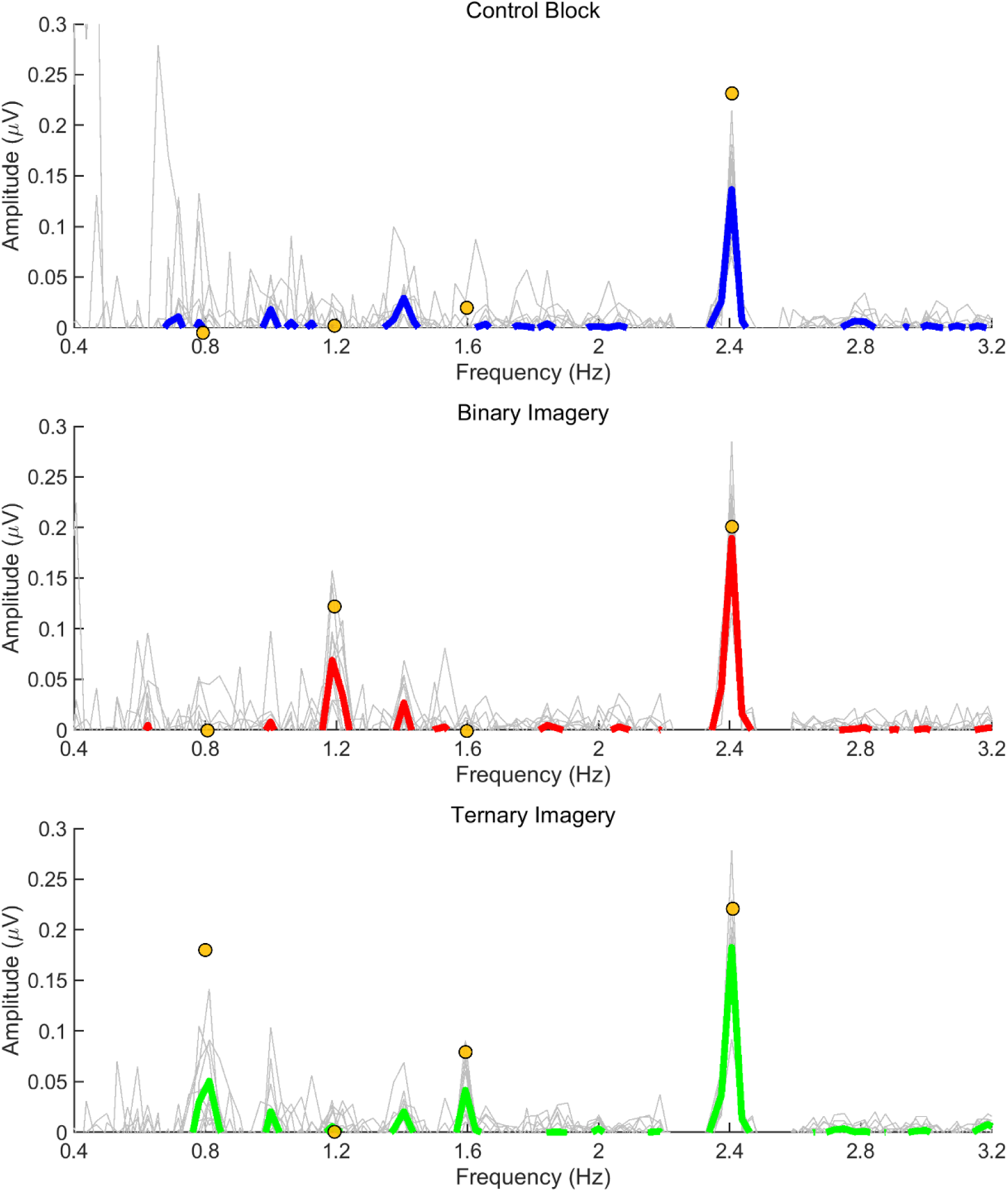
Lab-level average of the steady state-evoked potentials (SSEPs) elicited by the 2.4 Hz auditory stimulus in the control condition (top, blue), the 2.4 Hz stimulus plus binary imagery condition (middle, red), and the 2.4 Hz stimulus plus ternary imagery condition (bottom, green).The lab-level average frequency spectra are shown using a thick colored line, while single-lab averaged spectra are shown in thin gray lines. The frequency spectra represent the SNR-corrected amplitude of the EEG signal (in microvolts) as a function of frequency, averaged across all scalp electrodes, after applying the noise subtraction procedure. Yellow dots represent the reported median values from the original study (Nozaradan et al., 2011).

#### 6.2.1 Effect of Condition on Brain Activity at Binary Frequency (1.2 Hz)

##### Binary Control Effect

Figure 4 shows the point-estimate for the meta-analytic effect of condition on the binary frequency (1.2 Hz), in which the raw effect corresponds to the difference between the mean SNR-corrected amplitudes in the control task and the binary imagery condition. Our meta-analysis revealed an average raw mean difference of 0.041 µV, such that the SNR-corrected amplitude of brain activity was 0.041 µV higher during binary imagery compared to the control task. This effect ranged from -0.012 to 0.093 µV across the included studies. The 95% meta-analytic confidence interval ranged from -0.0004 to 0.082 µV, overlapping with zero (*p* = .052). In none of the 13 replication attempts did the 95% confidence interval overlap with the raw effect size from Nozaradan et al. (2011) (0.12); all 13 intervals were smaller than the one reported in Nozaradan et al. (2011).

**Figure 4.**
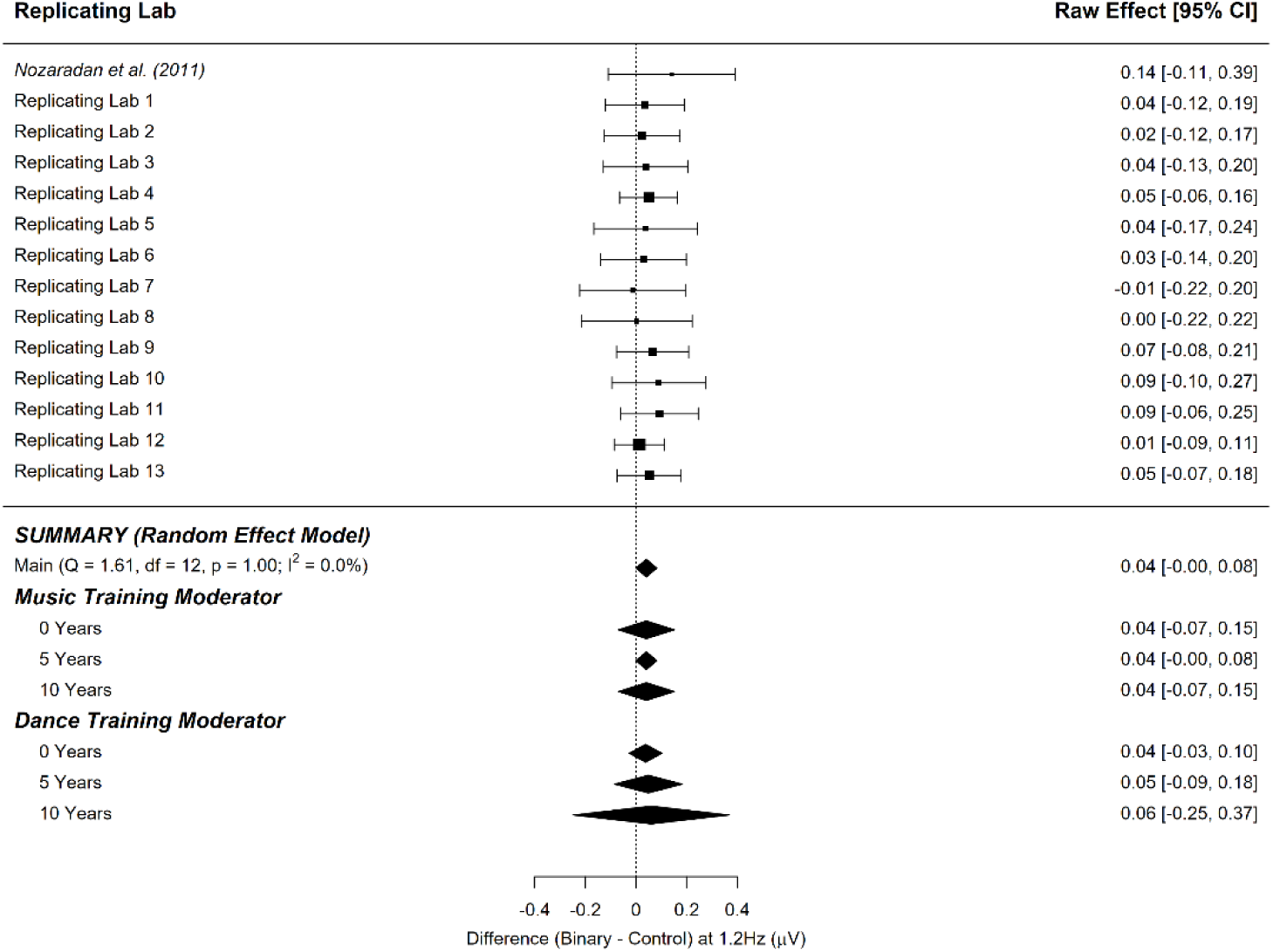
Point-estimate for the meta-analytic effect of condition (control task vs. binary imagery) on the binary frequency. Studies are listed alphabetically by the last name of the first author for that lab’s contribution, with the original study presented at the top. Squares indicate each lab’s mean difference, where the size of the square corresponds to the inverse of the standard error of the difference score, and the error bars indicate 95% confidence intervals (CI) around the mean difference. Diamonds indicate the random-effects meta-analytic effect size estimate, where the width represents the 95% CI. The first diamond represents a meta-analysis with no moderators. The next three diamonds represent the meta-analytic estimate with music training included as a moderator, and the final three diamonds represent the meta-analytic estimate with dance training included as a moderator. Note that none of the meta-analyses include the original Nozaradan et al. (2011) result.

The variability in the effect size among the studies (i.e., heterogeneity) was insignificant (*τ* = 0.00, *I^2^* = 0.00%, *H^2^* = 1.00), and the lack of detectable heterogeneity among the study effect sizes suggests that moderating variables may have little to no effect on the meta-analytic estimate of the effect size (*Q_13_* = 1.606, *p* = .999). Figure 4 shows that the effect of condition (control task vs. binary imagery) on the SNR-corrected amplitude of brain activity at the binary frequency was not substantially moderated by music training and was not substantially moderated by dance training.

##### Binary Active Effect

Figure 5 shows the point-estimate for the meta-analytic effect of condition on the binary frequency, in which the raw effect corresponds to the difference between the binary imagery condition and the ternary imagery condition. Our meta-analysis revealed an average raw mean difference of 0.032 µV, such that the SNR-corrected amplitude of brain activity was 0.032 µV higher during binary imagery compared to ternary imagery. This effect ranged from -0.012 to 0.120 µV across the included studies. The 95% meta-analytic confidence interval ranged from -0.010 to 0.075 µV, overlapping with zero (*p* = .132). In one out of 13 replication attempts, the 95% confidence interval overlapped the raw effect size from Nozaradan et al. (2011) (0.12 µV); of the 12 intervals that did not overlap the original effect size, all were smaller than the one reported in Nozaradan et al. (2011).

**Figure 5.**
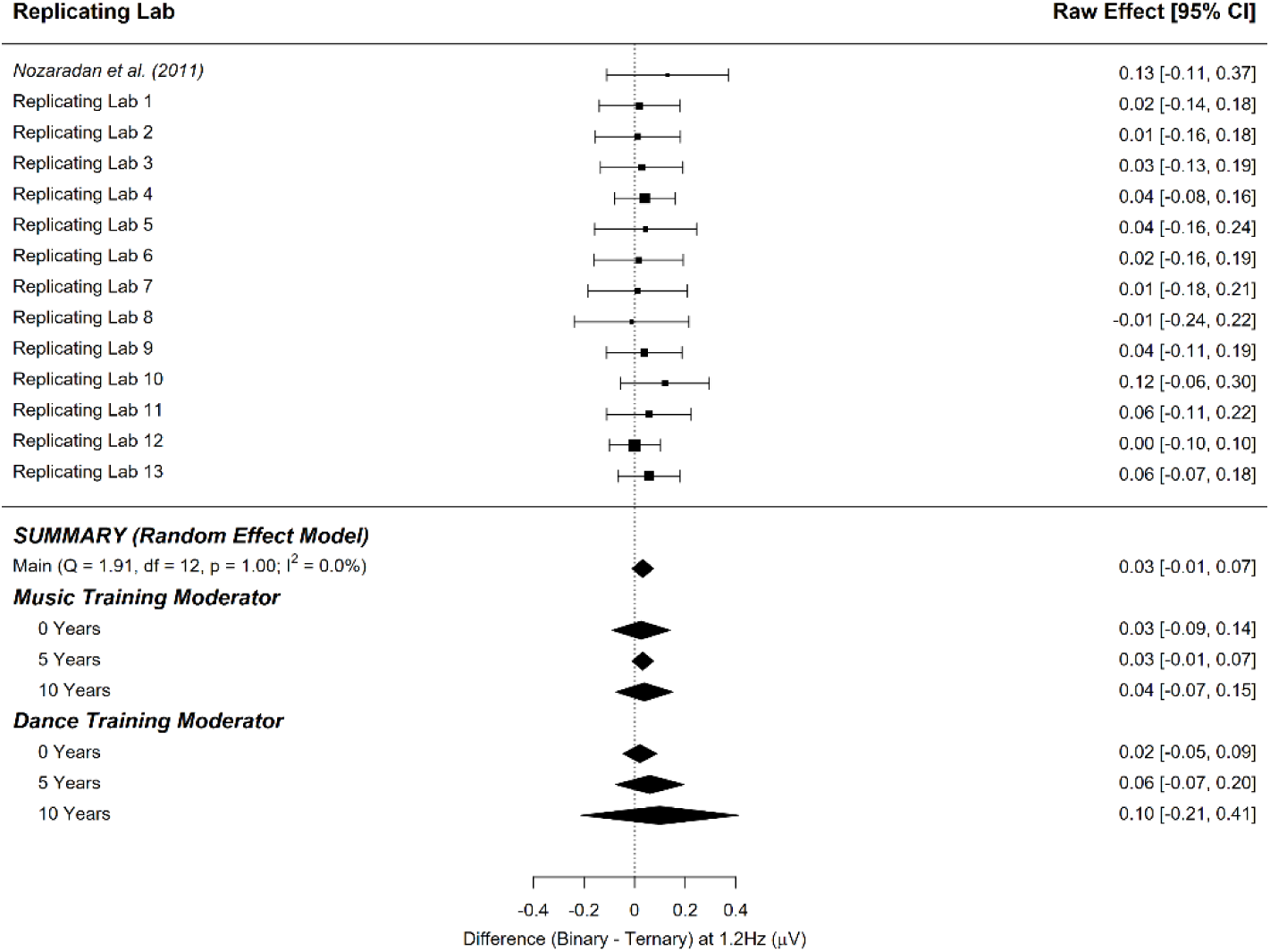
Point-estimate for the meta-analytic effect of condition (binary imagery vs. ternary imagery) on the binary frequency. Studies are listed alphabetically by the last name of the first author for that lab’s contribution, with the original study presented at the top. Squares indicate each lab’s mean difference, with the size of the square corresponding to the inverse of the standard error of the difference score, and the error bars indicate 95% confidence intervals (CI) around the mean difference. Diamonds indicate the random-effects meta-analytic effect size estimate, where the width represents the 95% CI. The first diamond represents a meta-analysis with no moderators. The next three diamonds represent the meta-analytic estimate with music training included as a moderator, and the final three diamonds represent the meta-analytic estimate with dance training included as a moderator. Note that none of the meta-analyses include the original Nozaradan et al. (2011) result.

The variability in the effect size among the studies (i.e., heterogeneity) was insignificant (*τ* = 0.00, *I^2^* = 0.00%, *H^2^* = 1.00), and the lack of detectable heterogeneity among the study effect sizes suggests that moderating variables may have little to no effect on the meta-analytic estimate of the effect size (*Q_13_*=1.907, *p* = .999). Figure 5 shows that the overall effect of condition (binary imagery vs. ternary imagery) on the SNR-corrected amplitude of brain activity at the binary frequency was not substantially moderated by music training and was not substantially moderated by dance training.

#### 6.2.2 Effect of Condition on Brain Activity at Ternary Frequency

##### Ternary Control Effect

Figure 6 shows the point-estimate for the meta-analytic effect of condition on the ternary frequency, where the raw effect corresponds to the difference between the control task and the ternary imagery condition. Our meta-analysis revealed an average raw mean difference of 0.034 µV, such that the SNR-corrected amplitude of brain activity was 0.034 µV higher during ternary imagery compared to the control task. This effect ranged from 0.005 to 0.149 µV across the included studies. The 95% meta-analytic confidence interval ranged from -0.010 to 0.077 µV, overlapping with zero (*p* = .126). In three out of 13 replication attempts, the 95% confidence interval overlapped the mean effect size from Nozaradan et al. (2011) (0.12 µV); of the 10 intervals that did not overlap the original effect size, all were smaller than the one reported in Nozaradan et al. (2011).

**Figure 6.**
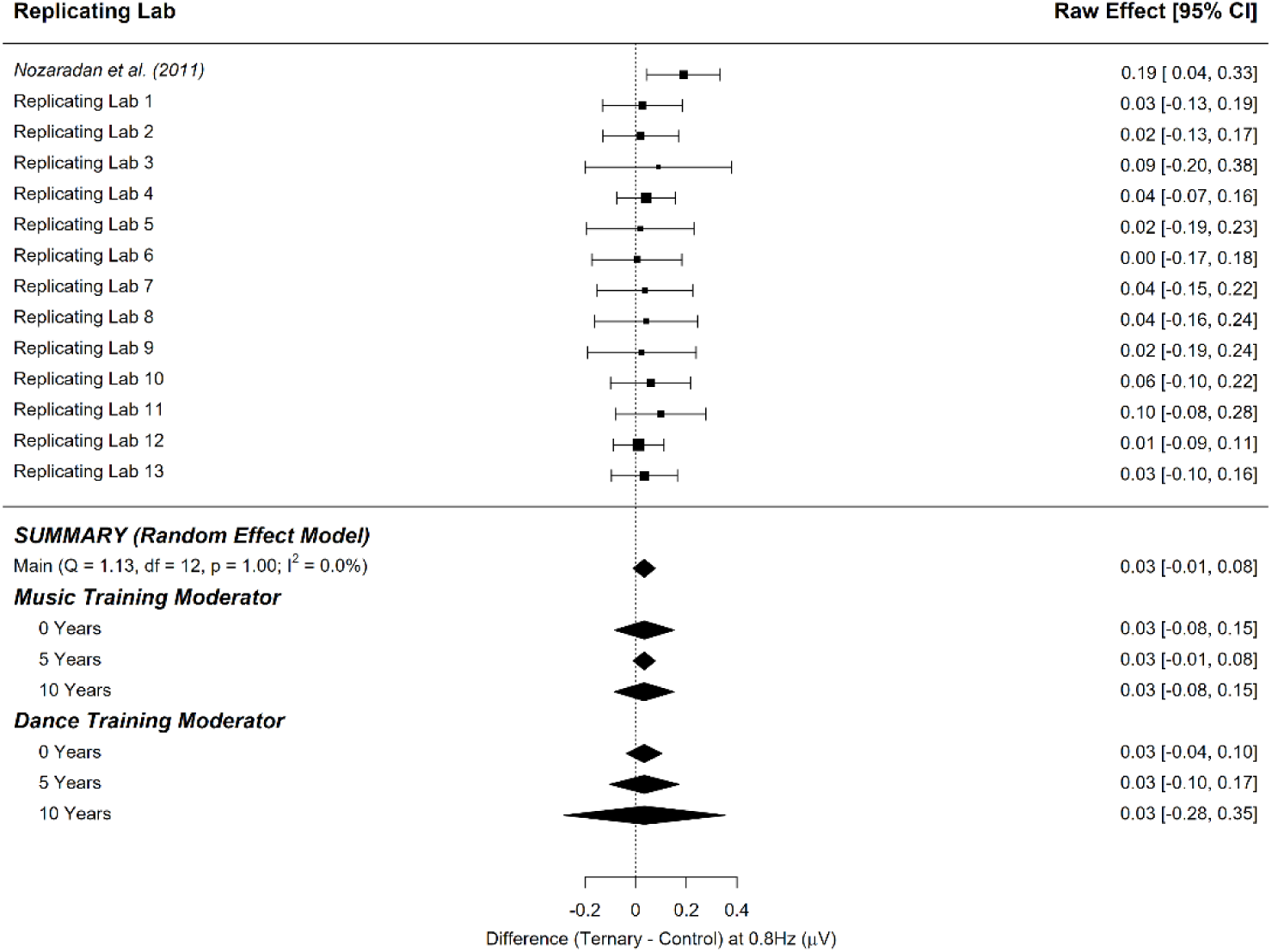
Point-estimate for the meta-analytic effect of condition (control task vs. ternary imagery) on the ternary frequency. Studies are listed alphabetically by the last name of the first author for that lab’s contribution, with the original study presented at the top. Squares indicate each lab’s mean difference, where the size of the square corresponds to the inverse of the standard error of the difference score, and the error bars indicate 95% confidence intervals (CI) around the mean difference. Diamonds indicate the random-effects meta-analytic effect size estimate, where the width represents the 95% CI. The first diamond represents a meta-analysis with no moderators. The next three diamonds represent the meta-analytic estimate with music training included as a moderator, and the final three diamonds represent the meta-analytic estimate with dance training included as a moderator. Note that none of the meta-analyses include the original Nozaradan et al. (2011) result.

The variability in the effect size among the studies (i.e., heterogeneity) was insignificant (*τ* = 0.00, *I^2^* = 0.00%, *H^2^* = 1.00), and the lack of detectable heterogeneity among the study effect sizes suggests that moderating variables may have little to no effect on the meta-analytic estimate of the effect size (*Q_13_* = 1.132, *p* = 1.00). Figure 6 shows that the overall effect of condition (control task vs. ternary imagery) on the SNR-corrected amplitude of brain activity at the ternary frequency was not substantially moderated by music training and was not substantially moderated by dance training.

##### Ternary Active Effect

Figure 7 shows the point-estimate for the meta-analytic effect of condition on the ternary frequency, with the raw effect corresponding to the difference between the ternary imagery condition and the binary imagery condition. Our meta-analysis revealed an average raw mean difference of 0.035 µV, such that the SNR-corrected amplitude of brain activity was 0.035 µV higher during ternary imagery compared to binary imagery. This effect ranged from -0.033 to 0.089 µV across the included studies. The 95% meta-analytic confidence interval ranged from -0.008 to 0.077 µV, overlapping with zero (*p* = .111). In three out of 13 replication attempts, the 95% confidence interval overlapped the raw effect size from Nozaradan et al. (2011) (0.12 µV); of the 10 intervals that did not overlap the original effect size, all were smaller than the one reported in Nozaradan et al. (2011).

**Figure 7.**
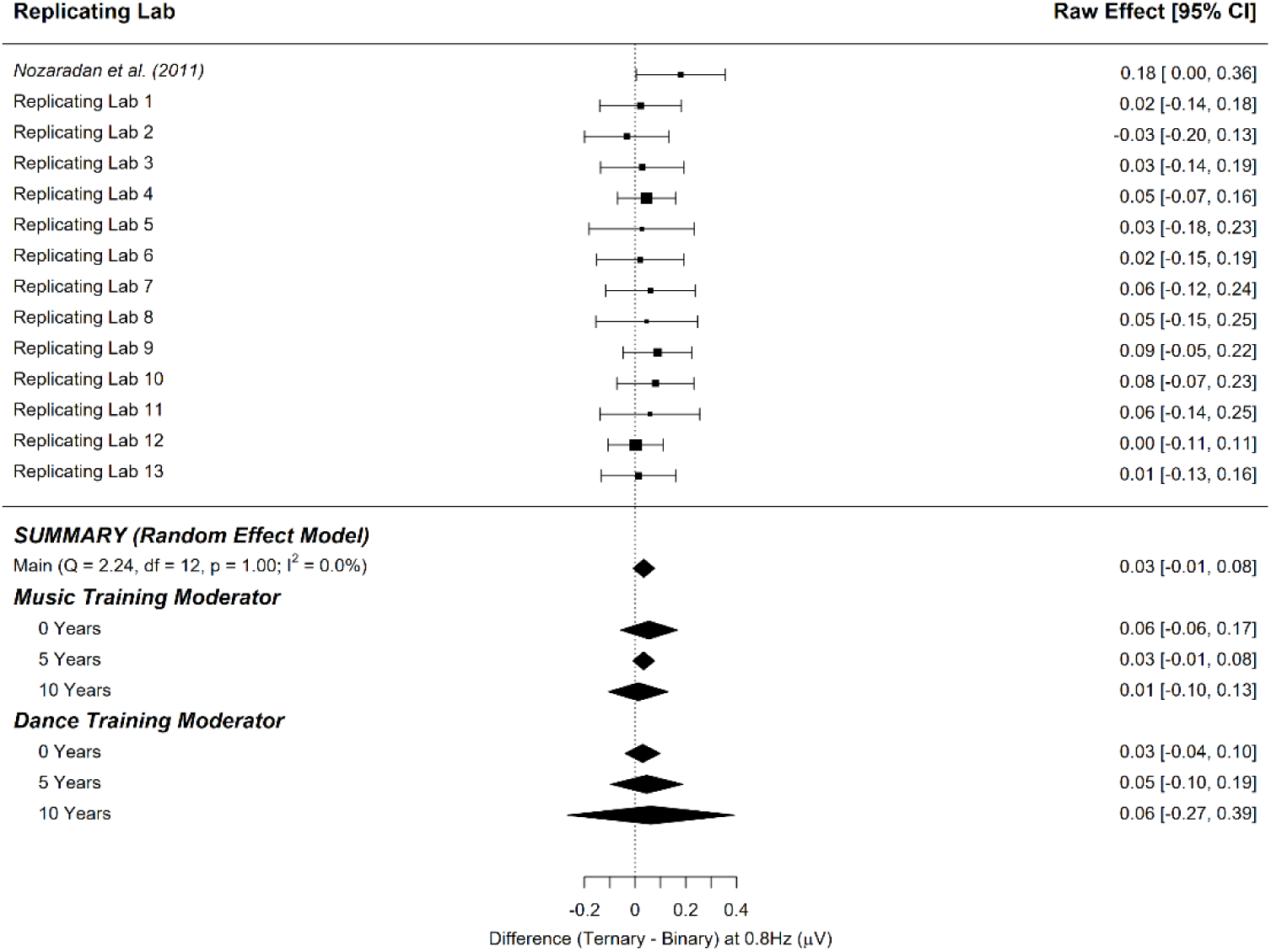
Point-estimate for the meta-analytic effect of condition (ternary imagery vs. binary imagery) on the ternary frequency. Studies are listed alphabetically by the last name of the first author for that lab’s contribution, with the original study presented at the top. Squares indicate each lab’s mean difference, where the size of the square corresponds to the inverse of the standard error of the difference score, and the error bars indicate 95% confidence intervals (CI) around the mean difference. Diamonds indicate the random-effects meta-analytic effect size estimate, where the width represents the 95% CI. The first diamond represents a meta-analysis with no moderators. The next three diamonds represent the meta-analytic estimate with music training included as a moderator, and the final three diamonds represent the meta-analytic estimate with dance training included as a moderator. Note that none of the meta-analyses include the original Nozaradan et al. (2011) result.

The variability in the effect size among the studies (i.e., heterogeneity) was insignificant (*τ* = 0.00, *I^2^* = 0.00%, *H^2^* = 1.00), and the lack of detectable heterogeneity among the study effect sizes suggests that moderating variables may have little to no effect on the meta-analytic estimate of the effect size (*Q_13_* = 2.237, *p* = .999). Figure 7 shows that the overall effect of condition (ternary imagery vs. binary imagery) on the SNR-corrected amplitude of brain activity at the ternary frequency was not substantially moderated by music training and was not substantially moderated by dance training.

#### 6.2.3 Summary of Meta-Analytic Effects of Condition

The meta-analytic estimates of the effect of condition on the imagery-related SNR-corrected amplitude estimated a raw difference of 0.03 – 0.04 µV when comparing the means of the corresponding beat imagery condition to either the other imagery condition or the control condition. All four estimated effect sizes had confidence intervals overlapping zero, providing no evidence against the null hypothesis (*p* ranges from *p* = .052 to *p* = .132). However, it is important to note that of the thirteen labs included in this analysis, eight produced data that violated assumptions of sphericity and thus required a Greenhouse Geiser correction (see Table 1). Thus, we also conducted our pre-registered meta-analytic estimates using raw differences between medians, instead of means, for the four estimated effect sizes in an exploratory analysis (see section 7.2).

### 6.3 One-Way ANOVAs and Post-Hoc Tests on Effect of Condition

Table 2 shows the resulting effect sizes (as measured by partial eta squared) of repeated measures one-way ANOVAs conducted for each individual lab on the effect of condition (control, binary, ternary), in which the dependent variable is the SNR-corrected amplitude of brain activity at the frequency of interest. The results of the original study are shown at the top, followed by all participating labs. Table 3 shows the results of post hoc t-tests conducted for each individual lab following the one-way ANOVAs for each frequency of interest. Specifically, these post hoc tests are conducted in accordance with the original study, such that for each lab, a t-test is conducted to demonstrate any significant difference in SNR-corrected amplitude of the brain activity between any two conditions (control vs. binary imagery, control vs. ternary imagery, and binary imagery vs. ternary imagery) at each of the frequencies of interest (stimulus [2.4 Hz], binary [1.2 Hz], ternary [0.8 Hz], and 1^st^ ternary harmonic [1.6 Hz]), resulting in nine *t*-tests for each of the participating labs. Results are reported for each lab, including the original study.

Of the labs who met the inclusion criteria for the study: 5 out of 13 labs showed a significant condition effect on the SNR-corrected amplitude of the ternary imagery SSEP (0.8Hz), 1 out of 13 labs showed a significant condition effect on the SNR-corrected amplitude of the binary imagery SSEP (1.2Hz), 5 out of 13 labs showed a significant condition effect on the SNR-corrected amplitude of the ternary harmonic SSEP (1.6Hz), and 7 out of 13 labs showed no evidence for a condition effect on the SNR-corrected amplitude of the Stimulus SSEP (2.4Hz). After accounting for multiple comparisons, only 2 labs replicated the condition effect at the ternary SSEP (0.8Hz), and only 3 labs replicated the condition effect at the binary SSEP (1.2Hz), with no labs replicating more than one effect. No labs showed a main effect of condition in all four ANOVAs, even before accounting for multiple comparisons. While 5 out of 13 labs showed a significant effect of condition on the Stimulus SSEP (2.4Hz), such that greater SNR-corrected amplitudes were observed for the imagery conditions compared to the control condition, only 1 of these results survived correction for multiple comparisons. All labs demonstrated average SSEPs at 2.4Hz that were statistically greater than 0 for all conditions, shown in Table S2.

To further investigate the relation between beat-related neural activity and beat-related task performance or participant factors, correlations were conducted on the magnitude of stimulus-specific and imagery-specific SSEPs, performance (accuracy and imagery success ratings) on the probe task performed during imagery blocks, music training (in years), and dance training (in years). These are presented below in Table 4, both for each individual lab and as a grand average. Note that the raw effect size estimates are also included in the correlation matrix, but these correlations were exploratory as they were not pre-registered (see section 7.5).

**Table 4.**
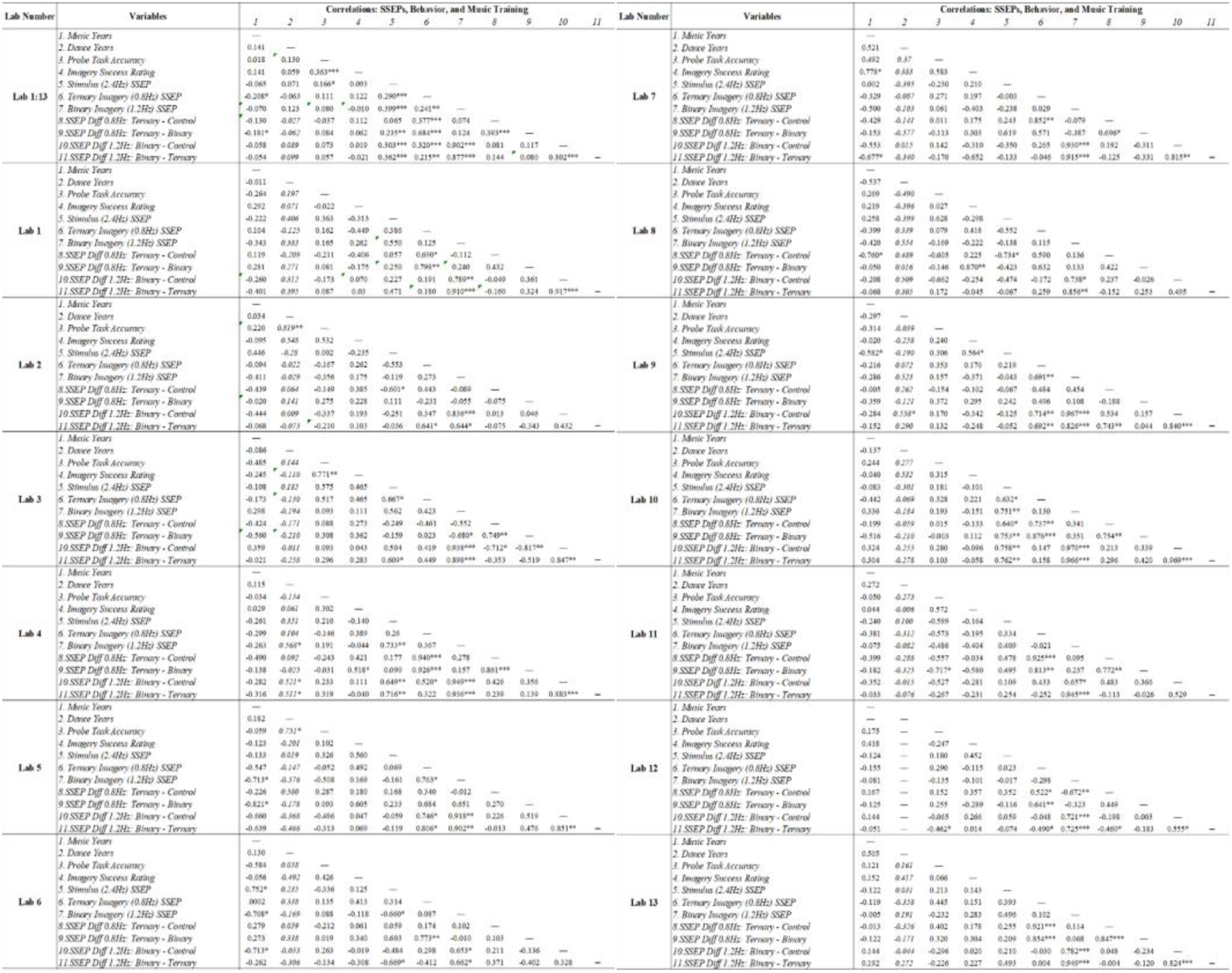
Results of Pearson’s correlations conducted across the entire dataset and by individual lab among the following variables: music training in years [Music Years], dance training in years [Dance Years], performance on the probe task [Probe Task Accuracy], confidence ratings of imagery success [Imagery Success Rating], SNR-corrected amplitude (EEG) in microvolts at the stimulus frequency (2.4Hz) [Stimulus SSEP], SNR-corrected amplitude (EEG) in microvolts at the binary beat frequency (1.2 Hz) during binary imagery [Binary Imagery SSEP], SNR-corrected amplitude (EEG) in microvolts at the ternary beat frequency (0.8 Hz) during ternary imagery [Ternary Imagery SSEP], and the four raw difference effects estimates by the meta-analyses. Note that correlations with the SSEP raw difference scores were not pre-registered and are thus considered exploratory. ^+^Correlations conducted with imagery ratings and probe accuracy were conducted within each respective task, such that binary imagery SSEP was correlated with probe accuracy/imagery success ratings from the binary trials only (and vice versa for ternary imagery). In correlations with music, dance, or with each other, the grand average of imagery ratings and probe accuracy were used.

### 6.4 Relation Between Brain Activity and Other Factors

The GEE binary logistic regression on nested trial data showed that the SNR-corrected amplitude of the Stimulus SSEP (Wald χ^2^ = 4.944, *p* = .026) was the only significant predictor of trial accuracy. Specifically, a higher magnitude of brain activity at the stimulus-frequency (2.4 Hz) predicted an accurate participant response on the probe task. No other significant predictors were observed. Parameter estimates and the modeled relationship with accuracy are shown in Figure 8 for all predictors entered into the model.

**Figure 8.**
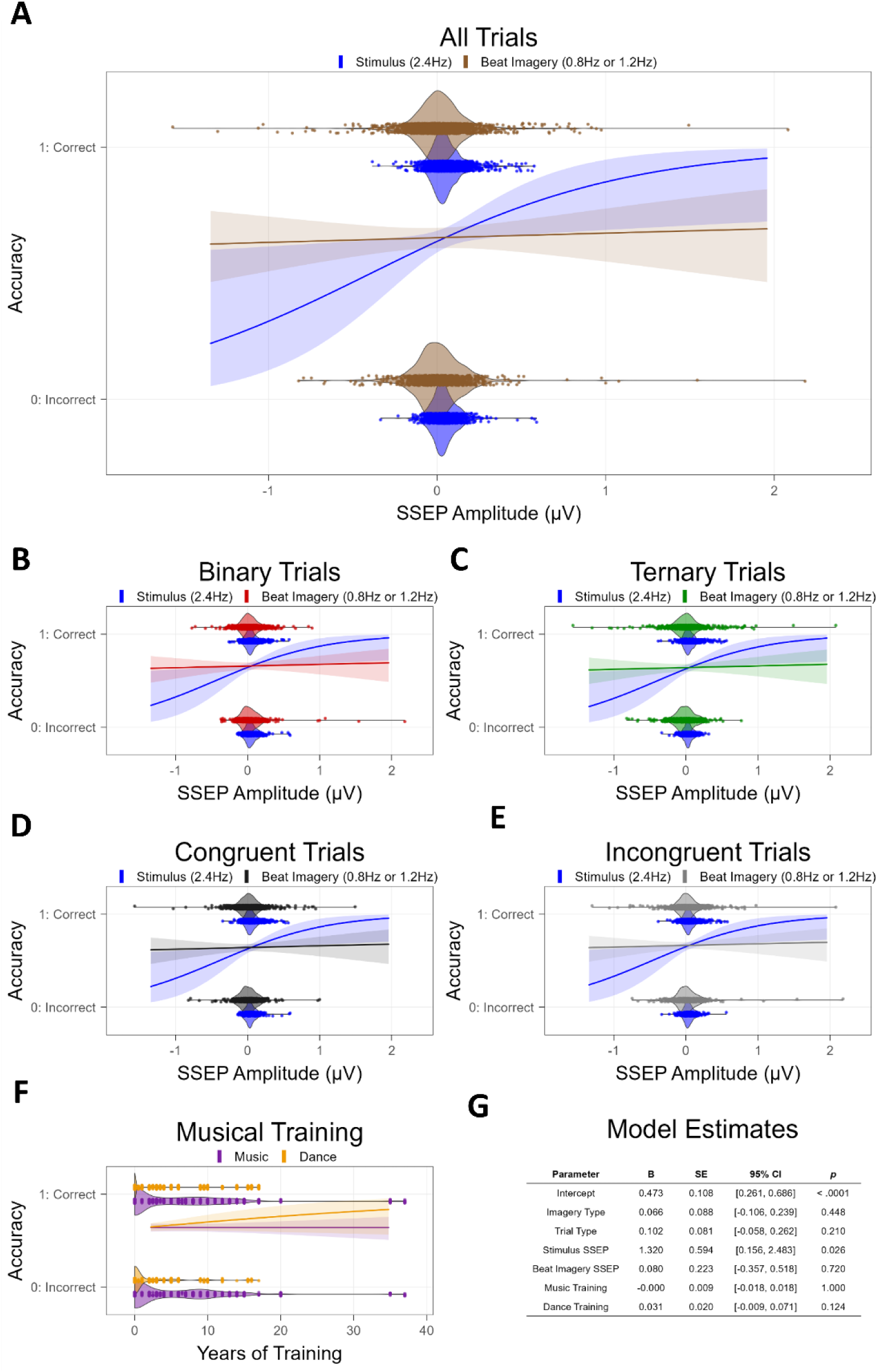
Results of trial-by-trial logistic regression examining whether frequency-tagged neural responses, musical training, and/or study factors predict performance on the imagery-related probe task. Raw values of continuous predictors are plotted along the x-axis, and accuracy on the probe task is plotted on the y-axis (0: incorrect response, 1: correct response). SSEP predictors represent SNR-corrected amplitude at the respective frequency in the EEG signal. A) Model predictions on all trials for SSEP predictors. B) Model predictions on binary imagery trials for SSEP predictors. C) Model predictions on ternary imagery trials for SSEP predictors. D) Model predictions on congruent (ON beat) trials for SSEP predictors. E) Model predictions on incongruent (OFF beat) trials for SSEP predictors. F) Model predictions on all trials for musical training predictors (Music, Dance). G) Resulting parameter estimates from trial-by-trial logistic regression.

## 7 Exploratory Analyses

### 7.1 Effect of Micro-Movements on SSEPs to Imagined Beat

While participants were asked to refrain from movement during the three conditions and the experimenter recorded any observable rhythmic movement by the participant (only observed in 1 participant, who was excluded), it is still possible that participants performed unobservable micro-movements to maintain the rhythmic structure, and such movement could contaminate the EEG response. Because the procedure to measure muscle activity was optional (SCM muscle in the neck and the FDI muscle in the hand) and due to the additional constraints of the COVID-19 pandemic, only one lab was able to collect such data (Lab #8). Of the eight participants from this lab (after pre-registered exclusions), only four participants had viable data from the hand and neck (due to issues in the signal reported by the experimenter). Thus, we did not have sufficient power to estimate the contribution of overt movement to the measured imagery -related SSEPs. Despite this, the movement log recorded by experimenters indicated that only one participant was observed moving rhythmically and was excluded. In addition, more recent research suggests that unobservable micromovements are not likely to contribute to the imagery-related SSEPs during rhythmic imagery, confirmed via both EMG response and motion capture (Cheng et al., 2021).

### 7.2 Meta-Analyses Using Median Raw Differences

After conducting the Univariate ANOVAs for the individual labs (see section 6.4 in Results and Table 2), it became clear that several lab samples violated the assumption of sphericity, suggesting non-normally distributed data. To this end, we conducted four exploratory meta-analyses: the same meta-analytic estimates from the pre-registered analysis plan were conducted using medians for each lab’s condition estimates instead of means. Exploratory meta-analyses on medians were conducted using the R package metamedian (McGrath et al., 2024).

#### 7.2.1 Effect of Condition on Brain Activity at Binary Frequency

##### Binary Control Effect

Figure S2 shows the point-estimate for the meta-analytic effect of condition on the binary frequency, in which the raw effect corresponds to the median difference between the control task and the binary imagery condition. Our meta-analysis revealed an average raw median difference of 0.025 µV, such that the SNR-corrected amplitude of brain activity was 0.025 µV higher during binary imagery compared to the control task. This effect ranged from -0.002 to 0.092 µV across the included studies. The 95% meta-analytic confidence interval ranged from 0.014 to 0.049 µV, not overlapping with zero (*p* < .001). In 0 out of 13 replication attempts, the third interquartile range overlapped the effect size from Nozaradan et al. (2011) (0.12 µV); 11 of the 13 intervals that did not overlap the original effect size were smaller than the one reported in Nozaradan et al. (2011). The variability in the effect size among the studies (i.e., heterogeneity) was significant (*τ* = 0.0002, *I^2^* = 49.6%, *H^2^* = 1.99, *Q_13_* = 25.901, *p* = .011). The increased variability among the study effect sizes leaves open the possibility that the strength of the effect varies as a function of one or more moderating variables.

##### Binary Active Effect

Figure S3 shows the point-estimate for the meta-analytic effect of condition on the binary frequency, in which the raw effect corresponds to the median difference between the binary imagery condition and the ternary imagery condition. Our meta-analysis revealed an average raw median difference of 0.024 µV, such that the SNR-corrected amplitude of brain activity was 0.024 µV higher during binary imagery compared to ternary imagery. This effect ranged from -0.010 to 0.104 µV across the included studies. The 95% meta-analytic confidence interval ranged from 0.013 to 0.036 µV, not overlapping with zero (*p* < .001). In none out of 13 replication attempts did the third interquartile range overlapped the effect size from Nozaradan et al. (2011) (0.12 µV); Of the 13 intervals that did not overlap the original effect size, all were smaller than the one reported in Nozaradan et al. (2011). The variability in the effect size among the studies (i.e., heterogeneity) was significant (*τ* = 0.0003, *I^2^*= 60.5%, *H^2^* = 2.53, *Q_13_* = 30.047, *p* = .003). The increased variability among the study effect sizes leaves open the possibility that the strength of the effect varies as a function of one or more moderating variables.

#### 7.2.2 Effect of Condition on Brain Activity at Ternary Frequency

##### Ternary Control Effect

Figure S4 shows the point-estimate for the meta-analytic effect of condition on the ternary frequency, where the raw effect corresponds to the difference between the control task and the ternary imagery condition. Our meta-analysis revealed an average raw median difference of 0.029 µV, such that the SNR-corrected amplitude of brain activity was 0.029 µV higher during ternary imagery compared to the control task. This effect ranged from 0.001 to 0.072 µV across the included studies. The 95% meta-analytic confidence interval ranged from 0.018 to 0.039 µV, not overlapping with zero (*p* < .001). In 2 out of 13 replication attempts, the third interquartile range overlapped the effect size from Nozaradan et al. (2011) (0.12 µV); Of the 11 intervals that did not overlap the original effect size, all were smaller than the one reported in Nozaradan et al. (2011). The variability in the effect size among the studies (i.e., heterogeneity) was insignificant (*τ* = 0.0001, *I^2^* = 29.3%, *H^2^* = 1.42), and the lack of detectable heterogeneity among the study effect sizes suggests that moderating variables may have little to no effect on the meta-analytic estimate of the effect size (*Q_13_* = 15.571, *p* = .212).

##### Ternary Active Effect

Figure S5 shows the point-estimate for the meta-analytic effect of condition on the ternary frequency, with the raw effect corresponding to the difference between the ternary imagery condition and the binary imagery condition. Our meta-analysis revealed an average raw median difference of 0.031 µV, such that the SNR-corrected amplitude of brain activity was 0.031 µV higher during ternary imagery compared to binary imagery. This effect ranged from -0.02 to 0.097 µV across the included studies. The 95% meta-analytic confidence interval ranged from 0.015 to 0.049 µV, not overlapping with zero (*p* < .001). In 3 out of 13 replication attempts, the third interquartile range overlapped the effect size from Nozaradan et al. (2011) (0.12 µV); Of the 10 intervals that did not overlap the original effect size, all were smaller than the one reported in Nozaradan et al. (2011). The variability in the effect size among the studies (i.e., heterogeneity) was significant (*τ* = 0.0006, *I^2^* = 0.714%, *H^2^* = 3.49, *Q_13_* = 39.586, *p* < .0001). The increased variability among the study effect sizes leaves open the possibility that the strength of the effect varies as a function of one or more moderating variables.

#### 7.2.3 Summary of Exploratory Meta-Analytic Effects of Condition

Given the violations of sphericity present in the individual lab’s data, the use of medians provides some valuable insight into differences among conditions, compared to the pre-registered estimates of differences among means (see section 6.3). The exploratory meta-analytic estimates of the effect of condition on the imagery-related SNR-corrected amplitude (SSEP) estimated a raw difference of 0.02 µV– 0.03 µV when comparing the medians of the corresponding beat imagery condition to either the other imagery condition or the control condition, which were relatively comparable in magnitude to the estimates obtained with means (0.03 – 0.04 µV) . Unlike the estimates with means, the estimates of effect sizes using medians had confidence intervals that did not overlap zero, providing some evidence against the null hypothesis (all *p*’s < .001). Despite the resulting evidence for significant effects related to the experimental manipulation (i.e., beat imagery), the size of the estimated effects is indeed quite small. The pre-registered meta-analytic estimates (using differences of means) also detected some heterogeneity when using medians but not with means. While the presence of heterogeneity leaves open the possibility that some moderating variable(s) may explain some of the variability among the contributing labs, the R package used for meta-analytic estimates of medians (metamedian) does not offer the capability to test the effects of moderating variables. Future research may still want to focus on potential moderating variables, particularly using more sensitive measures of musical aptitude, such as by measuring musical sophistication (e.g., using the Goldsmiths Musical Sophistication Index, Müllensiefen et al., 2014)) or by measuring beat perception/entrainment skills using the using the Beat Alignment Test (BAT) (Iversen & Patel, 2008).

### 7.3 Analysis of Behavioral Responses on Probe Task

To examine variability in performance on the Probe Task on the basis of our experimental factors, we conducted a mixed effects ANOVA for both probe task accuracy and imagery success ratings, each with two between participant factors (imagery order, lab membership) and two within participant factors (imagery type, probe type), where imagery order has two levels (binary first, ternary first), lab membership has thirteen levels, imagery type has two levels (binary, ternary), and probe type has two levels (“ON” beat, “OFF” beat).

Regarding probe task accuracy, there was a significant main effect of lab, *F*(1,126) = 2.067, *p* = .024, *η_p_ ^2^*= .16, and a significant probe type x imagery order interaction, *F*(1,126) = 7.808, *p* = .006, *η_p_ ^2^*= .06, with no other significant main effects nor interactions (all *p* > .05). Post-hoc tests revealed no significant differences among labs after correcting for multiple comparisons. In addition, accuracy was significantly greater for the binary probe than the ternary probe trials, but only for participants who completed ternary imagery first, *t*(126) = 3.321, *p* = .007, *Cohen’s d* = .30. Regarding imagery success ratings, there was a significant main effect of imagery type, *F*(1,126) = 15.899, *p* < .001, *η_p_^2^*= .11, and a significant probe type x imagery order interaction, *F*(1,126) = 6.205, *p* = .014, *η_p_^2^*= .05, with no other significant main effects or interactions (*p* > .05). Post hoc tests revealed significantly greater imagery success ratings for binary imagery compared to ternary imagery, *t*(126) = 3.987, *p* < .001, *Cohen’s d* = .26. Post-hoc comparisons exploring the probe type x imagery order interaction were all non-significant after correcting for multiple comparisons. Results from the probe task are presented in Figure 2. There was a significant correlation between probe task accuracy and imagery success ratings (*r* = 0.36, *p* < .001).

### 7.4 Exploratory Tests of the Relation Between Brain Activity and Other Factors

In order to further explore the relation between brain activity and behavioral performance, we performed the same pre-registered binary logistic regression separately for each imagery condition, removing “imagery type” as a factor. For ternary imagery trials, the SNR-corrected amplitude of the stimulus SSEP (Wald χ^2^ = 13.235, *p* < .001) and trial type (Wald χ^2^ = 6.743, *p* = .009) were significant predictors of trial accuracy, such that a higher SNR-corrected amplitude at the stimulus-frequency predicted higher accuracy on the probe task, and more correct responses were given for the OFF probe than the ON probe. No other significant predictors were observed. For binary imagery trials, none of the predictors were statistically significant. See Figure S6 for model estimates and predictor plots versus accuracy.

In addition, we explored additional correlations between the raw condition effect SSEP differences (used for the meta-analyses) with the behavioral task measures and the music training factors. While the SSEPs reported here are SNR-corrected and thus less prone than absolute amplitudes to extraneous participant factors (e.g., skull thickness), the raw SSEP differences between conditions may provide a more standardized measure of neural fluctuations in beat-related SSEPs. Thus, the four raw effect sizes investigated in the pre-registered meta-analyses were added to the correlation matrix as an exploratory investigation (see Table 4).

## 8 General Discussion

This Registered Report included data from 13 laboratories for a total of 152 participants, after lab- and individual-dataset exclusions. The data were collected by following a vetted experimental design with a preregistered analysis plan. The first aim of this RR was to estimate the true effect sizes reported in the original 2011 study by performing meta-analyses across a multi-lab design. The overall results demonstrate that the original effect of imagery reported by Nozaradan and colleagues (imagining that an amplitude-modulated repeating pure tone has a binary or ternary stress pattern) is much smaller than originally reported when tested with a larger overall sample than the original study. In particular, the pre-registered meta-analytic effect sizes tested ranged from 0.03 to 0.04 µV, whereas the original effect sizes ranged from 0.13 to 0.19 µV, consistent with studies showing that replicated effects are typically smaller and less likely to be statistically significant than the original effects (Open Science Collaboration, 2015). While all four of the pre-registered meta-analytic effects (calculated with means for each lab) produced confidence intervals that overlapped with 0, exploratory analyses using medians (due to multiple violations of sphericity across labs) did result in all four meta-analytic effect sizes having confidence intervals that did not overlap with 0. Furthermore, the (non-pre-registered) analyses that included all 152 participants together did result in significant effects of imagining a beat on brain activity. This suggests that if the effect of periodic imagery on brain activity is a true effect, then sample sizes even larger than our overall sample are likely necessary to detect them reliably. Because the effect sizes related to condition comparisons (e.g., binary beat vs. ternary beat) are estimated by the current study to be quite small and require a very large sample to detect, frequency-tagging may not be a feasible method to explore neural correlates of musical beat perception. Importantly, while the repeated-measures ANOVA results of imagery produced significant effects of condition at the grand average level (*N* = 152) for the imagery-related frequencies (binary: 1.2 Hz, ternary: 0.8 Hz, 1.6 Hz), there was also an additional significant effect of condition at the stimulus frequency (2.4 Hz), such that imagery conditions resulted in larger SSEPs than the control condition, and in two of four tests there was a significant condition x lab ID interaction. This may have implications for the effect of task demands on the size of SSEPs measured in response to auditory rhythm.

Due to the small sample size of the original study and the high amounts of musical training in the participants from the original study, we also tested for moderating effects of both musical and dance experience in our pre-registered meta-analyses. None of these moderation effects were significant. One weakness to this analysis is that only the average number of music years or dance years is considered for each lab, which reduces the influence of individual differences. To this point, we also conducted correlations between music/dance years of training and the four meta-analytic effects on a participant basis, and all correlations were minor and not significant (range of Pearson’s *r*: -.181 to .099, all *p*’s > .05). Our data showed no positive correlation between years of training and the SNR-corrected amplitude of imagery-related SSEPs, even across our entire sample (N = 152) . Contrary to our hypothesis, there was a significant negative correlation between music years of training and the SNR-corrected amplitude at the ternary beat imagery across the sample. Probing such a link, assuming it exists, may need a more robust measure of musical aptitude and experience, in addition to years of training. Alternatively, it is possible that musical expertise does not have a significant bearing on the strength of neural oscillations to perceived metrical structures. Indeed, while some prior evidence suggests significant group effects on frequency-specific EEG measures between musicians and non-musicians, a recent multisite registered replication report found that one of the seminal pieces of evidence for this sentiment – that musicians show enhanced frequency following responses (FFRs) to speech syllables as compared to nonmusicians – does not replicate (Whiteford et al., 2025). Nevertheless, our lack of evidence for a relation between music training and neural oscillations is perhaps best considered inconclusive, because of the small size of the main effect of beat imagery on brain activity. It is likely that any moderation by musical or dance experience would only change the condition effect by a small fraction; therefore, to reliably detect any effect of music training, one would likely require several times the number of participants we tested across in this study.

The second main aim of this RR was to assess the relationship between imagery-related neural responses (SSEP amplitude) and imagery-related behavioral responses. Surprisingly, the imagery-related stimulus SSEP amplitude (binary: 1.2 Hz, ternary: 0.8 Hz) was not related to the performance on the imagery-related task, but rather the SNR-corrected amplitude of neural activity at the stimulus frequency (2.4Hz, the frequency of the periodic physical stimulus) was significantly predictive of performance, regardless of imagery condition. This raises questions about the assumption that the SNR-corrected amplitude of the imagery-related SSEP responses reflects top-down perception of beat and meter. The data in the current study suggest that rather than representing the strength of beat or meter entrainment, SSEPs may reflect something else, such as how well a stimulus was encoded or how much attention the participant was directing to the task (Gibbings et al., 2023; Saupe et al., 2009). This conclusion is also supported by the significant effect of condition that we observed for the stimulus SSEP (2.4 Hz), where the imagery conditions produced significantly larger amplitudes at this frequency, compared to the control condition. Of note, the original study found no such effect, suggesting that the influence of attentional resources on SSEPs should be considered in future work. Few prior studies have related the modulation of beat-related SSEPs to behavioral responses from the listener. While some published work suggests correlations between SSEPs and groove ratings (Cameron et al., 2019; Zalta et al., 2024) and temporal perturbation/precision (Nozaradan et al., 2016), the extent to which perception-related SSEPs represent general timing mechanisms versus beat- or meter-related neural processes remains unclear.

Indeed, one explanation is that the current imagery task is not suitable to measure beat or meter perception. However, the current study demonstrates a similar pattern of results relating brain (SSEPs) to behavior (task-related listener responses) as a recent investigation of SSEPs to induced (and not imagined) musical beat (Nave et al., 2022). To our knowledge, this is the only other existing investigation of beat-related SSEPs that controlled for stimulus properties *and* statistically compared them to behavioral responses from a concurrent beat-related task. Listeners judged whether drum probes matched the beat of a rhythm after a musical context naturally induced one of two beat percepts in the rhythm. Crucially, while results suggested group-level differences in SSEP amplitudes as predicted between the two beat contexts, Nave and colleagues (2022) found evidence that beat-related SSEPs were not always positively related to performance on the beat-related task. In the 2022 paper, a logistic regression model revealed a relation between the beat-related task performance and the beat-related SSEP amplitude that depended on the stimulus properties; larger beat-related SSEP amplitudes predicted better performance only for the ternary beat condition (but only for the faster tempo), while greater beat-related SSEPs predicted *worse* performance for the binary beat condition (but only for the slower tempo), and in all other conditions there was no significant relationship between the beat-related SSEP and performance.

In the current study, to further explore whether the beat-related SSEPs are perhaps related to beat-related performance *differently* based on imagery condition, we conducted two exploratory logistic regressions: one for binary trials and one for ternary trials. In both cases, beat-related SSEP amplitude was not a significant predictor of beat-related task accuracy. It is worth noting that, despite the fact that beat-related SSEPs were not a significant predictor of task accuracy in either imagery condition, the *direction* of the predictive relationship between beat-related SSEP amplitude and task accuracy depended on imagery condition (Binary imagery: Β = -0.551; Ternary imagery: Β = 0.487). This trend in the exploratory regressions results mirrors findings from the 2022 study, demonstrating an apparent positive relation between the ternary SSEP amplitude and performance on the ternary imagery task, and a negative relation between the binary SSEP amplitude and performance on the binary imagery condition (see Figure S6). Together, these results further suggest that frequency-tagged responses at the perceptual frequency (i.e., perceived beat or meter) do not necessarily correlate positively with listener performance on a perception-related task, and that researchers should use caution when reverse-inferring that frequency-tagged beat-related responses indicate beat perception.

Even though the effect sizes we detected were much smaller than the original study, many studies have been published since Nozaradan et al. (2011) that examine effects on beat-related SSEPs. Some of these studies report positive evidence for non-stimulus-driven modulatory effects of attention and learning on SSEPs (Celma-Miralles & Toro, 2019; Chemin et al., 2014; Gibbings et al., 2023; Nave et al., 2022), similar to the Nozaradan et al. (2011) study. However, most studies demonstrate effects that are more likely to be stimulus driven, such as low-complexity/highly-periodic vs. high-complexity/weakly-periodic rhythms (Edalati et al., 2023; Lenc et al., 2023; Mathias et al., 2020; Nozaradan et al., 2016), music vs. speech (Harding et al., 2019; Vanden Bosch der Nederlanden et al., 2020), auditory vs. non-auditory modalities (Celma-Miralles et al., 2016; Gilmore & Russo, 2021), and rhythms composed of low- vs. high-frequency tones (Lenc et al., 2018). While beat-related SSEP amplitude differences have been observed in missing pulse rhythms (e.g., no energy at 2Hz in the stimulus), the estimated beat-related (1Hz) SSEPs were “averaged over 10% of the sensors with the highest 2 Hz amplitude”, thereby excluding 90% of their data where the missing frequency of 2Hz was not represented in the brain by the listener (Tal et al., 2017b).

A positive feature of these SSEP studies since 2011 is that they generally used sample sizes larger than Nozaradan et al. (2011), which had a sample size of N = 8. Another positive feature of such studies is that they measured SSEPs (e.g., SNR-corrected amplitude peaks in the spectrum of brain activity at frequencies of interest) that can be isolated at expected frequencies in amplitude spectra of EEG activity, and whose presence is clearly related to experimental stimulation or manipulation and not to other oscillatory activity such as ongoing brain rhythms. This assessment of generally strong frequency-domain peaks is based in part on visual inspection of their spectral data plots, although some studies statistically tested whether SSEP peaks were significantly above a background noise level. On the other hand, very few of them (e.g. Tierney & Kraus, 2015) used sample sizes close to our sample size, raising concerns that some if not most of these studies reported inflated estimates of the true population effects under investigation. Of particular concern is that some studies used sample sizes not much larger than Nozaradan et al. (2011) to compare multiple groups (e.g., Celma-Miralles & Toro, 2019; Cirelli et al., 2016; Laffere et al., 2021; Nozaradan et al., 2017; Stupacher et al., 2017) or to test correlations between SSEP amplitude and behavioral or individual difference measures (Cameron et al., 2019; Nguyen et al., 2023; Nozaradan et al., 2016; Tal et al., 2017b; Zalta et al., 2024; Zhao & Kuhl, 2020), even though such effects typically require relatively large sample sizes (Brysbaert, 2019). Relatedly, frequency tagging approaches have become increasingly used by researchers to investigate entrainment to beat-related frequencies in preverbal infants, even though infants are likely to have far noisier data than adult participants (Edalati et al., 2023; Flaten et al., 2022; Lenc et al., 2023; Nguyen et al., 2023). Nevertheless, these studies did not typically test larger numbers of participants to compensate for the extra noise compared to adult studies, which themselves are likely to be underpowered.

This raises concerns of false positives in SSEP studies overall due to the file drawer effect or p-hacking (Masicampo & Lalande, 2012) and is consistent with some studies reporting non-significant effects in key tests (e.g., Celma-Miralles et al., 2016; Cirelli et al., 2016; Flaten et al., 2022; Laffere et al., 2021; Nguyen et al., 2023; Stupacher et al., 2017; Tichko et al., 2022; Zhao & Kuhl, 2020). Many other studies report key effects with p-values between *p* = .05 and *p* = .01 (e.g., Cameron et al., 2019; Edalati et al., 2023; Gibbings et al., 2023; Lenc et al., 2018; Nave et al., 2022; Nozaradan et al., 2012; Tal et al., 2017b; Vanden Bosch der Nederlanden et al., 2020) — sometimes with tests uncorrected for multiple comparisons — another sign of effects that might not replicate (Open Science Collaboration, 2015). However, several studies have been published with findings that are significant at more stringent p-values (e.g., Harding et al., 2019; Lenc et al., 2023; Mathias et al., 2020). Given the mixed evidence from various types of studies, the generally small sample sizes used, and the results of the current replication project, we argue that findings in this literature are not robust enough to support a direct relation between beat-related SSEPs and beat-related human behavior and/or perception, other than the fact that stimulus-driven activity can be recorded reliably at various frequencies that are relevant to musical rhythm and beat perception, and that the stimulus-driven frequency encoding may be predictive of task-related performance. More work is necessary to fully understand the relation between frequency-tagged cortical responses and related human behaviors.

While this study was pre-registered and peer-reviewed as a Registered Report (RR), rather than a Registered Replication Report (RRR), we took care to design the study as a direct replication of Nozaradan et al. (2011). We employed only minor changes to the original methods, such that the data could be interpreted both as a direct replication and with the extended dataset, now including behavioral responses. To consider whether the present findings represent successful replication of the original study Nozaradan et al. (2011), we must primarily consider the pre-registered analyses. Our pre-registered analyses suggest no significant effect of imagery condition; all four meta-analytic effect sizes estimated had confidence intervals that overlapped with 0, indicative of a failed replication. While exploratory analyses suggest a small effect may be present, detecting the effect required a second non-pre-registered analysis, notably with a very large sample size, nearly 20 times as large as the original study (*N*=8 vs *N* =152), and this produced estimated effect sizes that were 4 to 5 times smaller than suggested by the original study. While the use of these metrics is consistent with past replication studies (Errington et al., 2021; Open Science Collaboration, 2015), additional metrics such as holistic subjective evaluations by researchers may also be legitimate and can allow for more nuanced evaluations. In the current study, a reasonable holistic evaluation could be that although the observed effect sizes were much smaller than in the original study and thus require a larger sample to detect, they were in the expected direction, and thus it may be premature to conclude that there is no such effect.

In addition to the need for conducting novel and further replication studies with much larger sample sizes, one may also want to consider additional data analysis techniques that may complement or even outperform frequency-tagging in the efforts to quantify the relationship between neural activity and human beat perception. Such methods could include but are not limited to: 1) investigation of other frequency bands in the neural signal, such as via spectral analyses to estimate beat-related β-band and γ-band modulation (Fujioka et al., 2009; Snyder & Large, 2005); 2) other methods estimating synchronization of the neural signal with the stimulus envelope, such as temporal response functions (TRFs) (David et al., 2007), Phase-Locking Value (PLV) (Tal et al., 2017; Tichko et al., 2022), and canonical correlation analyses (de Cheveigné et al., 2018); 3) investigation of other event-related potentials (which may contribute to the presence of steady state responses), such as contingent negative variation (CNV) (Nobre & van Ede, 2018); 4) multivariate pattern analyses such as multiscale entropy (MSE, a measure of signal irregularity) (Bouwer et al., 2023). In addition, Bayesian statistics would be useful for potentially providing direct evidence to the contrary, (i.e., the absence of a relationship between beat-related SSEPs and beat perception and behavior (Schmalz et al., 2023), should indeed no direct relationship exist.

## 9 Final Disclosures

### 9.1 Author Contributions

Authors of the final paper made a substantive contribution to either planning and writing, creating analyses, materials and methods, or data collection. The authors from the parent lab are listed first, followed by all other authors in alphabetical order.

Karli M. Nave: Conceptualization, Data curation, Formal analysis, Funding Acquisition, Investigation, Methodology, Project administration, Resources, Software, Supervision, Validation, Visualization, Writing – original draft, Writing – review & editing.

Erin E. Hannon: Conceptualization, Funding Acquisition, Methodology, Resources, Supervision, Validation, Writing – original draft, Writing – review & editing.

Joel S. Snyder: Conceptualization, Funding Acquisition, Methodology, Resources, Supervision, Validation, Writing – original draft, Writing – review & editing.

Emma Alexandrov: Investigation.

Sarah C. Allen: Investigation.

Fernando Barbosa: Resources, Software, Supervision, Writing – review & editing.

Mara Breen: Funding Acquisition, Resources, Supervision, Writing – review & editing.

Alexandre Celma-Miralles: Investigation, Writing – review & editing.

Thibault Chabin: Formal analysis, Writing – review & editing.

Joshua R. de Leeuw: Funding Acquisition, Resources, Supervision, Writing – review & editing.

Fernando Ferreira-Santos: Resources, Software, Supervision, Writing – review & editing.

Ahren B. Fitzroy: Investigation, Software, Supervision, Writing – review & editing.

Ningyao Geng: Investigation.

Sean A. Gilmore: Investigation, Writing – review & editing.

Reyna L. Gordon: Supervision, Writing – review & editing.

Jessica A. Grahn: Funding Acquisition, Resources, Supervision, Writing – review & editing.

Paz Har-shai Yahav: Investigation, Writing – review & editing.

Kiara Holm: Investigation.

Falk Huettig: Funding Acquisition, Supervision, Writing – review & editing.

Anne Keitel: Investigation, Supervision, Writing – review & editing.

Christian Keitel: Resources, Software, Supervision, Writing – review & editing.

Eniko Ladanyi: Investigation, Software, Supervision, Writing – review & editing.

Yiguang Liu: Investigation, Writing – review & editing.

Psyche Loui: Investigation, Funding Acquisition, Resources, Supervision, Writing – review & editing.

Grace A. Lourie: Investigation, Resources, Writing – review & editing.

Inês Macedo: Investigation, Software, Writing – review & editing.

Cyrille L. Magne: Resources, Software, Supervision, Writing – review & editing.

Cecilie Møller: Investigation, Funding Acquisition, Writing – review & editing.

David Moreau: Investigation, Resources, Supervision, Writing – review & editing.

Srishti Nayak: Investigation, Software, Supervision, Writing – review & editing.

Markus Ostarek: Supervision, Writing – review & editing.

Dingyi Pan: Investigation.

Rita Pasion: Investigation, Resources, Software, Writing – review & editing.

Mariana R. Pereira: Resources, Writing – review & editing.

Danna S. Pinto: Investigation, Writing – review & editing.

Eva D. Poort: Investigation, Writing – review & editing.

Meg Renzelman: Investigation.

Frank A. Russo: Resources, Writing – review & editing.

Jan Stupacher: Investigation, Funding Acquisition, Writing – review & editing.

Parker Tichko: Investigation, Resources, Software, Supervision, Writing – review & editing.

Lucy Wight: Investigation, Writing – review & editing.

Chu Yi Yu: Investigation, Writing – review & editing.

Elana M. Zion Golumbic: Funding Acquisition, Resources, Supervision, Writing – review & editing.

## 9.3 Acknowledgements

We thank Sylvie Nozaradan for providing the stimulus used in the original study. We thank the researchers from participating labs who read and suggested changes to the manuscript and analysis plan before the pre-data manuscript was submitted. In addition, we would like to thank the following collaborators for their contributions and feedback on the pre-data manuscript: Laura Cirelli, Shannon Heald, Molly Henry, Howard Nusbaum, Adam Tierney, Laurel Trainor, and Ted Zanto. Finally, we would like to thank those who expressed their support by signing up to contribute a dataset but were unable to complete their contribution due to the COVID-19 pandemic or other obstacles, which includes the following colleagues: Ayse Akgoz, Eleonora Beier, Laura Cirelli, Fernanda Ferreira, Asil Özdoğru, Günay Sağir, Tamara Swaab, and Laurel Trainor.

## 9.5 Supplemental Materials

The OSF project page can be found here: https://osf.io/rpvde. Pre-Registered Materials and the Final Study Data and Materials can be accessed at https://osf.io/rpvde/files/osfstorage. Raw EEG data are available on Harvard Dataverse and can be accessed at: https://doi.org/10.7910/DVN/ELZFWK

## 9.6 Prior Versions

The Stage 1 provisionally-accepted manuscript can be found on our OSF page at: https://osf.io/b2pvz.

## Supplementary Figures

**Figure S1.**
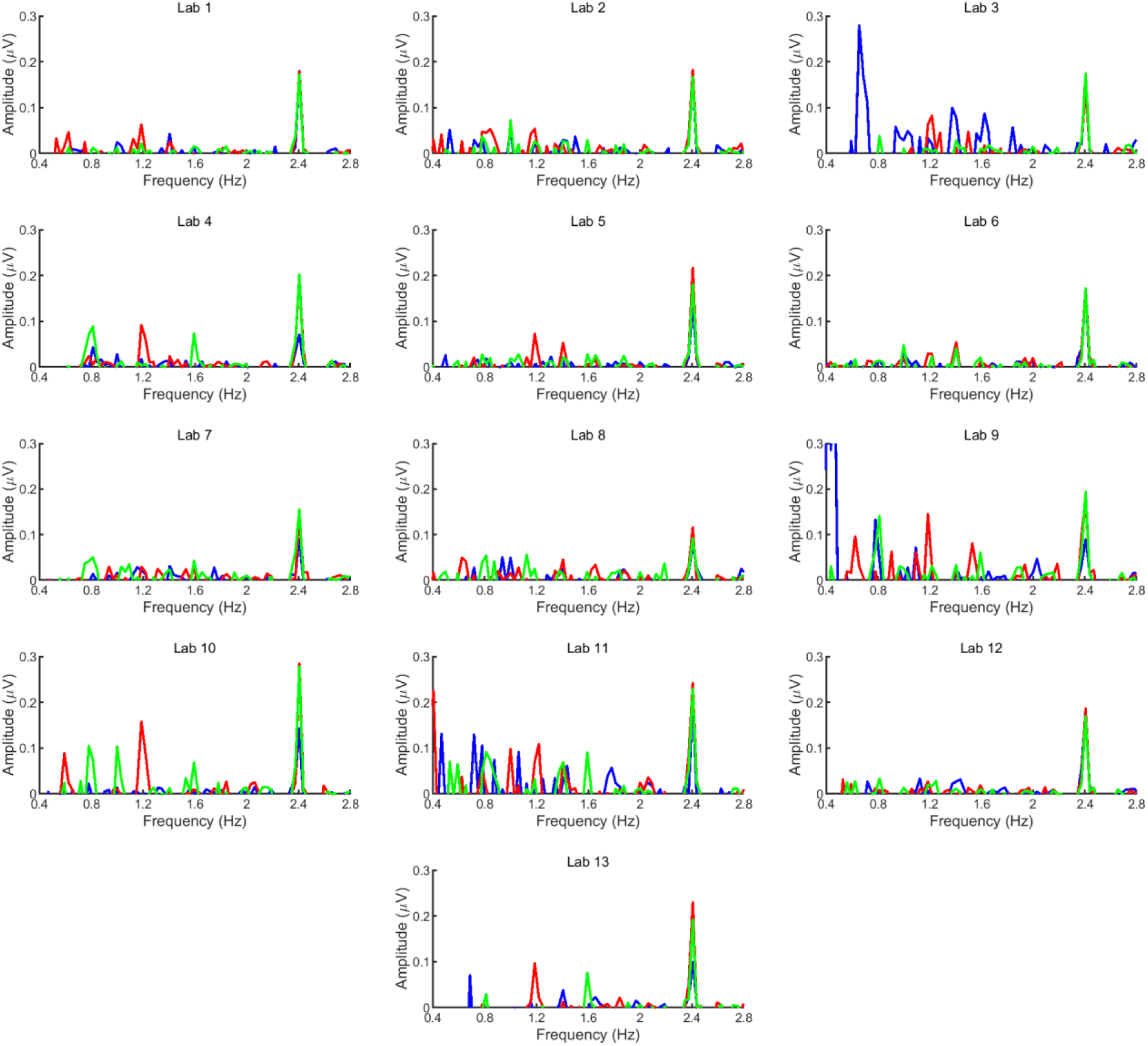
Averages of the stimulus- and beat-related steady-state EPs elicited by the 2.4 Hz auditory beat in the control condition (blue), the binary beat imagery condition (red), and the ternary beat imagery condition (green) for all participating labs. The frequency spectra represent the SNR-corrected amplitude of the EEG signal (in microvolts) as a function of frequency, averaged across all scalp electrodes, after applying the noise subtraction procedure.

**Figure S2.**
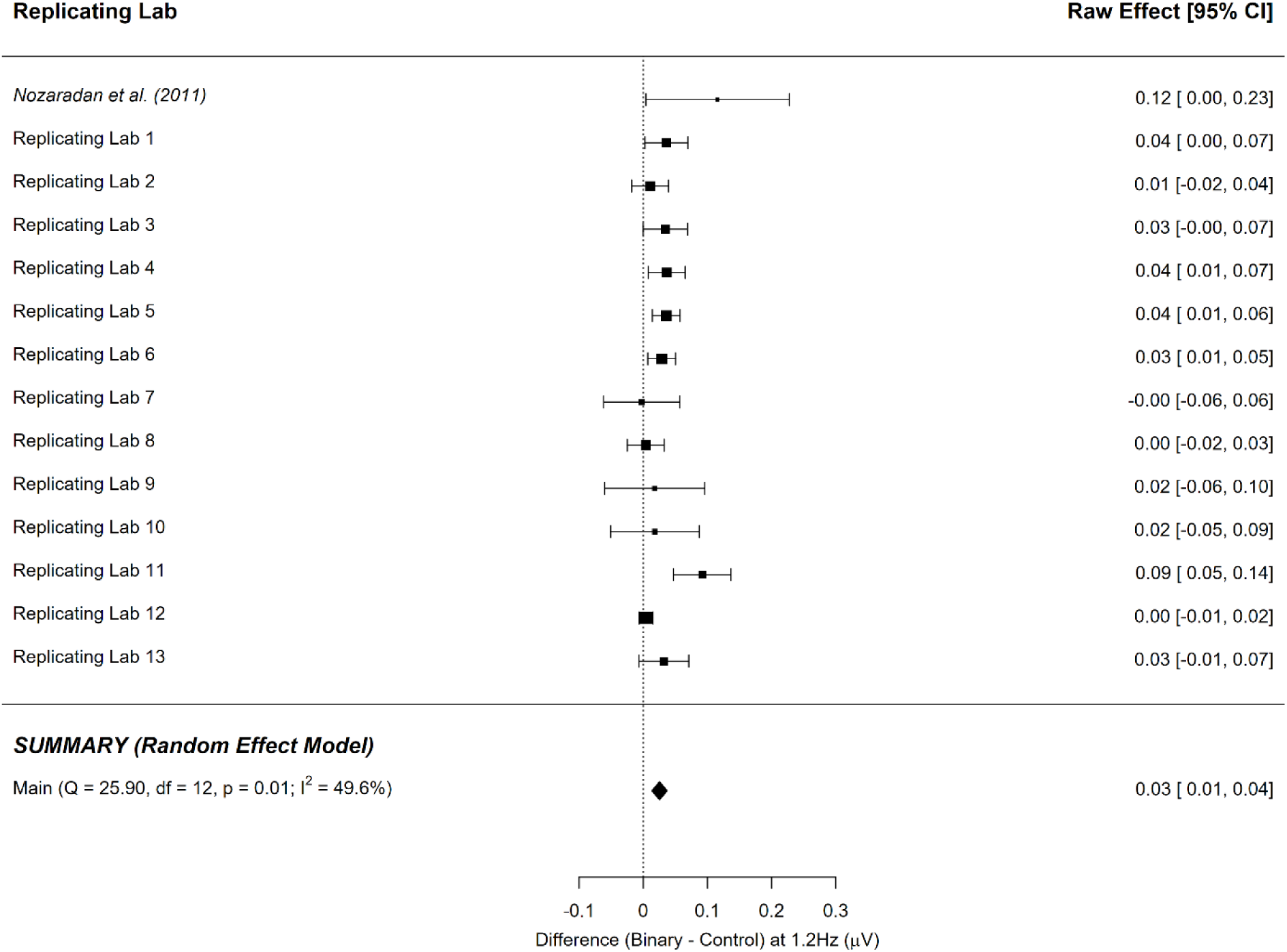
Point-estimate for the meta-analytic effect of condition (control task vs. binary imagery) on the binary frequency using medians. Squares indicate each lab’s median difference, where the size of the square corresponds to the inverse of the standard error of the difference score, and the error bars indicate 95% confidence intervals (CI) around the median difference. Diamonds indicate the random-effects meta-analytic effect size estimate, where the width represents the 95% CI. The diamond represents the meta-analytic estimate (Note: this estimate does not include the original Nozaradan et al. (2011) result.

**Figure S3.**
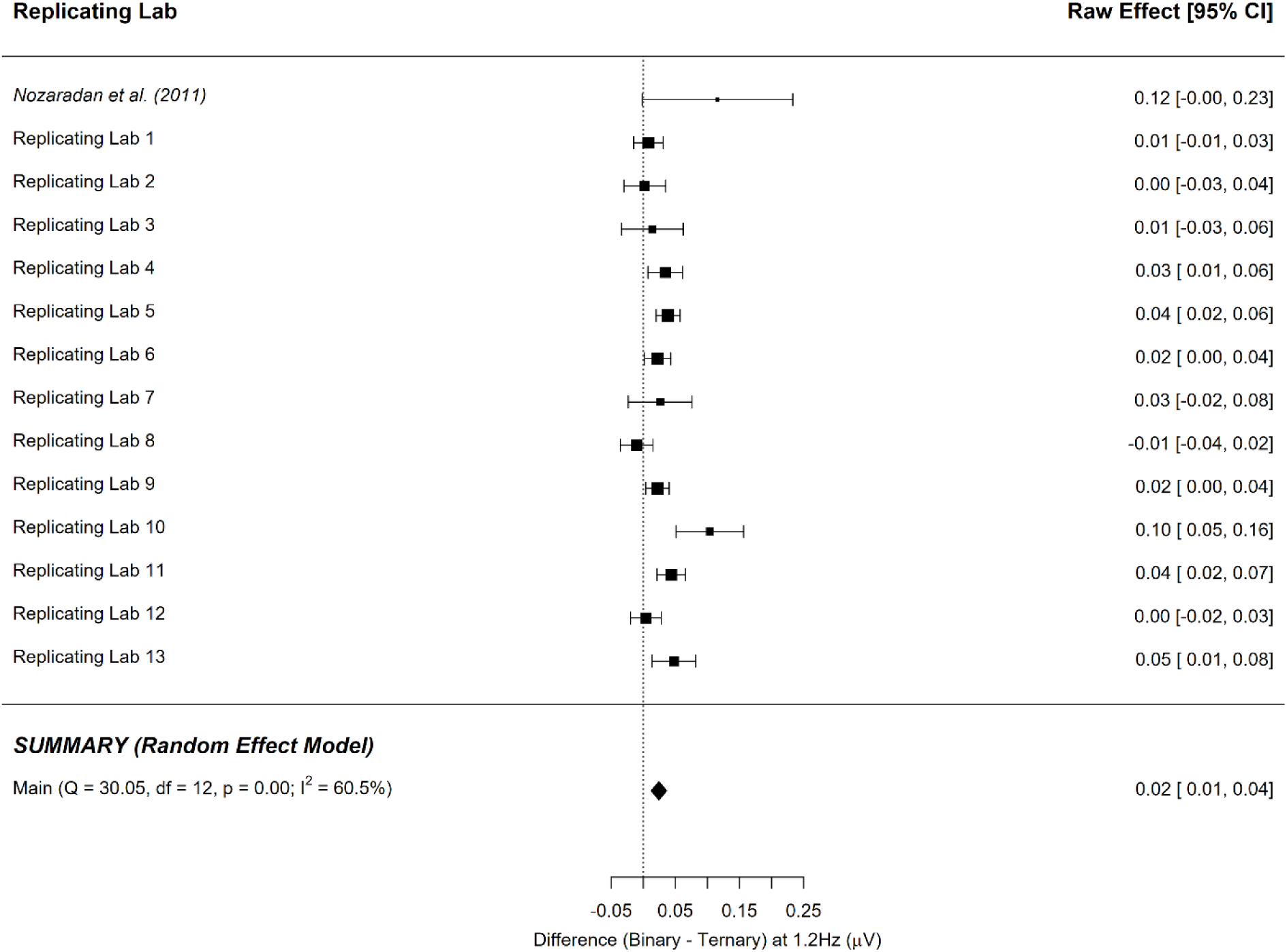
Point-estimate for the meta-analytic effect of condition (binary imagery vs. ternary imagery) on the binary frequency using medians. Squares indicate each lab’s median difference, where the size of the square corresponds to the inverse of the standard error of the difference score, and the error bars indicate 95% confidence intervals (CI) around the median difference. Diamonds indicate the random-effects meta-analytic effect size estimate, where the width represents the 95% CI. The diamond represents the meta-analytic estimate (Note: this estimate does not include the original Nozaradan et al. (2011) result.

**Figure S4.**
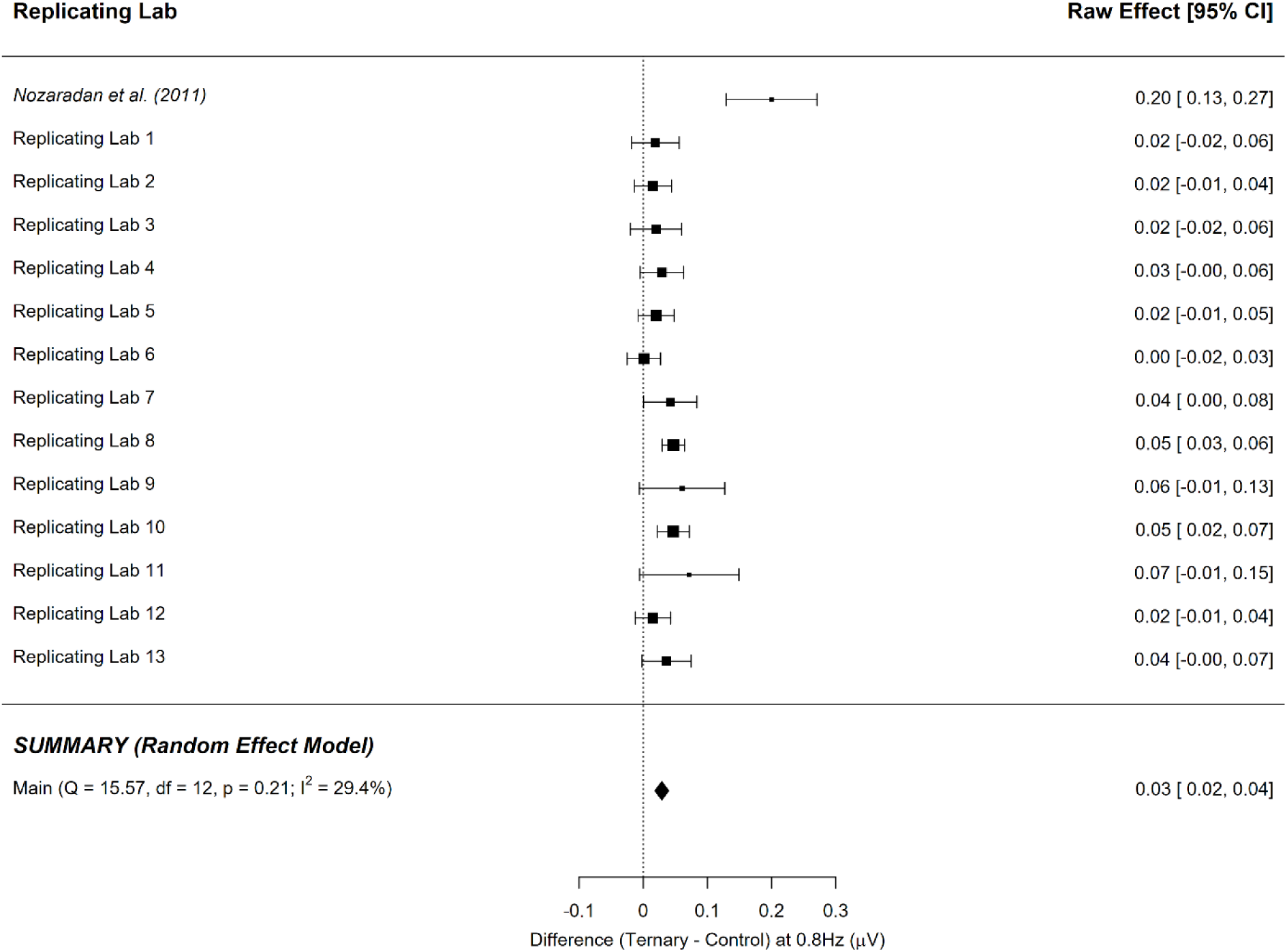
Point-estimate for the meta-analytic effect of condition (control task vs. ternary imagery) on the ternary frequency using medians. Squares indicate each lab’s median difference, where the size of the square corresponds to the inverse of the standard error of the difference score, and the error bars indicate 95% confidence intervals (CI) around the median difference. Diamonds indicate the random-effects meta-analytic effect size estimate, where the width represents the 95% CI. The diamond represents the meta-analytic estimate (Note: this estimate does not include the original Nozaradan et al. (2011) result.

**Figure S5.**
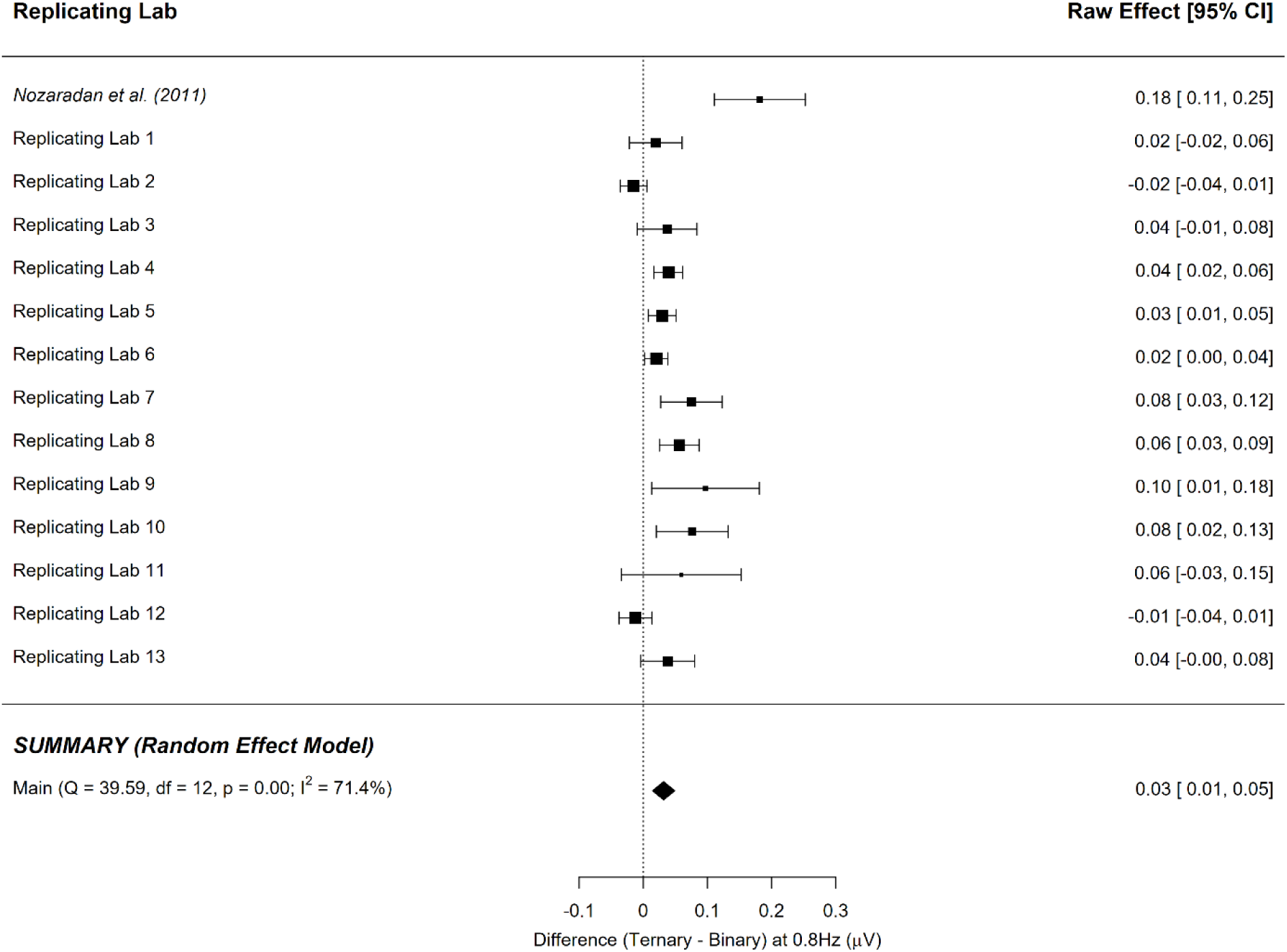
Point-estimate for the meta-analytic effect of condition (ternary imagery vs. binary imagery) on the ternary frequency using medians. Squares indicate each lab’s median difference, where the size of the square corresponds to the inverse of the standard error of the difference score, and the error bars indicate 95% confidence intervals (CI) around the median difference. Diamonds indicate the random-effects meta-analytic effect size estimate, where the width represents the 95% CI. The diamond represents the meta-analytic estimate (Note: this estimate does not include the original Nozaradan et al. (2011) result.

**Figure S6.**
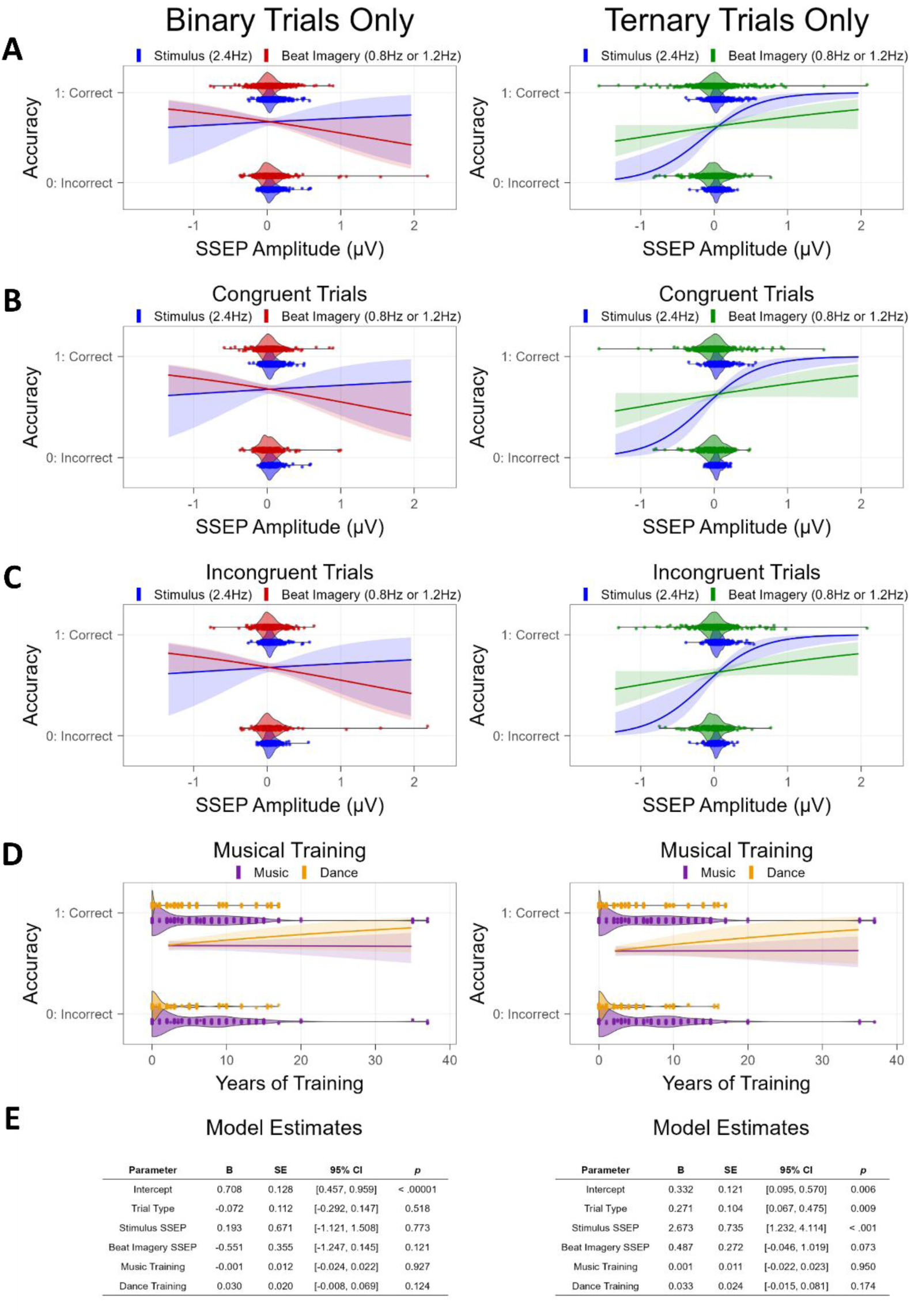
Results of trial-by-trial logistic regressions conducted separately for binary imagery trials (left) and ternary imagery trials (right), examining whether frequency-tagged neural responses, musical training, and/or study factors predict performance on the imagery-related probe task. Raw values of continuous predictors are plotted along the x-axis, and accuracy on the probe task is plotted on the y-axis (0: incorrect response, 1: correct response). SSEP predictors represent SNR-corrected amplitude at the respective frequency in the EEG signal. A) Model predictions on all trials for SSEP predictors. B) Model predictions on congruent (ON beat) trials for SSEP predictors. C) Model predictions on incongruent (OFF beat) trials for SSEP predictors. D) Model predictions on all trials for musical training predictors (Music, Dance). E) Resulting parameter estimates from trial-by-trial logistic regression.

---

**Table S1.**
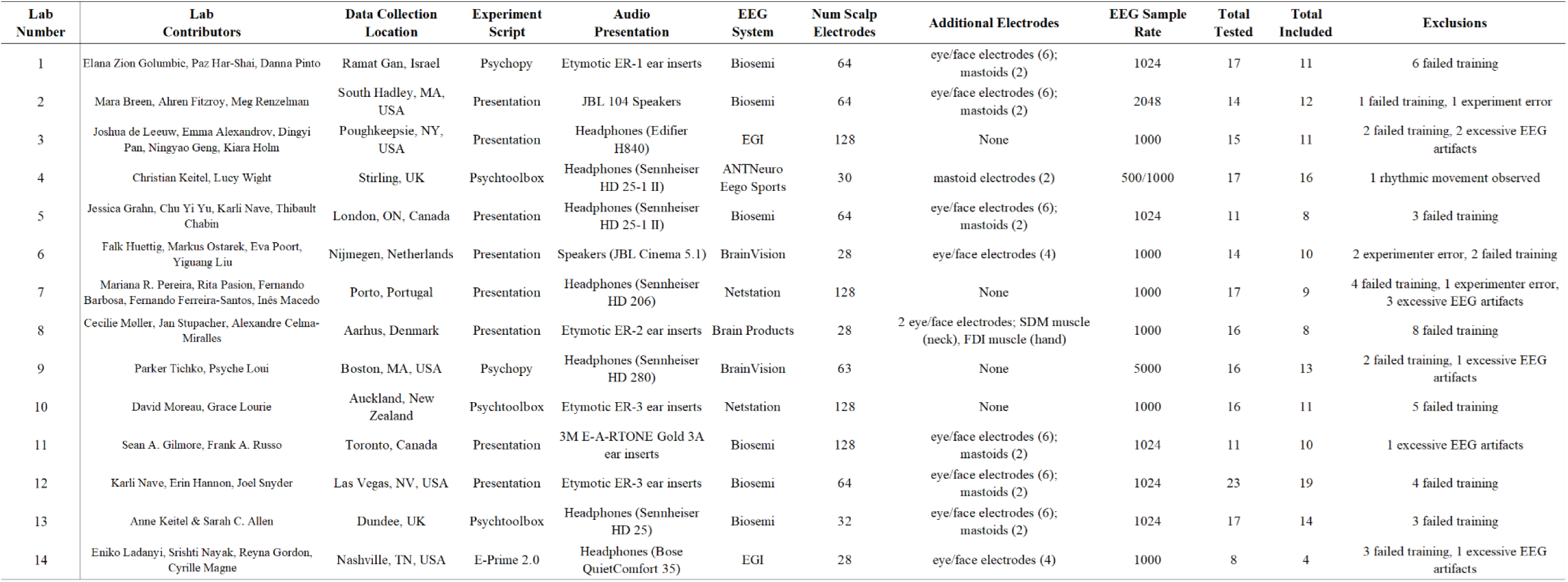
Additional details for all contributing labs.

**Table S2.** Results from one-tailed t-test (test value: 0) to investigate whether the SNR-corrected amplitude of the SSEPs are significantly greater than 0 µV.

| Lab Number | One-Sample t-tests ( One-Tailed: Amplitude > 0 ) |  |  |  |  |  |  |  |  |  |  |  |
| --- | --- | --- | --- | --- | --- | --- | --- | --- | --- | --- | --- | --- |
|  | 0.8 Hz<br>(Ternary Frequency) |  |  | 1.2 Hz<br>(Binary Frequency) |  |  | 1.6 Hz<br>(1st Ternary Harmonic) |  |  | 2.4 Hz<br>(Stimulus Frequency) |  |  |
|  | Control | Binary | Ternary | Control | Binary | Ternary | Control | Binary | Ternary | Control | Binary | Ternary |
| 0 | $t(7) = 0.1$<br>$p = 0.91$ | $t(7) = 1.4$<br>$p = 0.21$ | $t(7) = 5.7$<br>$p = 0.001$ | $t(7) = -0.2$<br>$p = 0.81$ | $t(7) = -3.1$<br>$p = 0.01$ | $t(7) = -0.1$<br>$p = 0.92$ | $t(7) = -0.3$<br>$p = 0.74$ | $t(7) = -0.4$<br>$p = 0.71$ | $t(7) = 26.8$<br>$p < .0001$ | $t(7) = -8.1$<br>$p < 0.0001$ | $t(7) = -4.3$<br>$p = 0.003$ | $t(7) = -4.4$<br>$p = 0.03$ |
| 1 | $t(10) = -2.492$<br>$p = 0.984$ | $t(10) = -1.983$<br>$p = 0.962$ | $t(10) = 0.201$<br>$p = 0.422$ | $t(10) = -1.115$<br>$p = 0.880$ | $t(10) = -2.081$<br>$p = 0.032$ | $t(10) = 1.110$<br>$p = 0.147$ | $t(10) = -0.129$<br>$p = 0.550$ | $t(10) = -0.814$<br>$p = 0.783$ | $t(10) = 2.107$<br>$p = 0.031$ | $t(10) = -1.115$<br>$p < .001$ | $t(10) = 3.497$<br>$p = 0.003$ | $t(10) = 6.568$<br>$p < .001$ |
| 2 | $t(11) = -0.280$<br>$p = 0.608$ | $t(11) = 2.177$<br>$p = 0.026$ | $t(11) = 1.322$<br>$p = 0.107$ | $t(11) = 0.556$<br>$p = 0.295$ | $t(11) = 2.066$<br>$p = 0.032$ | $t(11) = 0.662$<br>$p = 0.261$ | $t(11) = -2.354$<br>$p = 0.981$ | $t(11) = -2.033$<br>$p = 0.967$ | $t(11) = -0.038$<br>$p = 0.515$ | $t(11) = -7.035$<br>$p < .001$ | $t(11) = 5.957$<br>$p < .001$ | $t(11) = 4.384$<br>$p < .001$ |
| 3 | $t(11) = -0.975$<br>$p = 0.824$ | $t(11) = -1.195$<br>$p = 0.870$ | $t(11) = -0.314$<br>$p = 0.620$ | $t(11) = -0.139$<br>$p = 0.554$ | $t(11) = 1.875$<br>$p = 0.045$ | $t(11) = 0.865$<br>$p = 0.204$ | $t(11) = 1.016$<br>$p = 0.167$ | $t(11) = -0.576$<br>$p = 0.711$ | $t(11) = -0.067$<br>$p = 0.526$ | $t(11) = 5.549$<br>$p < .001$ | $t(11) = 3.531$<br>$p = 0.003$ | $t(11) = 5.846$<br>$p < .001$ |
| 4 | $t(15) = 2.911$<br>$p = 0.005$ | $t(15) = 2.019$<br>$p = 0.031$ | $t(15) = 3.434$<br>$p = 0.002$ | $t(15) = -0.268$<br>$p = 0.604$ | $t(15) = 2.956$<br>$p = 0.005$ | $t(15) = 1.062$<br>$p = 0.153$ | $t(15) = -0.109$<br>$p = 0.543$ | $t(15) = -3.491$<br>$p = 0.998$ | $t(15) = 1.356$<br>$p = 0.098$ | $t(15) = 4.070$<br>$p < .001$ | $t(15) = 4.536$<br>$p < .001$ | $t(15) = 3.120$<br>$p = 0.004$ |
| 5 | $t(7) = -0.045$<br>$p = 0.517$ | $t(7) = -1.171$<br>$p = 0.860$ | $t(7) = 2.316$<br>$p = 0.027$ | $t(7) = -1.256$<br>$p = 0.875$ | $t(7) = 3.030$<br>$p = 0.010$ | $t(7) = -2.241$<br>$p = 0.970$ | $t(7) = -1.628$<br>$p = 0.926$ | $t(7) = -0.442$<br>$p = 0.664$ | $t(7) = 0.602$<br>$p = 0.283$ | $t(7) = 6.674$<br>$p < .001$ | $t(7) = 6.671$<br>$p < .001$ | $t(7) = 5.298$<br>$p < .001$ |
| 6 | $t(9) = 0.654$<br>$p = 0.265$ | $t(9) = -1.217$<br>$p = 0.873$ | $t(9) = 1.370$<br>$p = 0.102$ | $t(9) = -2.054$<br>$p = 0.965$ | $t(9) = 2.899$<br>$p = 0.009$ | $t(9) = 0.227$<br>$p = 0.413$ | $t(9) = -0.924$<br>$p = 0.810$ | $t(9) = 0.299$<br>$p = 0.386$ | $t(9) = 1.821$<br>$p = 0.051$ | $t(9) = 4.828$<br>$p < .001$ | $t(9) = 4.030$<br>$p = 0.001$ | $t(9) = 5.678$<br>$p < .001$ |
| 7 | $t(8) = 0.196$<br>$p = 0.425$ | $t(8) = -1.607$<br>$p = 0.927$ | $t(8) = 2.796$<br>$p = 0.012$ | $t(8) = 1.805$<br>$p = 0.054$ | $t(8) = 0.639$<br>$p = 0.270$ | $t(8) = -0.492$<br>$p = 0.682$ | $t(8) = -1.721$<br>$p = 0.938$ | $t(8) = 0.885$<br>$p = 0.201$ | $t(8) = 1.343$<br>$p = 0.108$ | $t(8) = 2.756$<br>$p = 0.012$ | $t(8) = 3.055$<br>$p = 0.008$ | $t(8) = 3.968$<br>$p = 0.002$ |
| 8 | $t(7) = -1.565$<br>$p = 0.919$ | $t(7) = -2.041$<br>$p = 0.960$ | $t(7) = 2.442$<br>$p = 0.022$ | $t(7) = -1.014$<br>$p = 0.828$ | $t(7) = -0.688$<br>$p = 0.743$ | $t(7) = 0.525$<br>$p = 0.308$ | $t(7) = -0.631$<br>$p = 0.726$ | $t(7) = 0.065$<br>$p = 0.475$ | $t(7) = 0.163$<br>$p = 0.437$ | $t(7) = 3.260$<br>$p = 0.007$ | $t(7) = 2.995$<br>$p = 0.010$ | $t(7) = 3.591$<br>$p = 0.004$ |
| 9 | $t(12) = 0.864$<br>$p = 0.202$ | $t(12) = -0.769$<br>$p = 0.772$ | $t(12) = 3.297$<br>$p = 0.003$ | $t(12) = -1.269$<br>$p = 0.886$ | $t(12) = 2.422$<br>$p = 0.016$ | $t(12) = 0.911$<br>$p = 0.190$ | $t(12) = -0.643$<br>$p = 0.734$ | $t(12) = -0.576$<br>$p = 0.712$ | $t(12) = 1.035$<br>$p = 0.161$ | $t(12) = 2.322$<br>$p = 0.019$ | $t(12) = 4.728$<br>$p < .001$ | $t(12) = 3.697$<br>$p = 0.002$ |
| 10 | $t(10) = -0.209$<br>$p = 0.581$ | $t(10) = -1.943$<br>$p = 0.960$ | $t(10) = 2.272$<br>$p = 0.023$ | $t(10) = 0.056$<br>$p = 0.478$ | $t(10) = 1.755$<br>$p = 0.055$ | $t(10) = -2.213$<br>$p = 0.974$ | $t(10) = 0.996$<br>$p = 0.178$ | $t(10) = 0.791$<br>$p = 0.224$ | $t(10) = -1.206$<br>$p = 0.128$ | $t(10) = 4.759$<br>$p < .001$ | $t(10) = 5.872$<br>$p < .001$ | $t(10) = 5.842$<br>$p < .001$ |
| 11 | $t(9) = -1.630$<br>$p = 0.931$ | $t(9) = 0.477$<br>$p = 0.322$ | $t(9) = 1.941$<br>$p = 0.042$ | $t(9) = -2.685$<br>$p = 0.987$ | $t(9) = 3.608$<br>$p = 0.003$ | $t(9) = -0.339$<br>$p = 0.629$ | $t(9) = -1.367$<br>$p = 0.898$ | $t(9) = -3.038$<br>$p = 0.993$ | $t(9) = 1.045$<br>$p = 0.162$ | $t(9) = 2.739$<br>$p = 0.011$ | $t(9) = 4.864$<br>$p < .001$ | $t(9) = 8.585$<br>$p < .001$ |
| 12 | $t(18) = -0.037$<br>$p = 0.515$ | $t(18) = 0.585$<br>$p = 0.283$ | $t(18) = 0.822$<br>$p = 0.211$ | $t(18) = -0.374$<br>$p = 0.644$ | $t(18) = 1.644$<br>$p = 0.059$ | $t(18) = 1.695$<br>$p = 0.054$ | $t(18) = -1.506$<br>$p = 0.925$ | $t(18) = 0.830$<br>$p = 0.209$ | $t(18) = 0.987$<br>$p = 0.168$ | $t(18) = 8.669$<br>$p < .001$ | $t(18) = 8.293$<br>$p < .001$ | $t(18) = 7.591$<br>$p < .001$ |
| 13 | $t(13) = -5.493$<br>$p = 1.000$ | $t(13) = -1.321$<br>$p = 0.895$ | $t(13) = -0.368$<br>$p = 0.641$ | $t(13) = -0.767$<br>$p = 0.772$ | $t(13) = 2.888$<br>$p = 0.006$ | $t(13) = -2.641$<br>$p = 0.990$ | $t(13) = -1.362$<br>$p = 0.902$ | $t(13) = -1.537$<br>$p = 0.926$ | $t(13) = 2.328$<br>$p = 0.018$ | $t(13) = 3.616$<br>$p = 0.002$ | $t(13) = 4.454$<br>$p < .001$ | $t(13) = 3.862$<br>$p < .001$ |
| Lab 1:13 * | $t(151) = -1.082$<br>$p = 0.859$ | $t(151) = -1.085$<br>$p = 0.860$ | $t(151) = 5.195$<br>$p < .001$ | $t(151) = -2.048$<br>$p = 0.979$ | $t(151) = 6.310$<br>$p < .001$ | $t(151) = 0.522$<br>$p = 0.301$ | $t(151) = -0.843$<br>$p = 0.800$ | $t(151) = -2.055$<br>$p = 0.979$ | $t(151) = 3.461$<br>$p < .001$ | $t(151) = 14.943$<br>$p < .001$ | $t(151) = 16.342$<br>$p < .001$ | $t(151) = 15.312$<br>$p < .001$ |

## Appendix A. Individual Lab Details

See below for each lab’s list of contributing authors, funding acknowledgements, and pre-registration details^8^. Pre-registration details reflect each lab’s committed pre-registration plan prior to collecting data.

### Lab 1: braindynlab-biu

**Contributing Authors:** Paz Har-Shai Yahav¹, Danna Pinto¹, Elana M. Zion Golumbic¹

¹ The Gonda Center for Multidisciplinary Brain Research, Bar Ilan University, Ramat Gan, Israel

**Funding Acknowledgements:** Israel Science Foundation Grant (ISF #2339/20)

**Pre-Registration**

#### I. Participants

- **Stopping Criteria:** When we reach our sample size commitment
- **Sample Size Commitment:** Full block of 16, and then will stop
- **Compensation:** Course credit (1 credit per hour)
- **Normal Hearing:** Verified via self-report

#### II. Experiment Set-Up

- **Testing Language:** Hebrew* (*translated by P. Har-Shai*)
- **Experiment Room:** Sound-attenuated booth (single-walled)
- **Location of Training Experimenter During Experiment:** In the same room/booth as the participant during the entire study
- **Location of Data Experimenter During Experiment:** In a separate area/room outside of the testing booth during the entire study
- **Experiment Script to be Used:** Psychopy script (written by P. Tichko)

#### III. Audio Delivery During Experiment

- **Training Experimenter:** Sennheiser 280 Pro headphones (experimenter removed these during test trials)
- **Participant:** Ear inserts (Etymotic ER-1; dB levels not measured)

#### IV. Changes in Pre-registration Due to COVID-19 Pandemic

- **Data Collection Completed:** February 20, 2020

**The lab’s pre-registration was followed in all other respects.**

### Lab 2: capslab-mhc

**Contributing Authors:** Mara Breen¹, Ahren B. Fitzroy¹, Meg Renzelman¹ ¹ Mount Holyoke College, South Hadley, MA, United States

**Funding Acknowledgements:** James S. McDonnell Foundation Scholar Award in Understanding Human Cognition

**Pre-Registration**

#### I. Participants

- **Stopping Criteria:** When the official time for data collection ends (i.e., will continue testing as many participants as possible until data collection closes or the specified alternative stopping date)
- **Sample Size Commitment:** Full block of 16, but will continue collecting until the end date for data collection
- **Compensation:** Course credit (1 credit)
- **Normal Hearing:** Hearing tests conducted using an audiometer

#### II. Experiment Set-Up

- **Testing Language:** English
- **Experiment Room:** Room with sound-proofing materials (e.g., sound-attenuating walls, curtains, etc.)
- **Location of Training Experimenter During Experiment:** In the same room/booth as the participant to ensure compliance
- **Location of Data Experimenter During Experiment:** In a separate control room or outside the testing booth
- **Experiment Script to be Used:** Presentation script

#### III. Audio Delivery During Experiment

- **Training Experimenter:** Speakers (listened to masking music through headphones during test trials)
- **Participant:** Ear inserts (Etymotic ER-3; dB levels not measured)

#### IV. Changes in Pre-registration Due to COVID-19 Pandemic

- **Data Collection Completed:** March 6, 2020
- The lab’s pre-registration was followed in all other respects.

### Lab 3: cogsci-vassar

**Contributing Authors:** Emma Alexandrov¹, Joshua R. de Leeuw¹, Ningyao Geng¹, Kiara Holm¹, Dingyi Pan¹

¹ Department of Cognitive Science, Vassar College, Poughkeepsie, NY, United States

**Funding Acknowledgements:** None

**Pre-Registration**

#### I. Participants

- **Stopping Criteria:** When the specific number of participants listed below is tested, accounting for exclusions prior to data review
- **Sample Size Commitment:** Full block of 16, and will stop after reaching 16
- **Compensation:** Not specified
- **Normal Hearing:** Verified via self-report

#### II. Experiment Set-Up

- Testing Language: English
- Experiment Room: Quiet location in the building
- Location of Training Experimenter During Experiment: In the same room/booth as the participant to ensure compliance
- Location of Data Experimenter During Experiment: In a separate control room or outside the testing booth
- Experiment Script to be Used: Presentation script

#### III. Audio Delivery During Experiment

- **Training Experimenter:** Headphones (Edifier H840; experimenter removed these during test trials)
- **Participant:** Headphones (Edifier H840; dB levels not measured)

#### IV. Changes in Pre-registration Due to COVID-19 Pandemic

- **Data Collection Completed:** November 23 (year unspecified)
- **Compensation:** Participants were paid $10/hour

**The lab’s pre-registration was followed in all other respects.**

### Lab 4: dvplab-ustir

**Contributing Authors:** Christian Keitel¹, Lucy Wight²

¹ Department of Psychology, University of Dundee, Dundee, UK ² Department of Psychology, University of Stirling, Stirling, UK

**Funding Acknowledgements:** RSE Saltire Facilitation Network Award (Reference Number 1963)

**Pre-Registration**

#### I. Participants

- **Stopping Criteria:** When the specific number of participants listed below is tested, accounting for exclusions prior to data review
- **Sample Size Commitment:** Full block of 16, and will stop after reaching 16
- **Compensation:** Financial compensation (£7.50/hour) or course credit (2 “tokens”)
- **Normal Hearing:** Verified via self-report only

#### II. Experiment Set-Up

- **Testing Language:** English
- **Experiment Room:** Room with occasional mild/moderate noise from adjacent rooms
- **Location of Training Experimenter During Experiment: In the same room/booth as the participant to ensure compliance**
- **Location of Data Experimenter During Experiment: In a separate control room or outside the testing booth**
- **Experiment Script to be Used:** Psychtoolbox

#### III. Audio Delivery During Experiment

- **Training Experimenter:** Headphones (Philips FX3BK/00; experimenter removed these during test trials)
- **Participant:** Headphones (Sennheiser HD 25-1 II; dB levels measured: 37 dB in silence, 40.5 dB to sound)

#### IV. Changes in Pre-registration Due to COVID-19 Pandemic

- **Data Collection Completed:** December 2, 2021
- **Location of Training Experimenter During Experiment:** Due to COVID-19 guidance, the training experimenter only stayed for the practice and monitored compliance through a window.

**The lab’s pre-registration was followed in all other respects.**

### Lab 5: grahnlab

**Contributing Authors:** Thibault Chabin^1^, Jessica A. Grahn^1^, Karli M. Nave^1,2^, Chu Yi Yu^1^

¹ Department of Psychology, Centre for Brain and Mind, University of Western Ontario, London, ON, Canada

^2^ Department of Psychology, University of Nevada Las Vegas, Las Vegas, NV, United States

**Funding Acknowledgements:** Discovery Grant and Steacie Award from NSERC to JAG, NSERC-CREATE Complex Dynamics Fellowship (to KMN)

**Pre-Registration**

#### I. Participants

- **Stopping Criteria: When the specific number of participants listed below is tested, accounting for exclusions prior to data review**
- **Sample Size Commitment: Full block of 16, and will stop after reaching 16**
- **Compensation: $10/hour or 1 credit/hour or no financial compensation with course credit**
- **Normal Hearing:** Verified via self-report only

#### II. Experiment Set-Up

- **Testing Language: English**
- **Experiment Room: Sound-“proof” booth (single-walled)**
- **Location of Training Experimenter During Experiment: In the same room/booth as the participant to ensure compliance**
- **Location of Data Experimenter During Experiment: In a separate control room or outside the testing booth**
- **Exper**iment Script to be Used: **MATLAB**

#### III. Audio Delivery During Experiment

- **Training Experimenter: Headphones (Sennheiser HD 25-1 II; experimenter removed these during test trials)**
- **Participant:** Headphones (Sennheiser HD 25-1 II; dB levels measured: 30.6 dB in silence, 84.6 dB to sound using standardized testing)

#### IV. Changes in Pre-registration Due to COVID-19 Pandemic

- **Data Collection Completed: April 29, 2022**
- **Compensation: All participants were paid $10/hour**
- **Experiment Script to be Used:** Presentation

**The lab’s pre-registration was followed in all other respects.**

### Lab 6: huettig-mpi

**Contributing Authors:** Falk Huettig¹, Yiguang Liu², Markus Ostarek¹, Eva D. Poort¹

¹ Max Planck Institute for Psycholinguistics, Nijmegen, The Netherlands

² School of International Studies, Zhejiang University, Hangzhou, China

**Funding Acknowledgements:** None

**Pre-Registration**

#### I. Participants

- **Stopping Criteria: When the specific number of participants listed below is tested, accounting for exclusions prior to data review**
- **Sample Size Commitment: Full block of 16, and will stop after reaching 16**
- **Compensation: Not specified**
- **Normal Hearing:** Verified via self-report only

#### II. Experiment Set-Up

- **Testing Language: English**
- **Experiment Room: Sound-“proof” booth (single-walled)**
- **Location of Training Experimenter During Experiment: In the same room/booth as the participant to ensure compliance**
- **Location of Data Experimenter During Experiment: In a separate control room or outside the testing booth**
- **Experiment Script to be Used:** Presentation

#### III. Audio Delivery During Experiment

- **Training Experimenter: Speakers (JBL Cinema 5.1; experimenter listened to masking music through headphones during testing)**
- **Participant:** Speakers (JBL Cinema 5.1; dB levels not measured)

#### IV. Changes in Pre-registration Due to COVID-19 Pandemic

- **Data Collection Completed: October 14, 2020**
- **Compensation:** Participants were paid €10/hour

**The lab’s pre-registration was followed in all other respects.**

### Lab 7: labnpf-up

**Contributing Authors:** Fernando Barbosa¹, Fernando Ferreira-Santos², Rita Pasion², Mariana R. Pereira¹, Inês Macedo¹

¹ Laboratory of Neuropsychophysiology, Faculty of Psychology and Education Sciences,

University of Porto, Porto, Portugal

² HEI-Lab: Digital Human-Environment Interaction Labs, Universidade Lusófona, Porto, Portugal

**Funding Acknowledgements:** FUNDAÇÃO PARA A CIÊNCIA E TECNOLOGIA (FCT), UNDER HEI-LAB R&D UNIT (UIDB/05380/2020)

**Pre-Registration**

#### I. Participants

- **Stopping Criteria: When I have tested the specific number of participants listed below, accounting for exclusions prior to looking at the data.**
- **Sample Size Commitment: Full block of 16, and we will stop after we reach 16.**
- **Compensation: Financial compensation.**
- **Normal Hearing:** Relied on self-report only.

#### II. Experiment Set-Up

- **Testing Language: Portuguese (translated by ?).**
- **Experiment Room: Room with sound-proofing materials (e.g., sound attenuating walls, curtains, etc.).**
- **Location of Training Experimenter During Experiment: In the same room/booth as the participant, and they will remain there throughout the experiment to ensure participant compliance with instructions.**
- **Location of Data Experimenter During Experiment: In a separate control room or outside the testing booth for the entire experiment.**
- **Experiment Script to be Used:** Presentation.

#### III. Audio Delivery During Experiment

- **Training Experimenter: Headphones (Sennheiser HD 206; experimenter removed these during test trials).**
- **Participant:** Headphones (Sennheiser HD 206; dB were measured; 35-47 dB in silence).

#### IV. Changes in Pre-registration Due to COVID Pandemic

- **Data collection completed: N/A.**
- **Compensation: Participants were compensated with a 10€ shopping voucher.**
- **dB in Silence: 32.1 dB was measured (October 8th, 2021).**
- **dB to Sound:** Record of measurement missing.

**The lab’s pre-registration was followed in all other respects.**

### Lab 8: mib-au

**Contributing Authors:** Alexandre Celma-Miralles^1^, Cecilie Møller^1^, Jan Stupacher^1^

¹ Department of Clinical Medicine – Center for Music in the Brain, Aarhus University & The Royal Academy of Music, Aarhus, Denmark

**Funding Acknowledgements:** CM and JS were supported by Seed Funding from the Interacting Minds Centre, Aarhus University (2019-128). The Center for Music in the Brain is funded by the Danish National Research Foundation (DNRF 117).

**Pre-Registration**

#### I. Participants

- **Stopping Criteria: When I have tested the specific number of participants listed below, accounting for exclusions prior to looking at the data.**
- **Sample Size Commitment: Full block of 16, and we will stop after we reach 16.**
- **Compensation: Financial compensation (225 DKK).**
- **Normal Hearing:** Relied on self-report only.

#### II. Experiment Set-Up

- **Testing Language: English.**
- **Experiment Room: Room with sound-proofing materials (e.g., sound attenuating walls, curtains, etc.).**
- **Location of Training Experimenter During Experiment: In the same room/booth as the participant, and they will remain there throughout the experiment to ensure participant compliance with instructions.**
- **Location of Data Experimenter During Experiment: In a separate control room or outside the testing booth for the entire experiment.**
- **Experiment Script to be Used:** Presentation.

#### III. Audio Delivery During Experiment

- **Training Experimenter: Headphones (Beyerdynamics DT770 PRO; experimenter removed these during the test trials).**
- **Participant:** Ear inserts (Etymotic ER2; dB were not measured).

#### IV. Changes in Pre-registration Due to COVID Pandemic

- **Data collection completed:** June 12th, 2019.

**The lab’s pre-registration was followed in all other respects.**

### Lab 9: mindlab-neu

**Contributing Authors:** Psyche Loui^1^, Parker Tichko^1^

^1^ Department of Music, Northeastern University, Boston, MA, United States

**Funding Acknowledgements:** NIH R21AH075232, NIH R01AG078376, NSF-CAREER 1945436, NSF-BCS 2240330

**Pre-Registration**

#### I. Participants

- **Stopping Criteria: When I have tested the specific number of participants listed below, accounting for exclusions prior to looking at the data.**
- **Sample Size Commitment: Full block of 16, and we will stop after we reach 16.**
- **Compensation: Financial compensation ($15/hour).**
- **Normal Hearing:** Relied on self-report only.

#### II. Experiment Set-Up

- **Testing Language: English.**
- **Experiment Room: Sound -“proof” booth (single walled).**
- **Location of Training Experimenter During Experiment: In the same room/booth as the participant, and they will remain there throughout the experiment to ensure participant compliance with instructions.**
- **Location of Data Experimenter During Experiment: In a separate control room or outside the testing booth for the entire experiment.**
- **Experiment Script to be Used:** Psychopy.

#### III. Audio Delivery During Experiment

- **Training Experimenter: Headphones (Sennheiser 280; experimenter removed these during the test trials).**
- **Participant:** Headphones (Sennheiser 280; dB were not measured).

#### IV. Changes in Pre-registration Due to COVID Pandemic

- **Data collection completed:** November 2nd, 2021.

**The lab’s pre-registration was followed in all other respects.**

### Lab 10: bdl_uakl

**Contributing Authors:** Grace A. Lourie^1^, David Moreau^1^

^1^ School of Psychology and Centre for Brain Research, University of Auckland, Auckland, New Zealand

**Funding Acknowledgements:** None

**Pre-Registration**

#### I. Participants

- **Stopping Criteria:** When I have tested the specific number of participants listed below, accounting for exclusions prior to looking at the data.
- **Sample Size Commitment:** Half block of 8, but will continue collecting until the end date for data collection.
- Compensation:
- **Normal Hearing:** Conducted hearing tests (KUDUwave Software or equivalent).

#### II. Experiment Set-Up

- **Testing Language: English**
- **Experiment Room: Sound -“proof” booth (single walled).**
- **Location of Training Experimenter During Experiment: In the same room/booth as the participant, and they will remain there throughout the experiment to ensure participant compliance with instructions.**
- **Location of Data Experimenter During Experiment: In a separate control room or outside the testing booth for the entire experiment.**
- **Experiment Script to be Used:** Psychtoolbox.

#### III. Audio Delivery During Experiment

- **Training Experimenter: Headphones (Etimotic ER3; experimenter removed these during the test trials).**
- **Participant:** Headphones (Etimotic ER3; dB were measured; 23 dB in silence, 26 dB to sound (headphones in contact with decibel meter).

#### IV. Changes in Pre-registration Due to COVID Pandemic

- **Data collection completed: March 5th, 2020.**
- **Compensation:** Participants were compensated with either a voucher ($10/hour) or course credits (0.5 units/half hour), if they were recruited from UofA Experiential Learning Components (PSYCH 204 and PSYCH 207).

**The lab’s pre-registration was followed in all other respects.**

### Lab 11: smartlab-ryerson

**Contributing Authors:** Sean A. Gilmore^1^, Frank A. Russo^1^

^1^ Department of Psychology, Toronto Metropolitan University, Toronto, ON, Canada

**Funding Acknowledgements:** Natural Science and Engineering Research Council (2017-06969)

**Pre-Registration**

#### I. Participants

- **Stopping Criteria: When I have tested the specific number of participants listed below, accounting for exclusions prior to looking at the data.**
- **Sample Size Commitment: Full block of 16, and will stop after we reach 16.**
- **Compensation: Course credit (2 ½ credits).**
- **Normal Hearing:** Relied on self-report only.

#### II. Experiment Set-Up

- **Testing Language: English.**
- **Experiment Room: Sound-“proof” booth (single walled).**
- **Location of Training Experimenter During Experiment: In the same room/booth as the participant, and they will remain there throughout the experiment to ensure participant compliance with instructions.**
- **Location of Data Experimenter During Experiment: In a separate control room or outside the testing booth for the entire experiment.**
- **Experiment Script to be Used:** Presentation.

#### III. Audio Delivery During Experiment

- **Training Experimenter: Ear inserts (3M E-A-RTONE Gold 3A; experimenter removed these during test trials).**
- **Participant:** Ear inserts (3M E-A-RTONE Gold 3A; dB were not measured).

#### IV. Changes in Pre-registration Due to COVID Pandemic

- **Data collection completed:** July 5th, 2022.

**The lab’s pre-registration was followed in all other respects.**

### Lab 12: unlv-acnl

**Authors:** Karli M. Nave^1,2^, Joel S. Snyder^1^, Erin E. Hannon^1^

¹ Department of Psychology, University of Nevada Las Vegas, Las Vegas, NV, United States

² Department of Psychology, Centre for Brain and Mind, University of Western Ontario, London, ON, Canada

**Funding Acknowledgements:** UNLV Barrick Fellowship (2018-2019, to KMN), UNLV Summer Doctoral Fellowship (2019 & 2020, to KMN), UNLV Top Tier Doctoral Graduate Research Assistantship (2019-2021, to EEH)

**Pre-registration**

#### I. Participants

- **Stopping Criteria:** Data collection deadline (will continue testing as many participants as possible before that date).
- **Sample Size Commitment:** Full block of 16, then will continue collecting until the end date for data collection.
- **Compensation:** Course credit (1 credit per hour).
- **Normal Hearing:** Verified via a hearing (audiometer) test.

#### II. Experiment Set-Up

- **Testing Language:** English.
- **Experiment Room:** Sound-attenuated booth (single-walled).
- **Location of Training Experimenter During Experiment:** In the same room/booth as the participant during the entire study.
- **Location of Data Experimenter During Experiment:** In a separate area/room outside of the testing booth during the entire study.
- **Experiment Script to be Used:** Presentation script* (written by K. Nave).

*Default script provided for use by participating labs.

#### III. Audio Delivery During Experiment

- **Training Experimenter:** Sennheiser 280 Pro headphones (experimenter removed these during test trials).
- **Participant:** Ear inserts (Etymotic ER-3; dB levels measured: 20 dB in silence, 60 dB to sound).

#### IV. Changes in Pre-registration Due to COVID Pandemic

- **Data collection completed:** September 20th, 2021.

**The lab’s pre-registration was followed in all other respects.**

### Lab 13: UoD-keitellab

**Contributing Authors:** Sarah C. Allen¹, Anne Keitel¹

^1^ Department of Psychology, University of Dundee, Dundee, UK

**Funding acknowledgements:** Medical Research Council (Grant Number MR/W02912X/1; RSE Saltire Facilitation Network Award (Reference Number 1963)

**Pre-Registration**

#### I. Participants

- **Stopping Criteria: When I have tested the specific number of participants listed below, accounting for exclusions prior to looking at the data**
- **Sample Size Commitment: Full block of 16, and will stop after we reach 16**
- **Compensation: Financial compensation (Â£15)**
- **Normal Hearing:** Relied on self-report only

#### II. Experiment Set-Up

- **Testing Language: English**
- **Experiment Room: Room with occasional mild/moderate noise from adjacent room(s)**
- **Location of Training Experimenter During Experiment: In the same room/booth as the participant, and they will remain there throughout the experiment to ensure participant compliance with instructions**
- **Location of Data Experimenter During Experiment: In a separate control room or outside the testing booth for the entire experiment**
- **Experiment Script to be Used:** Psychtoolbox

#### III. Audio Delivery During Experiment

- **Training Experimenter: Headphones (Diablo Draco HS880; experimenter removed these during the test trials)**
- **Participant:** Headphones (Sennheiser HD 25; dB were measured; ∼39 dB in silence, ∼46 dB to sound)

#### IV. Changes in Pre-registration Due to COVID Pandemic

- **Data collection completed:** December 2nd, 2021
- Location of Training Experimenter during Experiment: The experimenter left the room during the experiment phase
- **The lab’s pre-registration was followed in all other respects.**

### Lab 14: vumc-mtsu

**Contributing Authors:** Reyna L. Gordon^1^, Eniko Ladanyi^1,2^, Cyrille L. Magne^3^, Srishti Nayak^1,2^

^1^ Department of Otolaryngology - Head and Neck Surgery, Vanderbilt University Medical Center, Nashville, TN, United States

^2^ Department of Linguistics, University of Potsdam, Potsdam, Germany

^3^ Department of Psychology, Middle Tennessee State University, Murfreesboro, TN, United States

**Funding acknowledgements:** NIH DP2HD098859, NSF 1926794, NSF 1926736

**Pre-Registration**

#### I. Participants

- **Stopping Criteria: When I have tested the specific number of participants listed below, accounting for exclusions prior to looking at the data**
- **Sample Size Commitment: Half block of 8, and will stop after we reach 8**
- **Compensation: Financial compensation ($20)**
- **Normal Hearing:** Used an audiometer to conduct hearing tests

#### II. Experiment Set-Up

- **Testing Language: English**
- **Experiment Room: Sound-“proof” booth (double walled)**
- **Location of Training Experimenter During Experiment: The experimenter will leave the booth during the testing phase, and the participant will be monitored through a camera from outside**
- **Location of Data Experimenter During Experiment: In a separate control room or outside the testing booth for the entire experiment**
- **Experiment Script to be Used:** E-Prime 2.0

#### III. Audio Delivery During Experiment

- **Training Experimenter: Headphones (Sony WH-1000X MH; experimenter removed these during the test trials)**
- **Participant:** Headphones (Bose QuietComfort 35; dB was not measured)

#### IV. Changes in Pre-registration Due to COVID Pandemic

- **Data collection completed:** April 26th, 2022

**The lab’s pre-registration was followed in all other respects.**

## Footnotes

1 In the original study (Nozaradan et al., 2011), beat perception is defined as “the ability to perceive periodicities from sounds that are not necessarily periodic in reality,” similar to our definition here. However, the original study also referred to 2.4 Hz as the ‘beat frequency,’ even though 2.4 Hz is in fact the stimulus periodicity, and thus a response at that frequency reflects the physical stimulus and does not necessarily entail any perceptual inference. By contrast, the imagery task requires listeners to infer a single beat level that occurs every two (or three) stimulus events. We argue that meter perception would entail inferring multiple hierarchical beat levels simultaneously (for example the ternary or binary beat pattern plus the downbeat of the measure). However, in this task listeners are only required to infer one beat level. For this reason, in this paper we refer to 2.4 Hz as the stimulus-related frequency and 0.8Hz/1.2Hz as the beat- or imagery-related frequencies.

2 Note the original study reported a One-Way ANOVA as their primary analysis, followed up by post-hoc testing. While this RR reports these results from each individual lab in a table, the meta-analyses were conducted on raw mean differences of the SNR-corrected amplitude of electric activity at the imagery-related frequencies among the three conditions.

3 Note that labs could choose to contribute a half sample of eight participants, if necessary. Some labs who originally committed to a full sample (*n* = 16) changed to a half sample (*n* = 8) after the COVID-19 Pandemic (see Appendix A for individual lab details)

4 In the case that a lab had approval from their ethics board to run emancipated minors under the age of 18, this was allowed.

5 Prior to conducting the RR study, the parent lab conducted a pilot behavioral experiment to test the training instructions and listeners’ performance on the two behavioral measures. Sixteen adult listeners with normal hearing participated in a pilot study. All participants passed training and completed 10 test trials for both binary imagery and ternary imagery conditions. Participant accuracy on the probe task (M = 0.67, SD = 0.18) was significantly above chance performance (t(15) = 3.709, p = .002), with no effect of imagery type or probe type and no interactions (p > .05). Their imagery success ratings (M = 5.24, SD = 1.37) were significantly above the scale midpoint (t(15) = 3.608, p = .003), with no effect of imagery type or probe type and no interactions (p > .05). While probe task performance and imagery success ratings demonstrated a positive correlation, it was not statistically significant (r = 0.41, p = .119).

6 Lab 15 was excluded prior to data analysis due to errors in their trial event code timings that could not be resolved by the first author nor the lab authors, at which point they requested their data be excluded.

7 Of the 16 participants included for Lab 4, five had their EEG data collected with a sampling frequency of 500Hz. These participants were processed with the steps that follow, except note the following differences: frequency resolution = 0.06 Hz, frequency spectrum: 0 to 250 Hz, bins for subtraction ranged from ±0.18 to ±0.30 Hz. When sampling frequency (500 Hz, 1000 Hz) was entered as a between-participants variable in Lab 4’s One-Way ANOVAs (section 5.3), it resulted in no significant interactions with or main effects of sampling frequency. Thus, data were collapsed across sampling frequency for all subsequent analyses.

8 Other lab-specific information is provided in Table S1 and thus is not listed in this appendix. This includes: lab location, experiment script version, audio presentation method, EEG system, number of scalp electrodes, external (additional) electrode positions, sampling frequency (SF), low pass (LP) filter, number participants collected, number participants included (after exclusions), and exclusion reasons.

## References

Bouwer, F. L., Fahrenfort, J. J., Millard, S. K., Kloosterman, N. A., & Slagter, H. A. (2023). A Silent Disco: Differential Effects of Beat-based and Pattern-based Temporal Expectations on Persistent Entrainment of Low-frequency Neural Oscillations. Journal of Cognitive Neuroscience, 35(6), 990–1020. 10.1162/jocn_a_01985

Brysbaert, M. (2019). How Many Participants Do We Have to Include in Properly Powered Experiments? A Tutorial of Power Analysis with Reference Tables. Journal of Cognition, 2(1), 16. 10.5334/joc.72

Cameron, D. J., Zioga, I., Lindsen, J. P., Pearce, M. T., Wiggins, G. A., Potter, K., & Bhattacharya, J. (2019). Neural entrainment is associated with subjective groove and complexity for performed but not mechanical musical rhythms. Experimental Brain Research, 237(8), 1981–1991. 10.1007/s00221-019-05557-4

Celma-Miralles, A., de Menezes, R. F., & Toro, J. M. (2016). Look at the Beat, Feel the Meter: Top–Down Effects of Meter Induction on Auditory and Visual Modalities. Frontiers in Human Neuroscience, 10. 10.3389/fnhum.2016.00108

Celma-Miralles, A., & Toro, J. M. (2019). Ternary meter from spatial sounds: Differences in neural entrainment between musicians and non-musicians. Brain and Cognition, 136, 103594. 10.1016/j.bandc.2019.103594

Chemin, B., Mouraux, A., & Nozaradan, S. (2014). Body Movement Selectively Shapes the Neural Representation of Musical Rhythms. Psychological Science, 25(12), 2147–2159. 10.1177/0956797614551161

Cheng, T.-H. Z., Creel, S. C., & Iversen, J. R. (2021). How do you feel the rhythm: Dynamic motor-auditory interactions are involved in the imagination of hierarchical timing. Journal of Neuroscience. 10.1523/JNEUROSCI.1121-21.2021

Cirelli, L. K., Spinelli, C., Nozaradan, S., & Trainor, L. J. (2016). Measuring Neural Entrainment to Beat and Meter in Infants: Effects of Music Background. Frontiers in Neuroscience, 10. 10.3389/fnins.2016.00229

David, S. V., Mesgarani, N., & Shamma, S. A. (2007). Estimating sparse spectro-temporal receptive fields with natural stimuli. Network: Computation in Neural Systems, 18(3), 191–212. 10.1080/09548980701609235

de Cheveigné, A., Wong, D. D. E., Di Liberto, G. M., Hjortkjær, J., Slaney, M., & Lalor, E. (2018). Decoding the auditory brain with canonical component analysis. NeuroImage, 172, 206–216. 10.1016/j.neuroimage.2018.01.033

Dehaene-Lambertz, G., Pallier, C., Serniclaes, W., Sprenger-Charolles, L., Jobert, A., & Dehaene, S. (2005). Neural correlates of switching from auditory to speech perception. NeuroImage, 24(1), 21–33. 10.1016/j.neuroimage.2004.09.039

Delorme, A., & Makeig, S. (2004). EEGLAB: An open source toolbox for analysis of single-trial EEG dynamics including independent component analysis. Journal of Neuroscience Methods, 134(1), 9–21. 10.1016/j.jneumeth.2003.10.009

Ding, N., Melloni, L., Zhang, H., Tian, X., & Poeppel, D. (2016). Cortical tracking of hierarchical linguistic structures in connected speech. Nature Neuroscience, 19(1), 158–164. 10.1038/nn.4186

Edalati, M., Wallois, F., Safaie, J., Ghostine, G., Kongolo, G., Trainor, L. J., & Moghimi, S. (2023). Rhythm in the Premature Neonate Brain: Very Early Processing of Auditory Beat and Meter. Journal of Neuroscience, 43(15), 2794–2802. 10.1523/JNEUROSCI.1100-22.2023

Elhilali, M., Xiang, J., Shamma, S. A., & Simon, J. Z. (2009). Interaction between Attention and Bottom-Up Saliency Mediates the Representation of Foreground and Background in an Auditory Scene. PLOS Biology, 7(6), e1000129. 10.1371/journal.pbio.1000129

Flaten, E., Marshall, S. A., Dittrich, A., & Trainor, L. J. (2022). Evidence for top-down metre perception in infancy as shown by primed neural responses to an ambiguous rhythm. European Journal of Neuroscience, 55(8), 2003–2023. 10.1111/ejn.15671

Frigo, M., & Johnson, S. G. (1998). FFTW: An adaptive software architecture for the FFT. Proceedings of the 1998 IEEE International Conference on Acoustics, Speech and Signal Processing, ICASSP ‘98 (Cat. No.98CH36181), 3, 1381–1384 vol.3. 10.1109/ICASSP.1998.681704

Fujioka, T., Trainor, L. J., Large, E. W., & Ross, B. (2009). Beta and Gamma Rhythms in Human Auditory Cortex during Musical Beat Processing. Annals of the New York Academy of Sciences, 1169(1), 89–92. 10.1111/j.1749-6632.2009.04779.x

Fujioka, T., Trainor, L. J., Large, E. W., & Ross, B. (2012). Internalized Timing of Isochronous Sounds Is Represented in Neuromagnetic Beta Oscillations. Journal of Neuroscience, 32(5), 1791–1802. 10.1523/JNEUROSCI.4107-11.2012

Gibbings, A., Henry, M. J., Cruse, D., Stojanoski, B., & Grahn, J. A. (2023). Attention modulates neural measures associated with beat perception. European Journal of Neuroscience, 57(9), 1529–1545. 10.1111/ejn.15962

Gilmore, S. A., & Russo, F. A. (2021). Neural and Behavioral Evidence for Vibrotactile Beat Perception and Bimodal Enhancement. Journal of Cognitive Neuroscience, 33(4), 635–650. 10.1162/jocn_a_01673

Halpern, A. R., & Zatorre, R. J. (1999). When That Tune Runs Through Your Head: A PET Investigation of Auditory Imagery for Familiar Melodies. Cerebral Cortex, 9(7), 697–704. 10.1093/cercor/9.7.697

Harding, E. E., Sammler, D., Henry, M. J., Large, E. W., & Kotz, S. A. (2019). Cortical tracking of rhythm in music and speech. NeuroImage, 185, 96–101. 10.1016/j.neuroimage.2018.10.037

Iversen, J. R., & Patel, A. D. (2008). The Beat Alignment Test (BAT): Surveying beat processing abilities in the general population. Proceedings of the 10th International Conference on Music Perception and Cognition. https://www.researchgate.net/profile/John-Iversen-2/publication/228483453_The_Beat_Alignment_Test_BAT_Surveying_beat_processing_abilities_in_the_general_population/links/00b7d5233b33d2bd39000000/The-Beat-Alignment-Test-BAT-Surveying-beat-processing-abilities-in-the-general-population.pdf

Iversen, J. R., Repp, B. H., & Patel, A. D. (2009). Top-Down Control of Rhythm Perception Modulates Early Auditory Responses. Annals of the New York Academy of Sciences, 1169(1), 58–73. 10.1111/j.1749-6632.2009.04579.x

Jones, M. R., & Boltz, M. (1989). Dynamic attending and responses to time. Psychological Review, 96(3), 459.

Jongsma, M. L., Desain, P., & Honing, H. (2004). Rhythmic context influences the auditory evoked potentials of musicians and nonmusicians. Biological Psychology, 66(2), 129–152.

Jung, T.-P., Makeig, S., Humphries, C., Lee, T.-W., Mckeown, M. J., Iragui, V., & Sejnowski, T. J. (2000). Removing electroencephalographic artifacts by blind source separation. Psychophysiology, 37(2), 163–178.

Kim, C.-Y., & Blake, R. (2005). Psychophysical magic: Rendering the visible ‘invisible.’ Trends in Cognitive Sciences, 9(8), 381–388. 10.1016/j.tics.2005.06.012

Kosslyn, S. M., Alpert, N. M., Thompson, W. L., Maljkovic, V., Weise, S. B., Chabris, C. F., Hamilton, S. E., Rauch, S. L., & Buonanno, F. S. (1993). Visual Mental Imagery Activates Topographically Organized Visual Cortex: PET Investigations. Journal of Cognitive Neuroscience, 5(3), 263–287. 10.1162/jocn.1993.5.3.263

Kosslyn, S. M., Ball, T. M., & Reiser, B. J. (1978). Visual images preserve metric spatial information: Evidence from studies of image scanning. Journal of Experimental Psychology: Human Perception and Performance, 4(1), 47–60. 10.1037/0096-1523.4.1.47

Laffere, A., Dick, F., Holt, L. L., & Tierney, A. (2021). Attentional modulation of neural entrainment to sound streams in children with and without ADHD. NeuroImage, 224, 117396. 10.1016/j.neuroimage.2020.117396

Large, E. W., & Jones, M. R. (1999). The dynamics of attending: How people track time-varying events. Psychological Review, 106(1), 119.

Large, E. W., & Palmer, C. (2002). Perceiving temporal regularity in music. Cognitive Science, 26(1), 1–37.

Lenc, T., Keller, P. E., Varlet, M., & Nozaradan, S. (2018). Neural tracking of the musical beat is enhanced by low-frequency sounds. Proceedings of the National Academy of Sciences, 115(32), 8221–8226. 10.1073/pnas.1801421115

Lenc, T., Peter, V., Hooper, C., Keller, P. E., Burnham, D., & Nozaradan, S. (2023). Infants show enhanced neural responses to musical meter frequencies beyond low-level features. Developmental Science, 26(5), e13353. 10.1111/desc.13353

Makeig, S., Bell, A. J., Jung, T.-P., & Sejnowski, T. J. (1996). Independent Component Analysis of Electroencephalographic Data. In D. S. Touretzky, M. C. Mozer, & M. E. Hasselmo (Eds.), Advances in Neural Information Processing Systems 8 (pp. 145–151). MIT Press. http://papers.nips.cc/paper/1091-independent-component-analysis-of-electroencephalographic-data.pdf

Masicampo, E. J., & Lalande, D. R. (2012). A peculiar prevalence of p values just below .05. Quarterly Journal of Experimental Psychology (2006), 65(11), 2271–2279. 10.1080/17470218.2012.711335

Mathias, B., Zamm, A., Gianferrara, P. G., Ross, B., & Palmer, C. (2020). Rhythm Complexity Modulates Behavioral and Neural Dynamics During Auditory–Motor Synchronization. Journal of Cognitive Neuroscience, 32(10), 1864–1880. 10.1162/jocn_a_01601

McGrath, S., Zhao, X., Ozturk, O., Katzenschlager, S., Steele, R., & Benedetti, A. (2024). metamedian: An R package for meta-analyzing studies reporting medians. Research Synthesis Methods, 15(2), 332–346. 10.1002/jrsm.1686

Müllensiefen, D., Gingras, B., Musil, J., & Stewart, L. (2014). Measuring the facets of musicality: The Goldsmiths Musical Sophistication Index (Gold-MSI). Personality and Individual Differences, 60, S35. 10.1016/j.paid.2013.07.081

Nave, K. (2026). Replication of EEG Frequency Tagging DATA [Dataset]. Harvard Dataverse. 10.7910/DVN/ELZFWK

Nave, K. M., Hannon, E. E., & Snyder, J. S. (2022). Steady state-evoked potentials of subjective beat perception in musical rhythms. Psychophysiology, 59(2), e13963. 10.1111/psyp.13963

Nguyen, T., Reisner, S., Lueger, A., Wass, S. V., Hoehl, S., & Markova, G. (2023). Sing to me, baby: Infants show neural tracking and rhythmic movements to live and dynamic maternal singing. Developmental Cognitive Neuroscience, 64, 101313. 10.1016/j.dcn.2023.101313

Nobre, A. C., & van Ede, F. (2018). Anticipated moments: Temporal structure in attention. Nature Reviews. Neuroscience, 19(1), 34–48. 10.1038/nrn.2017.141

Nozaradan, S., Peretz, I., & Keller, P. E. (2016). Individual Differences in Rhythmic Cortical Entrainment Correlate with Predictive Behavior in Sensorimotor Synchronization. Scientific Reports, 6, 20612. 10.1038/srep20612

Nozaradan, S., Peretz, I., Missal, M., & Mouraux, A. (2011). Tagging the neuronal entrainment to beat and meter. The Journal of Neuroscience: The Official Journal of the Society for Neuroscience, 31(28), 10234–10240. 10.1523/JNEUROSCI.0411-11.2011

Nozaradan, S., Peretz, I., & Mouraux, A. (2012). Selective Neuronal Entrainment to the Beat and Meter Embedded in a Musical Rhythm. Journal of Neuroscience, 32(49), 17572–17581. 10.1523/JNEUROSCI.3203-12.2012

Nozaradan, S., Schönwiesner, M., Keller, P. E., Lenc, T., & Lehmann, A. (2018). Neural bases of rhythmic entrainment in humans: Critical transformation between cortical and lower-level representations of auditory rhythm. European Journal of Neuroscience, 47(4), 321–332. 10.1111/ejn.13826

Nozaradan, S., Schwartze, M., Obermeier, C., & Kotz, S. A. (2017). Specific contributions of basal ganglia and cerebellum to the neural tracking of rhythm. Cortex, 95, 156–168. 10.1016/j.cortex.2017.08.015

Oostenveld, R., Fries, P., Maris, E., & Schoffelen, J.-M. (2010). FieldTrip: Open Source Software for Advanced Analysis of MEG, EEG, and Invasive Electrophysiological Data. Computational Intelligence and Neuroscience, 2011, e156869. 10.1155/2011/156869

Open Science Collaboration. (2015). Estimating the reproducibility of psychological science. Science, 349(6251), aac4716. 10.1126/science.aac4716

Palmer, C., & Krumhansl, C. L. (1990). Mental representations for musical meter. Journal of Experimental Psychology: Human Perception and Performance, 16(4), 728.

Parbery-Clark, A., Skoe, E., & Kraus, N. (2009). Musical Experience Limits the Degradative Effects of Background Noise on the Neural Processing of Sound. Journal of Neuroscience, 29(45), 14100–14107. 10.1523/JNEUROSCI.3256-09.2009

Poikonen, H., Toiviainen, P., & Tervaniemi, M. (2016). Early auditory processing in musicians and dancers during a contemporary dance piece. Scientific Reports, 6, 33056.

Pressnitzer, D., & Hupé, J.-M. (2006). Temporal Dynamics of Auditory and Visual Bistability Reveal Common Principles of Perceptual Organization. Current Biology, 16(13), 1351–1357. 10.1016/j.cub.2006.05.054

Pylyshyn, Z. (1981). The imagery debate: Analogue media versus tacit knowledge. Psychological Review, 88(1), 16–45. 10.1037/0033-295X.88.1.16

Pylyshyn, Z. (2003). Return of the mental image: Are there really pictures in the brain? Trends in Cognitive Sciences, 7(3), 113–118. 10.1016/S1364-6613(03)00003-2

Rajendran, V. G., Harper, N. S., Garcia-Lazaro, J. A., Lesica, N. A., & Schnupp, J. W. (2017). Midbrain adaptation may set the stage for the perception of musical beat. Proceedings of the Royal Society B: Biological Sciences, 284(1866), 20171455.

Remez, R. E., Rubin, P. E., Pisoni, D. B., & Carrell, T. D. (1981). Speech perception without traditional speech cues. Science, 212(4497), 947–949. 10.1126/science.7233191

Rosen, A. C., Rao, S. M., Caffarra, P., Scaglioni, A., Bobholz, J. A., Woodley, S. J., Hammeke, T. A., Cunningham, J. M., Prieto, T. E., & Binder, J. R. (1999). Neural Basis of Endogenous and Exogenous Spatial Orienting: A Functional MRI Study. Journal of Cognitive Neuroscience, 11(2), 135–152. 10.1162/089892999563283

Saupe, K., Widmann, A., Bendixen, A., Müller, M. M., & Schröger, E. (2009). Effects of intermodal attention on the auditory steady-state response and the event-related potential. Psychophysiology, 46(2), 321–327. 10.1111/j.1469-8986.2008.00765.x

Schmalz, X., Biurrun Manresa, J., & Zhang, L. (2023). What is a Bayes factor? Psychological Methods, 28(3), 705–718. 10.1037/met0000421

Silberstein, R. B. (1995). Steady-state visually evoked potentials, brain resonance, and cognitive processes. Neocortical Dynamics and EEG Rhythms, 272–303.

Snyder, J. S., & Large, E. W. (2005). Gamma-band activity reflects the metric structure of rhythmic tone sequences. Brain Research. Cognitive Brain Research, 24(1), 117–126. 10.1016/j.cogbrainres.2004.12.014

Stupacher, J., Wood, G., & Witte, M. (2017). Neural Entrainment to Polyrhythms: A Comparison of Musicians and Non-musicians. Frontiers in Neuroscience, 11. 10.3389/fnins.2017.00208

Tal, I., Large, E. W., Rabinovitch, E., Wei, Y., Schroeder, C. E., Poeppel, D., & Golumbic, E. Z. (2017a). Neural entrainment to the beat: The “missing-pulse” phenomenon. Journal of Neuroscience, 37(26), 6331–6341.

Tal, I., Large, E. W., Rabinovitch, E., Wei, Y., Schroeder, C. E., Poeppel, D., & Golumbic, E. Z. (2017b). Neural Entrainment to the Beat: The “Missing-Pulse” Phenomenon. Journal of Neuroscience, 37(26), 6331–6341. 10.1523/JNEUROSCI.2500-16.2017

Tichko, P., Page, N., Kim, J. C., Large, E. W., & Loui, P. (2022). Neural Entrainment to Musical Pulse in Naturalistic Music Is Preserved in Aging: Implications for Music-Based Interventions. Brain Sciences, 12(12), 1676. 10.3390/brainsci12121676

Tierney, A., & Kraus, N. (2013). The Ability to Move to a Beat Is Linked to the Consistency of Neural Responses to Sound. Journal of Neuroscience, 33(38), 14981–14988. 10.1523/JNEUROSCI.0612-13.2013

Tierney, A., & Kraus, N. (2015). Neural Entrainment to the Rhythmic Structure of Music. Journal of Cognitive Neuroscience, 27(2), 400–408. 10.1162/jocn_a_00704

Tononi, G., Srinivasan, R., Russell, D. P., & Edelman, G. M. (1998). Investigating neural correlates of conscious perception by frequency-tagged neuromagnetic responses. Proceedings of the National Academy of Sciences, 95(6), 3198–3203. 10.1073/pnas.95.6.3198

Vanden Bosch der Nederlanden, C. M., Joanisse, M. F., & Grahn, J. A. (2020). Music as a scaffold for listening to speech: Better neural phase-locking to song than speech. NeuroImage, 214, 116767. 10.1016/j.neuroimage.2020.116767

Viechtbauer, W. (2010). Conducting Meta-Analyses in R with the metafor Package. Journal of Statistical Software, 36(1), 1–48. 10.18637/jss.v036.i03

Vuust, P., & Witek, M. A. G. (2014). Rhythmic complexity and predictive coding: A novel approach to modeling rhythm and meter perception in music. Frontiers in Psychology, 5, 1111. 10.3389/fpsyg.2014.01111

Whiteford, K. L., Baltzell, L. S., Chiu, M., Cooper, J. K., Faucher, S., Goh, P. Y., Hagedorn, A., Irsik, V. C., Irvine, A., Lim, S.-J., Mesik, J., Mesquita, B., Oakes, B., Rajappa, N., Roverud, E., Schrlau, A. E., Van Hedger, S. C., Bharadwaj, H. M., Johnsrude, I. S., … Oxenham, A. J. (2025). Large-scale multi-site study shows no association between musical training and early auditory neural sound encoding. Nature Communications, 16(1), 7152. 10.1038/s41467-025-62155-5

Winkler, I., Háden, G. P., Ladinig, O., Sziller, I., & Honing, H. (2009). Newborn infants detect the beat in music. Proceedings of the National Academy of Sciences, 106(7), 2468–2471.

Zalta, A., Large, E. W., Schön, D., & Morillon, B. (2024). Neural dynamics of predictive timing and motor engagement in music listening. Science Advances, 10(10), eadi2525. 10.1126/sciadv.adi2525

Zanto, T. P., Snyder, J. S., & Large, E. W. (2006). Neural correlates of rhythmic expectancy. Advances in Cognitive Psychology, 2(2–3), 221–231. 10.2478/v10053-008-0057-5

Zatorre, R. J., Halpern, A. R., Perry, D. W., Meyer, E., & Evans, A. C. (1996). Hearing in the Mind’s Ear: A PET Investigation of Musical Imagery and Perception. Journal of Cognitive Neuroscience, 8(1), 29–46. 10.1162/jocn.1996.8.1.29

Zhao, T. C., & Kuhl, P. K. (2020). Neural and physiological relations observed in musical beat and meter processing. Brain and Behavior, 10(11), e01836. 10.1002/brb3.1836

